# Gene interfered-ferroptosis therapy of cancer

**DOI:** 10.1101/2020.04.19.048785

**Authors:** Jinliang Gao, Tao Luo, Na Lin, Jinke Wang

## Abstract

Although some effective therapies have been available for cancer, it still poses a great threat to human health and life due to its drug resistance and low response in patients. Here, we developed a novel therapy named as gene interfered-ferroptosis therapy (GIFT) by combining iron nanoparticles and cancer-specific gene interference. Using a promoter consisted of a NF-κB decoy and a minimal promoter (DMP), we knocked down the expression of two iron metabolism-related genes (FPN and Lcn2) selectively in cancer cells. At the same time, we treated cells with Fe_3_O_4_ nanoparticles. As a result, a significant ferroptosis was induced in a wide variety of cancer cells representing various hematological and solid tumors. However, the same treatment had no effect on normal cells. By using AAV and PEI-coated Fe_3_O_4_ nanoparticles as gene vectors, we found that the tumor growth in mice could be also significantly inhibited by the intravenously injected GIFT reagents. By detecting ROS, iron content and gene expression, we confirmed that the mechanism underlying the therapy is gene inference-enhanced ferroptosis.

## 1. Introduction

Although some effective therapies have been available for cancer, it still poses a great threat to human health and life due to its drug resistance and low response in patients. Therefore, various cancer therapies have been being developed to induce cancer cell death through one form of cell death such as apoptosis, necrosis, necroptosis, and pyroptosis [1]. Recently, ferroptosis has attracted increasing attention as another form of cell death. Ferroptosis was identified as an iron-dependent nonapoptotic cell death in 2012 [2]. Subsequently, it was found that ferroptosis is triggered by iron-catalyzed lipid peroxidation initiated by nonenzymatic (Fenton reactions) and enzymatic mechanisms [lipoxygenases (LOXs)] [3]. In the former case, hydroxyl radicals is directly generated by the Fenton reaction between ferrous iron (Fe^2+^) and hydrogen peroxide and initiates nonenzymatic lipid peroxidation [4, 5].

Since being found, ferroptosis has attracted considerable attention due to its potential role as a target for novel therapeutic anticancer strategies [6-8]. Because ferroptosis is classified as a regulated necrosis and is more immunogenic than apoptosis, its encouraging anticancer effect may be achieved by immunogenicity [9]. More interestingly, it was found that ferroptosis can propagate among cells in a wave-like manner, exhibiting a potent killing effect on neighboring cells [10-12]. Another attractive point of ferroptosis is its potential to overcome drug resistance such as chemoresistance [13-15]. Therefore, the anti-tumor effects of ferroptosis has been being rapidly and widely explored in variant cancers [16]. However, most of these studies focused on finding various ferroptosis inducers of compounds, such as erastin and RSL3 for inhibiting or depleting system x_c_^-^, GPX4 and CoQ10 [17]. These compounds may be challenged by the same resistance problem as the traditional cancer drugs. For example, six high-confidence genes in erastin-induced ferroptosis were identified, and IREB2 was one of them. If IREB2 was silenced, the expression of iron uptake, metabolism and storage genes, such as TFRC, ISCU, FTH1, and FTL, would decrease, and the cells exhibited resistance to erastin-induced ferroptosis [2, 18, 19].

Iron is a key factor in ferroptosis. Iron oxide nanoparticles (IONPs) can release ferrous (Fe^2+^) or ferric (Fe^3+^) in acidic lysosome in cells. The released Fe^2+^ participates in the Fenton reaction to generate toxic hydroxyl radicals (•OH), one of reactive oxygen species (ROS), to induce ferroptosis of cancer cells [11, 20, 21]. Therefore, the ferroptosis-driven nanotherapeutics for cancer treatment become increasingly attractive [22]. Moreover, it was found that nanoparticle-induced ferroptosis in cancer cells eliminates all neighboring cells in a propagating wave [10]. However, cells are sensitive to iron concentration, and a little fluctuation in concentration can cause a great response [23, 24]. Due to the importance of iron to cells, cells have evolved a set of mechanisms to maintain intracellular iron homeostasis [25]. The intracellular iron is thus under delicate regulation to sustain iron homoeostasis [26-28]. Cells are therefore capable to effectively store and export the intracellular excess iron ions released from internalized IONPs. For example, in previous study, we found significant up-regulation of transcription of important genes responsible for exporting intracellular iron ion in cells treated by a DMSA-coated Fe_3_O_4_ nanoparticle [29]. Therefore, the ferroptosis induced by a single IONPs treatment is too little to be utilized to treat cancer in clinics.

Recently, an interesting study reported a IONPs-induced ferroptosis that was enhanced by a genetic change. In the study, it was found that a nanoparticle iron supplement, ferumoxytol, has an anti-leukemia effect in vitro and in vivo in leukemia cell with a low level of FPN [26]. This anti-tumor effect resulted from the inability of the type of leukemia cell to export intracellular Fe^2+^ generated from ferumoxytol, which make the type of leukemia cell susceptible to the elevated ROS generated by Fe^2+^ through Fenton reaction. However, this study relies on the rare naturally-occurred low expression of ferroptosis-related genes such as FPN, still not resolving the problem how to artificially lower the expression of these genes in tumors of patients so that this IONPs-induced ferroptosis therapy can be applied to more wide types of cancers.

Nuclear factor-kappa B (NF-κB) is a sequence-specific DNA-binding transcription factor that is over activated in almost all cancers [30, 31]. Therefore, to inactivate the protein, we designed a gene expression vector that can express an artificial microRNA targeting NF-κB RelA/p65 under the control of a NF-κB-specific promoter [32]. This promoter consists of a NF-κB decoy sequence and a minimal promoter and thus its transcriptional activation activity is dependent on the NF-κB activity in cells [32]. We therefore name it as DMP meaning Decoy-Minimal Promoter. Our previous results verified that DMP could be used to realize cancer cell-specific gene expression by depending on the NF-κB over-activity in cancer cells [33, 34]. By using the promoter, we developed two kinds of cancer gene therapy. One is a cancer immunotherapy induced by a neoantigen displayed on cancer cell surface [33]. The other is a cancer gene therapy induced by cutting telomere via CRISPR/Cas9 [34]. In these studies, we successfully controlled neoantigen and Cas9 expression selectively in cancer cells by the DMP promoter.

Based on these previous studies, we therefore deduced that inhibition of genes exporting intracellular iron ions should be fetal for the IONPs-treated cancer cells, because their inhibition would prevent cell from exporting IONPs-produced iron ions, which can thus enhance IONPs-induced ferroptosis. Base on this speculation, we developed a new cancer therapy, gene interfered-ferroptosis therapy (GIFT) by combining the DMP-controlled gene interference tools with IONPs in this study. By specifically knocking down expression of two iron metabolism-related genes, FPN and Lcn2, with DMP-controlled CRISPR/Cas13a and miRNA in cancer cells, the treatment of Fe_3_O_4_ nanoparticles induced dramatic ferroptosis in a wide variety of cancer cells that represent various hematological and solid tumors. However, the therapy has no effect on normal cells. By using both viral (adenovirus associated virus, AAV) and non-viral (PEI-modified Fe_3_O_4_ nanoparticles) vectors, the growth of xenografted tumors in mice were also significantly inhibited by the therapy.

## 2. Material and methods

### 2.1. Vector construction

The decoy minimal promoter (DMP), a chemically synthesized NF-κB-specific promoter which contains a NF-κB response sequence and a minimal promoter sequence, was cloned into pMD19-T simple (TAKARA) to obtain pMD19-T-DMP firstly, then the human codon optimized Cas13a coding sequence amplified from the pC013-Twinstrep-SUMO-huLwCas13a (Addgene) by PCR was cloned into pMD19-T-DMP to obtain pMD19-T-DMP-Cas13a,next the chemically synthesized U6 promoter sequence and the direct repeat sequence of guide RNA of Cas13a separated by BbsI restriction sites were cloned into pMD19-T-DMP-Cas13a, respectively, to generate pDMP-Cas13a/U6-gRNA backbone (referred to as pDCUg).

The guide RNAs (gRNAs) targeting no transcript (NT), human or murine ferroportin (FPN) and Lipocalin 2 (Lcn2) were designed by CHOPCHOP (http://chopchop.cbu.uib.no/). The complementary oligonucleotides containing a 28-bp gRNA target-specific region and two flanking BbsI sites were chemically synthesized and annealed into double-stranded oligonucleotides, and then ligated into pDCUg. The ligation reaction (10 μL) consisted of 10 units of BbsI (NEB), 600 units of T4 DNA ligase (NEB), 1× T4 DNA ligase buffer, 1nM double-stranded oligonucleotides, and 50 ng plasmid pDCUg. The ligation reaction was run on a PCR cycler as follows: 10 cycles of 37°C 5 min and 16°C 10 min, 37°C 30 min, and 80°C 5 min. The generated plasmids were named as pDCUg-NT, pDCUg-hFPN/pDCUg-mFPN, and pDCUg-hLcn2/pDCUg-mLcn2, respectively. Due to the difference between human and mouse gene, pDCUg expression vectors respectively targeted to the human FPN (hFPN) and Lcn2 (hLcn2) genes and mouse FPN (hFPN) and Lcn2 (mLcn2) genes were constructed. A plasmid co-expressing gRNAs targeting ferroportin and Lipocalin 2 simultaneously, named as pDCUg-hFL/pDCUg-mFL, was also constructed. All gRNAs targeting genes of interest were listed in Supplementary Table S1.

The universal miRNA expression vector pDMP-miR was constructed based on pCMV-miR which was previously kept by our laboratory by replacing the CMV promoter with DMP promoter. The miRNAs targeting human or murine FPN and Lcn2 were designed by BLOCK-iT™ RNAi Designer (https://rnaidesigner.thermofisher.com/rnaiexpress/) and synthesized by Sangon Biotech. The synthesized oligos were denatured and then annealed to obtain double-stranded DNA (dsDNA) which were then linked with the linear pDMP-miR vector cleaved with BsmBI. The generated miRNA expression vectors targeting the FPN and Lcn2 genes were named as pDMP-miR-hFPN/pDMP-miR-mFPN (referred to as pDMhF/pDMmF) and pDMP-miR-hLcn2/pDMP-miR-mLcn2 (referred to as pDMhL/pDMmL), respectively. Vectors were detected with PCR amplification and verified by DNA sequencing. A plasmid simultaneously expressing miRNAs targeting FPN and Lcn2, pDMP-miR-hFPN-DMP-miR-hLcn2/pDMP-miR-mFPN-DMP-miR-mLcn2 (referred to as pDMhFL/pDMmFL), was also constructed. The miR-Neg fragment was synthesized according to the sequence of plasmid pcDNA™ 6.2-GW/EmGFP-miR-Neg and ligated into pDMP-miR, named pDMP-miR-Neg (referred to as pDMNeg) as a negative control vector. All the miRNAs targeting genes of interest were listed in Supplementary Table S2.

The DCUg-NT/hFL/mFL and DMNeg/DMhFL/DMmFL sequences were amplified by PCR from pAAV-DCUg-NT/hFL/mFL and pAAV-DMNeg/DMhFL/ DMmFL, respectively. By using the MluI (upstream) and XbaI (downstream) restriction sites, the PCR fragments were cloned into pAAV-MCS (VPK-410, Stratagene) to construct the pAAV-DCUg-NT/hFL/mFL and pAAV-DMNeg/DMhFL/DMmFL vectors, respectively.

### 2.2. Nanoparticles, cells and culture

The DMSA-coated Fe_3_O_4_ magnetic nanoparticles (FeNPs) were provided by the Biological and Biomedical Nanotechnology Group of the State Key Lab of Bioelectronics, Southeast University, Nanjing, China. A magnetic transfection agent (MagTransf™) was purchased from the Nanjing nanoeast biotech co., ltd (Nanjing, China), which was referred to as FeNCs in this study. Cells used in this research included KG-1a (human acute myeloid leukemia cells), HL60 (human promyeloid acute leukemia cells) and WEHI-3 (mouse acute mononuclear leukemia cells), HEK-293T (human fetal kidney cells), HepG2 (human liver cancer cells), A549 (human lung cancer cells), HT-29 (human colon cancer cells), C-33A (human cervical cancer cells), SKOV3 (human ovarian cancer cells), PANC-1 (human pancreatic cancer cells), MDA-MB-453 (human breast cancer cells), Hepa1-6 (mouse hepatoma cells), B16F10 (mouse melanoma cells), BGC-823/MGC-803/SGC-7901 (Human gastric adenocarcinoma cells), KYSE450/KYSE510 (human esophageal carcinoma cells), HL7702 (human normal hepatocytes), and MRC5 (human embryonic fibroblasts). Three leukemia cell lines, KG-1a, HL60 and WEHI-3, were cultured in Iscove’s Modified Dulbecco’s Medium (IMEM) (Gibco). HEK-293T, HepG2, Hepa1-6, C-33A, PANC-1, MDA-MB-453, B16F10, MRC-5 cells were cultured in Dulbecco’s Modified Eagle Medium (DMEM) (Gibco). A549, HT-29, SKOV-3, BGC-823/MGC-803/SGC-7901, KYSE450/KYSE510 and HL7702 cells were cultured in Roswell Park Memorial Institute (RPMI) 1640 medium (Gibco). All media were supplemented with 10% fetal bovine serum (HyClone), 100 units/mL penicillin (Thermo Fisher Scientific), and 100 μg/mL streptomycin (Thermo Fisher Scientific). Cells were incubated at 37 °C in a humidified incubator containing 5% CO2.

### 2.3. FeNPs cytotoxicity measurement

To determine the optimal dosage of nanoparticles. The in vitro cytotoxicity of FeNPs was performed by using the CCK-8 assay. KG-1a, HL60, WEHI-3, HepG2, HL7702 and MRC-5 cells were seeded in 96-well plates at a density of 5000 cells/well, respectively. Cells were cultivated overnight and treated with various concentrations (0, 30, 50, 100, 150, 200, 250 μg/mL) of FeNPs for various times. Each treatment was performed with six groups of cells and each group had four replicates. 10 μL of Cell Counting Kit-8 (CCK-8) solution (BS350B, Biosharp) was added to each well at variant time points (0 d, 1 d, 2 d, 3 d, 4 d, and 5 d) post treatment. After incubating for another 1h at 37 °C, the optical density at 450 nm was measured using a microplate reader (BioTek).

### 2.4. Cell treatment by FeNPs-based GIFT

Cells were first transfected with plasmids using Lipofectamine 2000 (Thermo Fisher Scientific) according to the manufacturer’s instructions. Briefly, cells (1×10^5^) were seeded into 24-well plates overnight before transfection. Cells were then transfected with 500 ng of various plasmids including pDCUg-NT, pDCUg-hFPN/pDCUg-mFPN, pDCUg-hLcn2/pDCUg-mLcn2, pDCUg-hFL/pDCUg-mFL, pDMNeg, pDMhF/ pDMmF, pDMhL/pDMmL and pDMhFL/pDMmFL. The transfected cells were cultured for 24 h and then incubated with or without 50 μg/mL of FeNPs, and cells were cultured for another 72h. For HL7702 and MRC5, cells were first induced with or without TNF-α (10 ng/mL) for 1 h before FeNPs treatment. At 24 h, 48 h and 72 h post FeNPs administration, all cells were stained with acridine orange and ethidium bromide (AO&EB) following the manufacturer’s instructions. Cells were imaged under a fluorescence microscope (IX51, Olympus) to observe numbers of live and dead cells. In order to quantify cell apoptosis, cells were collected at 72 h post FeNPs administration and detected with the Annexin V-FITC Apoptosis Detection Kit (BD, USA) according to the manufacturer’s instructions. The fluorescence intensity of cells was quantified with CytoFLEX LX Flow Cytometer (Beckman).

### 2.5. ROS measurement

Cells were treated with FeNPs as previously described. Briefly, Cells were seeded at a density of 1×10^5^ cells/ml medium per well in 24-well plates for overnight growth. Cells were then transfected with 500 ng of various plasmids including pDCUg-NT, pDCUg-hFL/pDCUg-mFL, pDMNeg and pDMhFL/pDMmFL. The transfected cells were cultured for 24 h and then incubated with or without 50 μg/mL of the FeNPs, and cells were cultured for another 48h. The treated cells were stained with 2′,7′-dichlorodihydrofluorescein diacetate (DCFH-DA) using Reactive Oxygen Species Assay Kit (Beyotime) according to the manufacturer’s instructions. ROS changes indicated by fluorescence shift was analyzed on a CytoFLEX LX Flow Cytometer (Beckman).

### 2.6. Iron content measurement

Cells were treated with FeNPs as previously described. Intracellular iron was determined by a complete digest of the cell 48h post FeNPs administration. Cells were washed with PBS, collected and counted. Then the cells were precipitated by centrifugation, resuspended in 50 μL of 5 M hydrochloric acid, and incubated at 60°C for 4 h. The cells were centrifuged again and the supernatants were transferred to 96-well plates. The plates were incubated at room temperature for 10 min after that each well was added with 50 μL of freshly prepared detection reagent (0.08% K_2_S_2_O_8_, 8% KSCN and 3.6% HCl dissolved in water). The absorbance at 490 nm was measured using an absorption reader (BioTek). The iron content was determined by the obtained absorbance normalized with a standard curve generated with FeCl_3_ standard solution. The iron was reported as average iron contents per cell calculated as mean values divided by the number of cells in each sample. Each experiment was repeated in triplicate wells at least six times.

### 2.7. Western blot assay

Cells were seeded into 6-well plates at a density of 2×10^5^ cells per well and grown overnight. Cells in wells were transfected with 1000 ng of plasmids including pDCUg-NT, pDCUg-hFL, pDMNeg and pDMhFL, respectively. At 48 h post transfection, the whole-cell extracts were prepared using a phosphoprotein extraction kit (SA6034-100T, Signalway Antibody, USA) according to the manufacturer’s instructions. The protein lysates (20 μg/sample) were resolved by SDS-PAGE and the target proteins were detected with Western blot (WB) using the antibodies as follows: GAPDH Rabbit monoclonal antibody (ab181602, Abcam, UK), SLC40A1 Rabbit polyclonal antibody (ab58695, Abcam, UK), Lipocalin-2 Rabbit polyclonal antibody (ab63929, Abcam, UK). The second antibodies were IRDye® 800CW Goat anti-Rabbit IgG (Licor). The blots were detected and fluorescence intensity was quantified with the Odyssey Infrared Fluorescence Imaging System (Licor).

### 2.8. Virus preparation

HEK293T cells were seeded into 75 cm^2^ flasks at a density of 5×10^6^ cells per flask and cultivated for overnight. Cells were then co-transfected with two helper plasmids (pHelper and pAAV-RC; Stratagene) and one of the pAAV plasmids (pAAV-DCUg-NT, pAAV-DCUg-hFL/pAAV-DCUg-mFL, pAAV-DMNeg, pAAV-DMhFL/pAAV-DMmFL) using Lipofectamine 2000 according to the manufacturer’s instructions. Cells were cultured for another 72 h. Viruses were then collected and purified as previously described [35]. Titers of AAVs were determined by qPCR using the primers AAV-F/R (Table S3). Quantified viruses were aliquoted and kept at -80°C for later use. The obtained viruses were named as rAAV-DCUg-NT, rAAV-DCUg-hFL/rAAV-DCUg-mFL, rAAV-DMNeg, and rAAV-DMhFL/rAAV-DMmFL.

### 2.9. Virus evaluation

KG-1a, WEHI-3 and HL7702 cells were seeded into 24-well plates (1×10^5^ cells/well) and cultivated for 12 h. Cells were then transfected with the viruses including rAAV-DCUg-NT, rAAV-DCUg-hFL/rAAV-DCUg-mFL, rAAV-DMNeg, and rAAV-DMhFL/rAAV-DMmFL at the dose of 1×10^5^ vg per cell. The transfected cells were cultured for 24 h and then incubated with or without 50 μg/mL FeNPs. Cells were cultured for another 72 h and imaged by optical microscope (Olympus), and cell viability was evaluated using a CCK-8 assay (BS350B, Biosharp).

### 2.10. Cells treatment by FeNCs-based GIFT

In order to investigate whether a PEI-modified Fe_3_O_4_ NP can be used as Fe nano-carriers (FeNCs) for gene interference vectors, two FeNCs-based GIFT experiments were performed, in which two batches of FeNCs, named as FeNCs-1 and FeNCs-2, were used.

In the first FeNCs-based GIFT experiment, various plasmids (including pDCUg-NT, pDCUg-hFL, pDMNeg, and pDMhFL) were mixed with FeNCs-1 (1 μg DNA/μg FeNCs-1) to prepare plasmid DNA-loaded FeNCs (FeNCs@DNA), including FeNCs-1@pDCUg-NT, FeNCs-1@pDCUg-hFL, FeNCs-1@ pDMNeg and FeNCs-1@pDMhFL. Cells (1×10^5^) were seeded into 24-well plates and cultured overnight. Cells were then treated with FeNCs, FeNCs@DNA (totally 0.5 μg plasmid DNA), or plasmid DNA alone for 24 hours. Cells were further cultured with a medium with or without 50 μg/mL FeNPs for 72 h. Cells were stained with AO&EB at different time points (24 h, 48 h, and 72 h) and imaged.

In the second FeNCs-based GIFT experiment, two plasmids (pDMNeg and pDMhFL) were mixed with FeNCs-1 and FeNCs-2 (1 μg DNA/μg FeNCs) according to the manufacturer’s instructions to prepare FeNCs@DNA, including FeNCs-1@pDMhFL and FeNCs-2@pDMhFL. The prepared FeNCs@DNA was used immediately to treat cells or left for 24 h before treating cells. Cells (1×10^5^) were seeded into 24-well plates and cultured overnight. Cells were then cultured with a fresh medium contained 50 μg/mL FeNCs or FeNCs@DNA for 72 h. Cells were stained with AO&EB at different time points (24 h, 48 h, and 72 h) and imaged.

### 2.11. Animal studies

Four-week-old BALB/c female mice with an average weight of 20 g were purchased from the Changzhou Cavens Laboratory Animal Co. Ltd (China). All animal experiments in this study followed the guidelines and ethics of the Animal Care and Use Committee of Southeast University (Nanjing, China). BALB/c female mice were subcutaneously transplanted with1×10^7^ WEHI-3 cells into inner thighs. After breeding for 1 week, the tumor sizes were measured with a precision caliper. Tumor volumes were calculated using formula V = (ab^2^)/2, where a is the longest diameter and b is the shortest diameter. Three batches of animal experiments were performed.

In the first batch of animal experiment, the tumor-bearing mice were randomly divided into six treatment groups (PBS, n=6; FeNPs, n=6; rAAV-DCUg-NT, n=6; rAAV-DCUg-NT plus FeNPs, n=6; rAAV-DCUg-mFL, n=6; rAAV-DCUg-mFL plus FeNPs, n=7). Each group of mice was then intravenously injected with PBS, rAAV-DCUg-NT, rAAV-DCUg-NT, rAAV-DCUg-mFL, and rAAV-DCUg-mFL, respectively. The dosage of different viruses was always 1×1 0^10^ vg/mouse. Mice in groups of FeNPs, rAAV-DCUg-NT plus FeNPs, and rAAV-DCUg-mFL plus FeNPs were intravenously injected FeNPs (3 mg/kg) on the next day. Mice were euthanized and photographed on the seventh day post FeNPs injection, and the tumor was isolated and the tumor size was measured and calculated. Mice were dissected, and various tissues (including heart, liver, spleen, lung, kidney, and tumor tissues) were collected and cryopreserved in liquid nitrogen.

In the second batch of animal experiment, the tumor-bearing mice were randomly divided into five treatment groups (FeNPs, n=6; rAAV-DMNeg, n=6; rAAV-DMmFL, n=7; rAAV-DMNeg plus FeNPs, n=7; rAAV-DMmFL plus FeNPs, n=6). Each group of mice was then intravenously injected with FeNPs, rAAV-DMNeg, rAAV-DMmFL, rAAV-DMNeg plus FeNPs, rAAV-DMmFL plus FeNPs, respectively. The dosage of different viruses and FeNPs were 1×10^10^ vg/mouse and 3 mg/kg body weight, respectively. In order to simplify the drug administration, rAAV and FeNPs were mixed together and intravenously injected to mice one time in this batch of animal experiment. Mice were euthanized and photographed on the seventh day post FeNPs injection, and the tumor was isolated and the tumor size was measured and calculated. Mice were dissected, and various tissues were collected and cryopreserved in liquid nitrogen.

In the third batch of animal experiment, the tumor-bearing mice were randomly divided into six treatment groups (PBS, n=6; FeNCs, n=6; pAAV-DMNeg plus FeNCs, n=6; pAAV-DMmFL plus FeNCs, n=7; pAAV-DCUg-NT plus FeNCs, n=6; pAAV-DCUg-mFL plus FeNCs, n=7). Each group of mice was then intravenously injected with PBS, FeNCs, pAAV-DMNeg plus FeNCs, pAAV-DMmFL plus FeNCs, pAAV-DCUg-NT plus MT-FeNCs, pAAV-DCUg-mFL plus FeNCs, respectively. Various plasmids and FeNCs were mixed according to the manufacturer’s instruction before injection, the dosage of different plasmids and FeNCs were 2 mg/kg and 3 mg/kg, respectively. Mice were euthanized and photographed on the seventh day post FeNPs injection, and the tumor was isolated and the tumor size was measured and calculated. Mice were dissected, and various tissues were collected and cryopreserved in liquid nitrogen.

### 2.12. Quantitative PCR

Total RNA was isolated from cell lines at 48h post incubation with FeNPs or mouse tissues using TRIzol™ (Invitrogen) according to the manufacturer’s protocol. The complementary DNA (cDNA) was generated using the FastKing RT kit (TIANGEN) according to the manufacturer’s instruction. The genomic DNA (gDNA) was extracted from various tissues of mice using the TIANamp Genomic DNA Kit (TIANGEN). Amplification for the genes of interest from cDNA and gDNA was performed by qPCR using the Hieff qPCR SYBR Green Master Mix (Yeasen). Triplicate samples per treatment were evaluated on a ABI Step One Plus (Applied Biosystems). Relative mRNA transcript levels were compared to the GADPH internal reference and calculated as relative quantity (RQ) according to the following equation: RQ = 2^−ΔΔCt^. Virus DNA abundance were normalized to the GADPH internal reference and calculated according to the following equation: RQ = 2^−ΔCt^. Cas13a mRNA expression levels were showed as Ct values. All experiments were performed in triplicates and repeated a minimum of three times.

NF-κB RelA/p65 expression in cells were detected by quantitative PCR (qPCR) using the primers Human/Murine RelA-F/R and Human/Murine GAPDH-F/R. The results are normalized to GAPDH and analyzed by 2^−ΔCt^ method. All the qPCR primers were verified as being specific on the basis of melting curve analysis and were listed in Supplementary Table S3.

### 2.13. Statistical analysis

All data are presented as mean± s.e.m. and statistical analysis and graphs were performed with GraphPad Prism software (GraphPad Software). The data were statistically processed using analysis of variation and the Student’s t-test. Differences at p < 0.05 were considered statistically significant. *, p < 0.05; **, p < 0.01; ***, p < 0.001.

## 3. Results

### 3.1. Conceptualization of GIFT

Figures 1A and 1B schematically illustrate the principle of gene interfered-ferroptosis therapy (GIFT). GIFT consists of a gene interfering vector (GIV) and Fe_3_O_4_ nanoparticles (FeNPs). GIV is composed of a promoter named DMP and downstream effector gene. DMP is a NF-κB-specific promoter that consists of a NF-κB decoy and a minimal promoter. Because NF-κB is a transcription factor that is widely over-activated in cancers, the effector genes can be expressed in cancer cells by NF-κB binding to DMP. However, the effector genes cannot be expressed in normal cells due to lack of NF-κB. Therefore, DMP is a cancer cell-specific promoter. When the DMP-controlled CRISPR/Cas13a or miRNA GIVs are transfected into cancer cells, Cas13a or miRNA can be expressed. The expressed Cas13a protein can be assembled into Cas13a-gRNA complex by binding gRNA transcribed from a U6 promoter. The expressed miRNAs can associate with RISC. Both Cas13a-gRNA and miRNA-RISC complexes can target the interested mRNA to knock down the expression of target genes in cancer cells.

**Figure 1.**
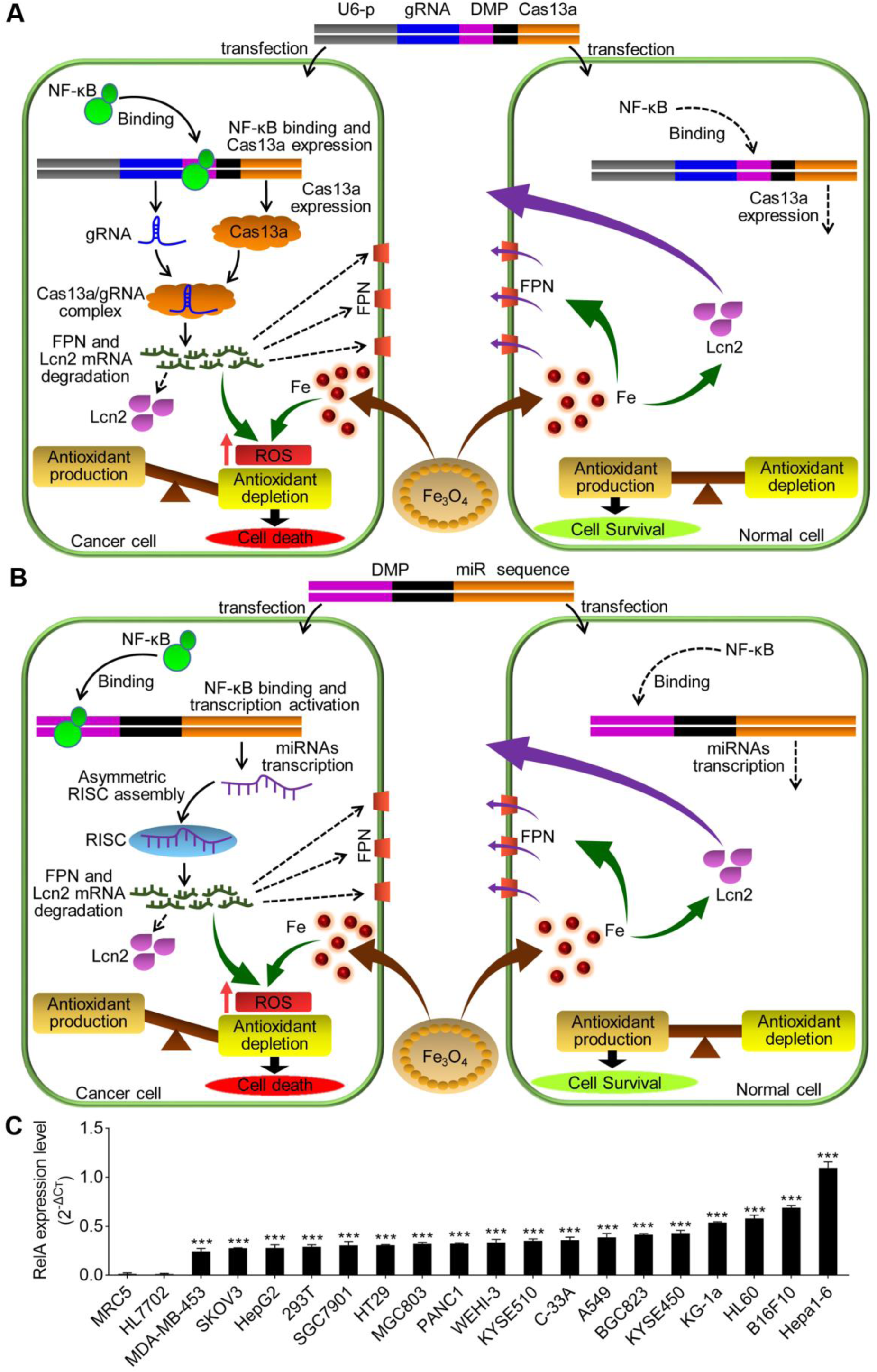
Schematic show of GIFT principle and NF-κB expression in variant cells. (A) Schematic show of CRISPR/Cas13a-based GIFT. (B) Schematic show of miRNA-based GIFT. U6-p, U6 promoter; gRNA, guide RNA; DMP, NF-κB decoy-minimal promoter; Fe_3_O_4_, FeNPs). (C) QPCR detection of NF-κB expression in variant cells. ***, p<0.001.

In this study, we selected two genes related to iron metabolism, FPN and Lcn2, as the target genes. Both the functions of FPN and Lcn2 in cells are related to the efflux of iron ions. Therefore, we speculated that knocking down the expression of these two genes in cancer cells by the DMP-controlled CRISPR/Cas13a or miRNA GIVs can prevent cells from exporting the intracellular iron ions produced by FeNPs. This can result in the accumulation of iron ions and cause a significant increase in intracellular ROS levels, thus leads to significant ferroptosis in cancer cells. In normal cells, because Cas13a or miRNA cannot be produced, the expression of these two genes cannot be affected, which allows the normal cells to actively export the intracellular iron ions produced by FeNPs and maintain the iron homeostasis, leading to no effects from FeNPs treatment.

### 3.2. Expression of NF-κB RelA in various cells

NF-κB is widely activated in nearly all types of tumor cells. However, it is expected that its expression is not the same among cancer cells [36, 37]. Because the intracellular NF-κB activity is critical to our study. Therefore, we first detected the expression of NF-κB RelA/p65 in three leukemia cell lines (KG-1a, HL60 and WEHI-3), fifteen other types of solid tumor cells, and two normal human cells (HL7702 and MRC5). The results demonstrated that NF-κB RelA/p65 expressed at different levels in all cancer cell lines, but did not in normal cells (Fig.1C). Therefore, DMP should drive cancer cell-specific gene expression.

### 3.3. Effects of FeNPs on viability of leukemia cells

To evaluate the cytotoxicity of FeNPs, we dynamically measured the cell viability of three leukemia cells, a solid tumor cell (HepG2) and two normal human cells that were treated by various concentrations of FeNPs for 5 days. The results showed FeNPs had no toxicity to all cells below the dose of 50 μg/mL (Fig.2A; Fig.3A). However, when the dosage was over 50 μg/mL, FeNPs showed significant toxicity to all cells including normal cells (Fig.2A; Fig.3A). Therefore, we used the FeNPs at the concentration of 50 μg/mL for subsequent investigation, which is equivalent to the dose of intravenous injection of 3 mg/kg in rodents [26].

**Figure 2.**
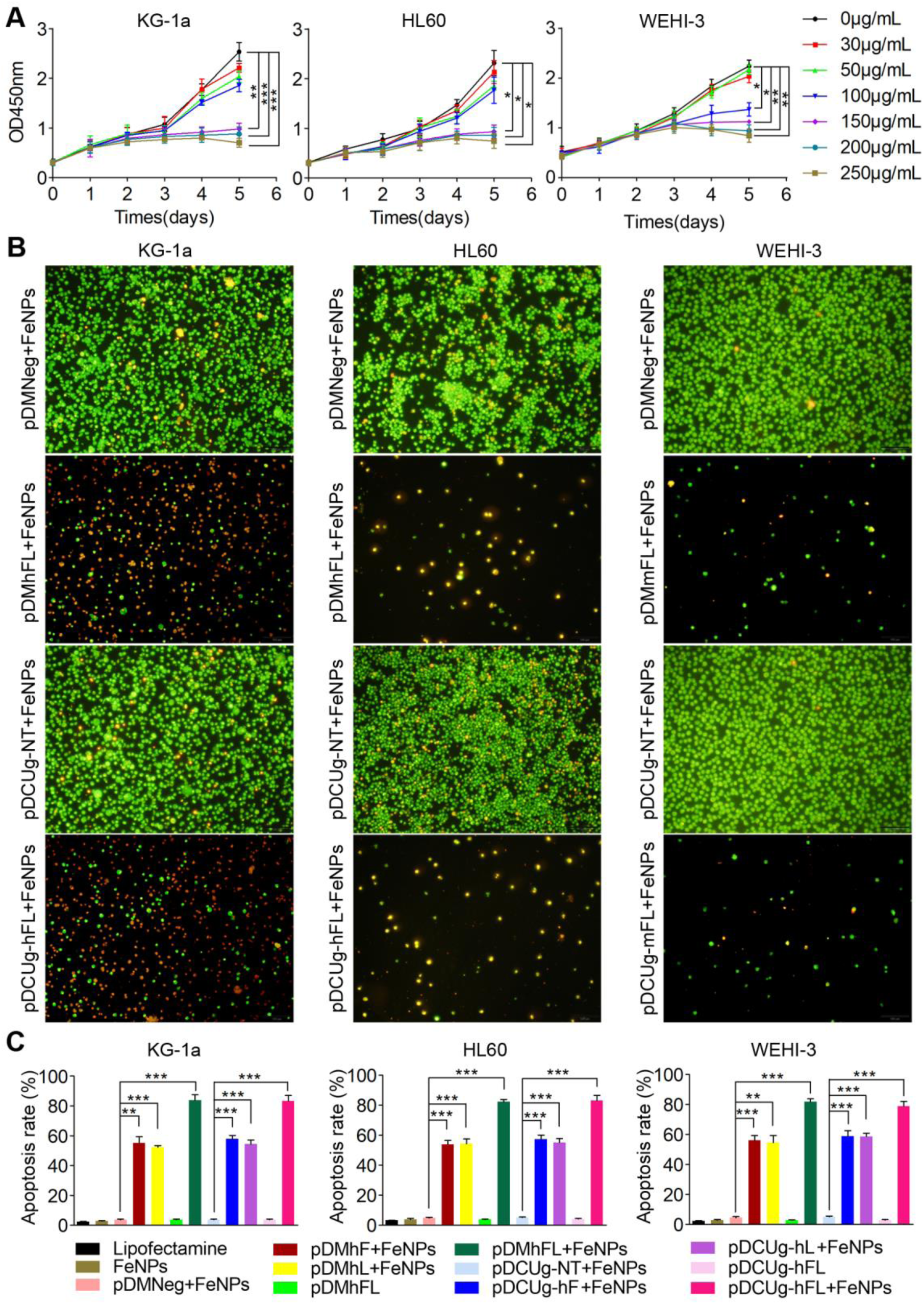
Treatment of leukemia cells with GIFT. (A) Viability of FeNPs-treated cells. (B) Cell imaging. Cells were stained by OA&EB and imaged. This figure only shows images of cells treated by pDCUg-hFL/mFL, pDMhFL/mFL, pDCUg-NT and pDMNeg together with FeNPs for 72 h. Images of cells of all other treatments are shown in Figure S1–S3. (C) Apoptosis of leukemia cells. Raw flow cytometer images are shown in Figure S4. All values are mean± s.e.m. with n=3. *, p<0.05; **, p<0.01; ***, p<0.001. Plasmid used are as follows: pDCUg is pDMP-Cas13a-U6-gRNA; pDM is pDMP-miRNA; pDCUg-NT targets no transcripts; pDCUg-hF/mF targets human or mouse (h/m) FPN; pDCUg-hL/mL targets h/m Lcn2; pDCUg-hFL/mFL targets h/m FPN and Lcn2; pDMNeg targets no transcripts; pDMhF/mF targets h/m FPN; pDMhL/mL targets to h/m Lcn2; and pDMhFL/mFL targets to h/m FPN and Lcn2.

**Figure 3.**
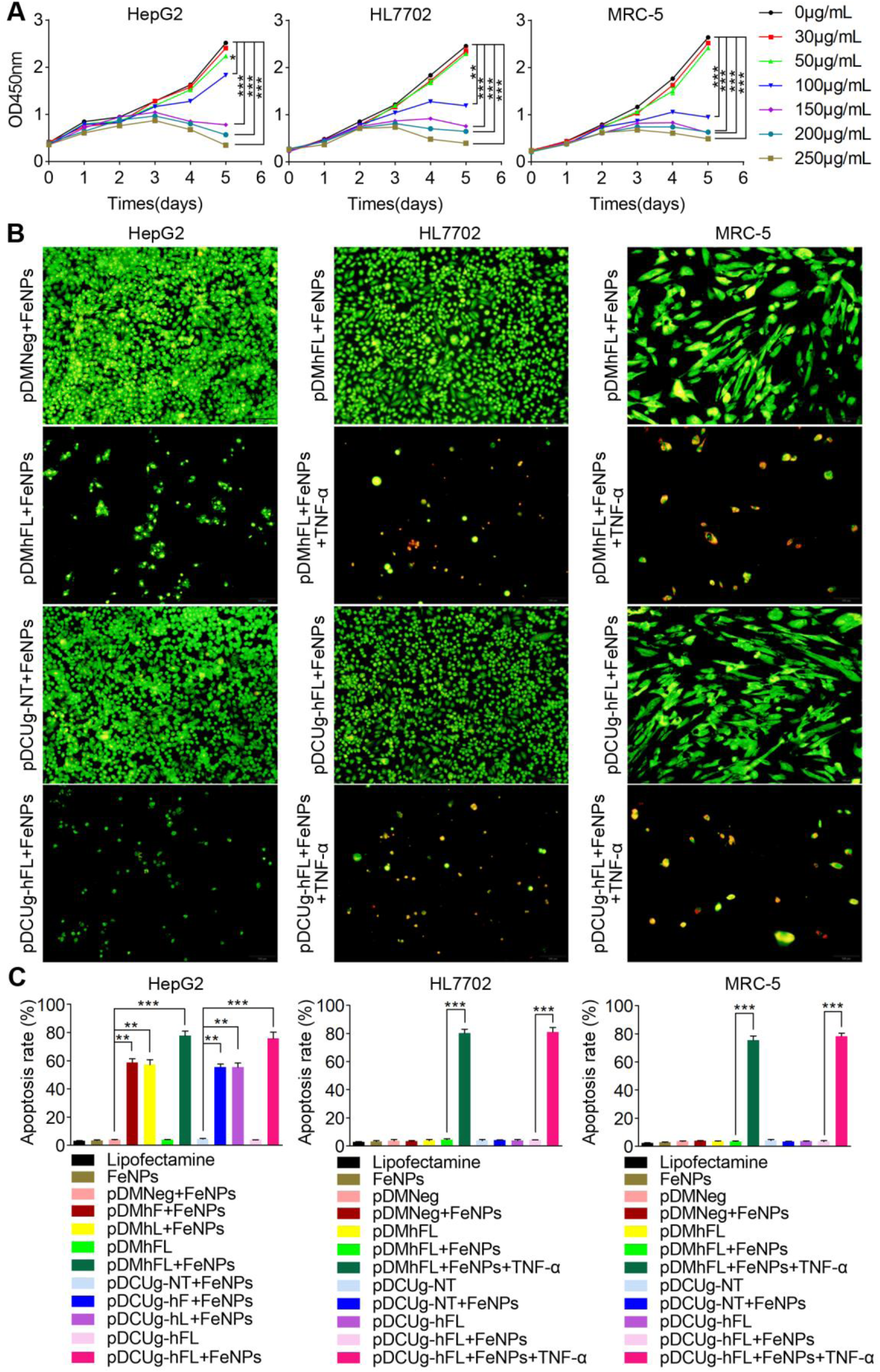
Treatment of a solid tumor and normal cells with GIFT. HepG2 is human liver cancer cell and HL7702 and MRC-5 are normal human cells. (A) Effect of FeNPs on cell viability. (B) Cell imaging. Cells were stained by OA&EB and imaged. This figure only shows images of cells treated by pDCUg-hFL, pDMhFL, pDCUg-NT and pDMNeg together with FeNPs for 72 h. Images of cells of all other treatments are shown in Figure S5–S7. (C) Cell apoptosis. Raw flow cytometer images are shown in Figure S8. All values are mean± s.e.m. with n=3. *, p<0.05; **, p<0.01; ***, p<0.001. Plasmid used are as same as Figure 2. Only vectors targeting human genes were used because the three cells are all human cells.

### 3.4. Antitumor effects of GIFT in vitro

To investigate the antitumor effects of GIFT, we first treated leukemia cells by GIFT. Three leukemia cells were first transfected by various plasmid vectors and then treated by FeNPs. The cell viability was detected by a AO&EB dual staining at three time points and measuring apoptosis at the end. The results showed that only the combinations of all GIVs with FeNPs caused the significant time-dependent apoptosis (Figures S1–S4; Figures 2B and 2C). All plasmids and FeNPs alone and the combination of negative control plasmids (pDCUg-NT and pDMNeg) and FeNPs did not affect cell viability (Figures S1–S3). Especially, the co-expressed GIVs (pDCUg-hFL/pDCUg-mFL and pDMhFL/ pDMmFL) showed the most significant cancer cell killing effect when combining with FeNPs (Figures 2B and 2C; Figures S1–S4), suggesting a synergistic effect of co-interfering two genes. These same effects of GIFT were then found on a human solid tumor cell HepG2 (Figure S5 and S6; Figures 3B and 3C).

To investigate the cancer cell specificity of GIFT, we next treated two normal human cells (HL7702 and MRC5) as treated HepG2. The results showed that all vectors alone and their combination with FeNPs did not significantly affect the two cells (Figures S6–S8), which is consistent with no NF-κB expression in the two cells (Figure 1C). To further validate the key role of NF-κB activation in GIFT, we transfected the two cells with pDCUg-hFL and pDMhFL respectively and then induced them with a NF-κB activator, TNF-α. The cells were then treated by FeNPs. We found that the two cells were also significantly killed by GIFT with the TNF-α inducement (Figures 3B and 3C; Figures S6–S8). This confirms that cells can be killed by GIFT only when only when NF-κB is activated, which is also confirmed by the treatment of HEK-293T cell with GIFT. The cell is a human embryonic kidney cell that expresses large T antigen after being transfected with a virus. Although the cell is not considered as a cancer cell, its NF-κB expression is significantly activated (Figure 1C). Therefore, the same GIFT effects as cancer cells were seen in this cell (Figure S9).

To investigate the wide spectrum of GIFT in killing cancer cells, we next similarly treated a variety of cancer cells representing different solid tumors in humans and mice, including A549, HT-29, C-33A, SKOV3, PANC-1, MDA-MB-453, BGC-823/MGC-803/SGC-7901, KYSE450/KYSE510, Hepa1-6, and B16F10, with the co-expressed GIVs (pDCUg-hFL/pDCUg-mFL and pDMhFL/pDMmFL). The results revealed that the combination of these GIVs with FeNPs produced the time-dependent significant killing effects in all cells (Figures S10–S22). Similarly, the vectors and FeNPs alone and the combination of negative control plasmids (pDCUg-NT and pDMNeg) with FeNPs did not significantly affect all cells at any processing time (Figure S10–S22).

To evaluate the interfering effect of DMP-Cas13a/U6-gRNA and DMP-miR systems, we next detected the expression of FPN and Lcn2 genes in treated cells. The results revealed that the expression of the two genes were significantly knocked down at both mRNA (Fig.4A) and protein (Fig.4B) levels by the targeting gRNAs/miRNAs in all detected cancer cells (KG-1a, HL-60 and HepG2). However, no changes were found in the HL7702 cell, further indicating the cancer cell specificity of the designed gene interfering systems. To further explore the cancer cell-specific expression of effector gene, Cas13a mRNAs were detected in the treated cells. The results revealed that Cas13a only expressed in all cancer cells treated by pDCUg vectors (Fig. 4A).

**Figure 4.**
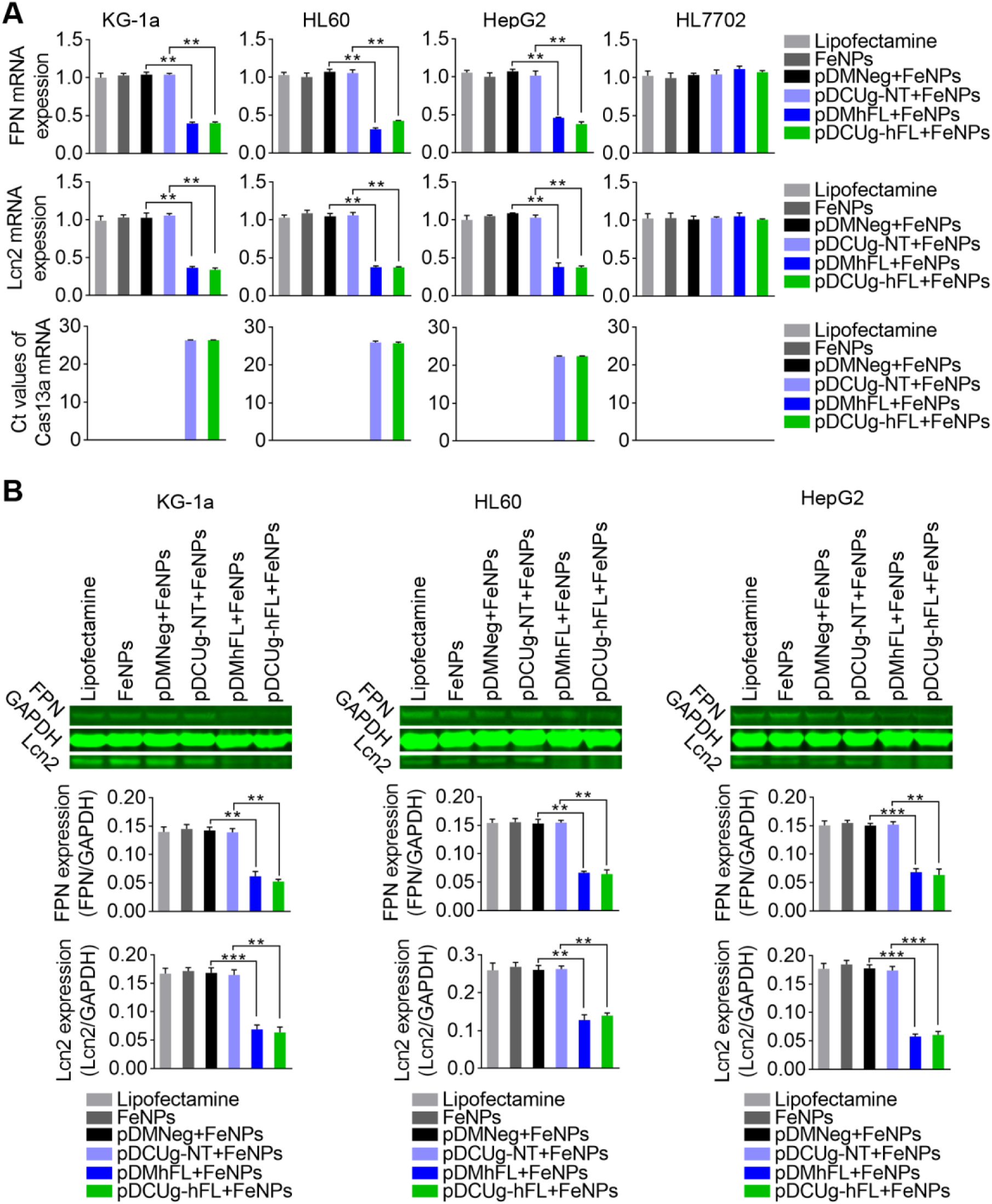
Knockdown effect of DMP-Cas13a/U6-gRNA and DMP-miR systems. Cells were transfected by vectors and cultured for 24 h, and then incubated with or without 50 μg/mL of FeNPs. Gene expression of cells were detected at 48 h post FeNPs administration. (A) QPCR analysis of mRNA expression. (B) Western blot assay of protein expression. The representative image and quantified optical density were shown. All values are mean± s.e.m. with n= 3. *, p < 0.05; **, p < 0.01; ***, p < 0.001.

It has been reported that iron-based nanomaterial can up-regulate ROS levels through the Fenton reaction [20]. To investigate whether ROS was produced in the FeNPs-treated cells, we measured the ROS levels in four cells under the various treatments. The results revealed that the combinations of GIVs with FeNPs resulted in the highest levels of ROS in all cancer cells (Figure 5A; Figure S23). However, the FeNPs alone and the combinations of negative control vectors with FeNPs only resulted in a little increase of ROS in all cancer cells (Figure 5A; Figure S23). In contrast, all the same treatments only resulted in a little increase of ROS in two normal cells (Figure 5A; Figure S23). However, when two normal cells were induced by TNF-α, the combinations of GIVs with FeNPs immediately resulted in high ROS levels in two normal cells (Figure 5A; Figure S23).

**Figure 5.**
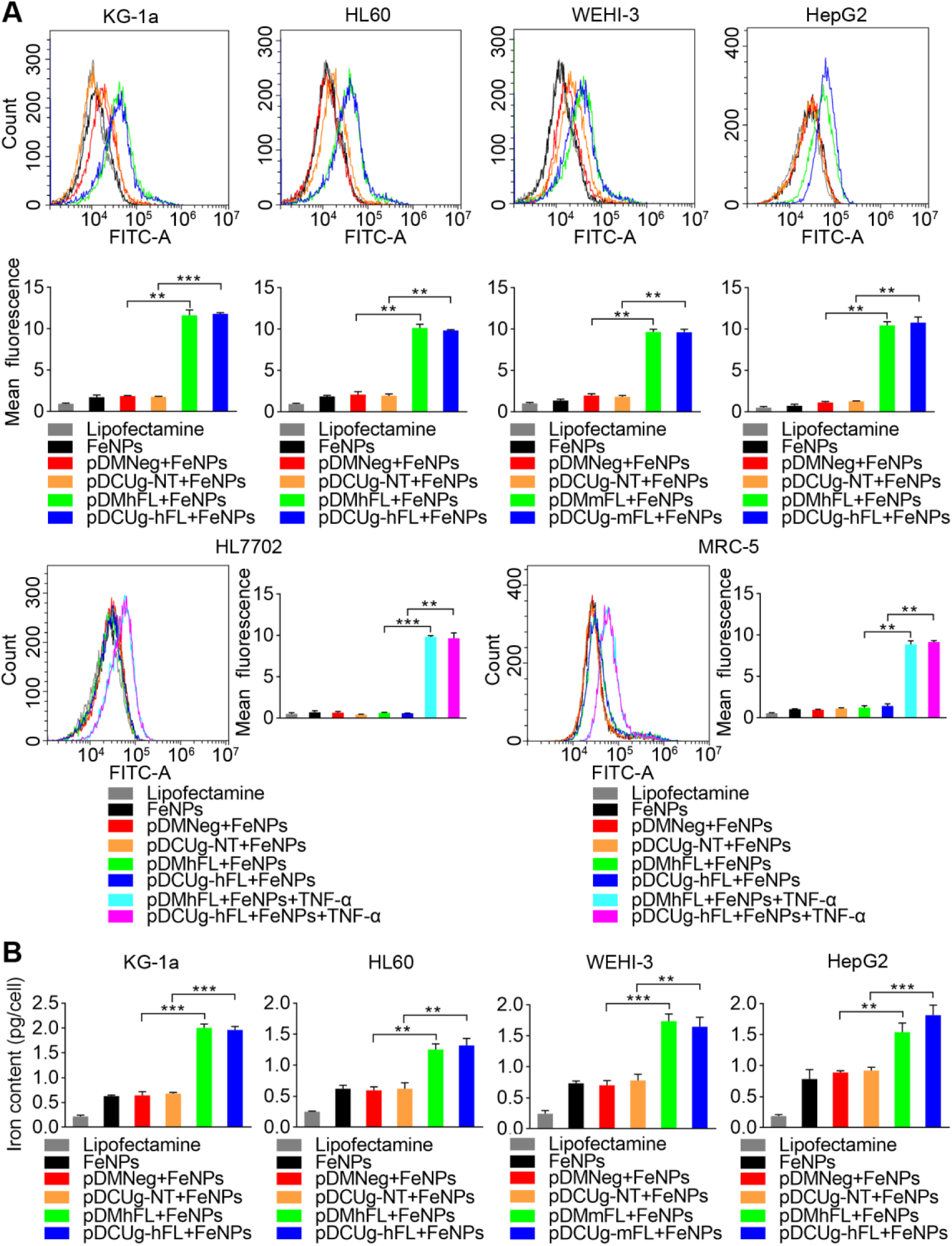
ROS production and iron content in GIFT-treated cells. Cells were assayed at 48 h post FeNPs treatment. (A) Measurement of ROS levels. The fluorescence shift and quantified fluorescence intensity are shown. (B) Measurement of iron content. All values are mean± s.e.m. with n= 3. *, p < 0.05; **, p < 0.01; ***, p < 0.001.

To further confirm the origin of ROS, we next measured the iron content in the treated cells. The results revealed that the intracellular iron content of four cancer cells significantly increased under the treatment of FeNPs together with GIVs (Fig.5B). However, the FeNPs alone and the combinations of negative control vectors with FeNPs only resulted in a limited increase of iron content (Fig.5B). These data are consistent with the ROS levels in the four cancer cells under the same treatment. These data also shows that the iron efflux was significantly inhibited by the knockdown of FPN and Lcn2 in cancer cells, which leads to a great increase of intracellular iron content and ROS. These results indicated that the mechanism underlying GIFT is genuinely the gene interfered ferroptosis.

### 3.5. Antitumor effects of virus-based GIFT in vitro

To explore the in vivo anti-tumor effects of GIFT, we next cloned the GIVs into AAV to prepare recombinant viruses (rAAV). The rAAVs were tested by transfecting three cells (KG-1a, WEHI-3 and HL7702). The results showed that the combination of gene interfering rAAVs (rAAV-DCUg-hFL/mFL and rAAV-DMhFL/DMmFL) with FeNPs resulted in significant death of two cancer cells. However, all rAAVs and FeNPs alone and the combination of negative control rAAVs (rAAV-DCUg-NT and rAAV-DMNeg) with FeNPs did not affect the growth of all cancer cells. In normal human cell HL7702, all treatments caused no significant cell death (Figure S24).

### 3.6. Antitumor effects of FeNCs-based GIFT in vitro

To find if IONPs can be used to transfer GIVs as AAV, we selected a PEI-modified Fe_3_O_4_ nanoparticle (referred to as FeNCs) as a DNA transfection agent. Two batches of FeNCs (FeNCs-1 and FeNCs-2) were evaluated by two experiments. In the first experiment, plasmids were added to FeNCs-1 to prepare FeNCs-1@DNA. The KG-1a and HepG2 cells were first treated with FeNCs-1@DNA (0.5 μg FeNCs-1) just for transfecting GIVs and then treated with FeNPs. The results showed only the combinations of that the FeNCs-1@DNA of GIVs (pDMhFL/pDMhFL) with FeNPs led to significant ferroptosis in two cells (Figure S25 and S26), whereas all FeNCs-1@DNA (0.5 μg FeNCs-1) and FeNPs alone and the combinations of negative control FeNCs-1@DNA (FeNCs-1@pDCUg-NT/pDMNeg) with FeNPs did not significantly affect the cell growth (Figure S25 and S26). These data indicate that FeNCs can be used as gene transfection agents in GIFT. In the second experiment, to further simplify the GIFT reagents, we removed FeNPs and treated the KG-1a cell only with FeNCs. The results indicated that the treatment of FeNCs@DNA (50 μg FeNCs) induced significant ferroptosis (Figure S27), whereas FeNCs (50 μg FeNCs) and DNA alone did not significantly affect the cell growth (Figure S27). To investigate the stability of FeNCs@DNA, we also treated the KG-1a cell with the FeNCs@DNA kept for 24 h. The results showed that FeNCs@DNA still had similar cancer cell killing effect.

### 3.7. Antitumor effect of GIFT in vivo

To investigate the in vivo antitumor effect of GIFT, we treated the cancer cell xenografted mouse with GIFT. The WEHI-3 cell was subcutaneously transplanted into BALB/c mice to make tumor-bearing mice. A total of three batches of animal treatments were performed. In the first batch of animal treatment, the tumor-bearing mice were treated with rAAV-DCUg-mFL. Six groups of mice were treated with variant reagents, respectively. The results showed that only the rAAV-DCUg-mFL+FeNPs treatment significantly inhibited tumor growth (Figures 6A and 6B). In the second batch of animal treatment, the tumor-bearing mice were treated with rAAV-DMmFL. Five groups of mice were treated with variant reagents, respectively. The results indicated that only the rAAV-DMmFL+FeNPs treatment significantly inhibited tumor growth (Figures 6A and 6B). In the third batch of animal treatment, the tumor-bearing mice were treated with FeNCs@rAAV-DCUg-mFL/DMmFL. Six groups of mice were treated with variant reagents, respectively. The results revealed that only the FeNCs@pAAV-DCUg-mFL/DMmFL treatments significantly inhibited tumor growth (Figures 7A and 7B). Any IONPs alone and their combination with negative control virus and nanocarrier did not affect tumor growth (Figures 6A and 6B; 7A and 7B).

**Figure 6.**
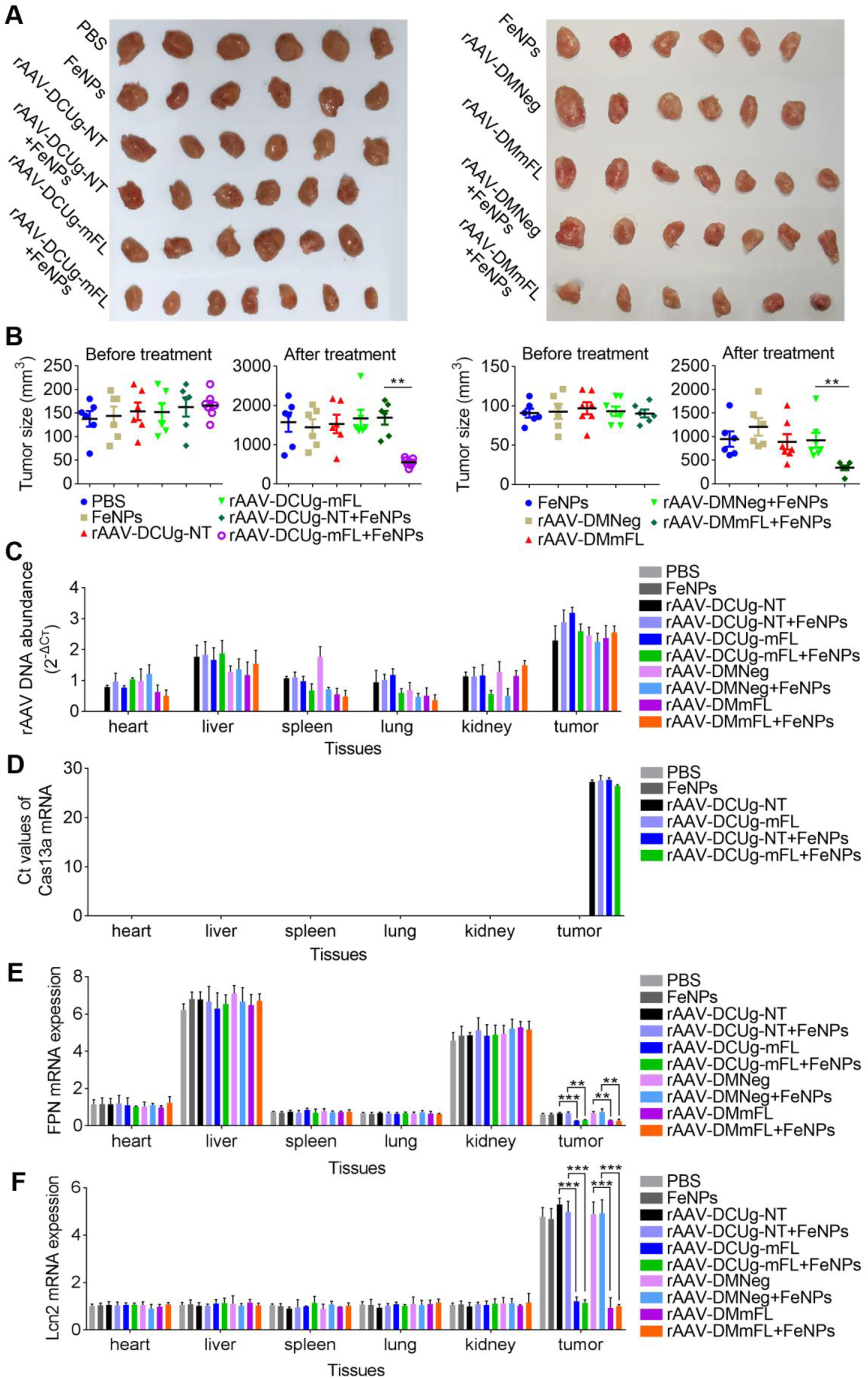
The in vivo antitumor effects of virus-based GIFT. (A) Tumors of mice treated by rAAV-DCU-g/ rAAV-DM and FeNPs. (B) Tumor volumes before and after treatment. (C) Abundance of virus DNA in tissues. (D) Cas13a expression in tissues. (E) FPN expression in tissues. (F) Lcn2 expression in tissues. RQ, relative quantification. All values are mean± s.e.m. with n= 3. *, p < 0.05; **, p < 0.01 ; ***, p < 0.001.

**Figure 7.**
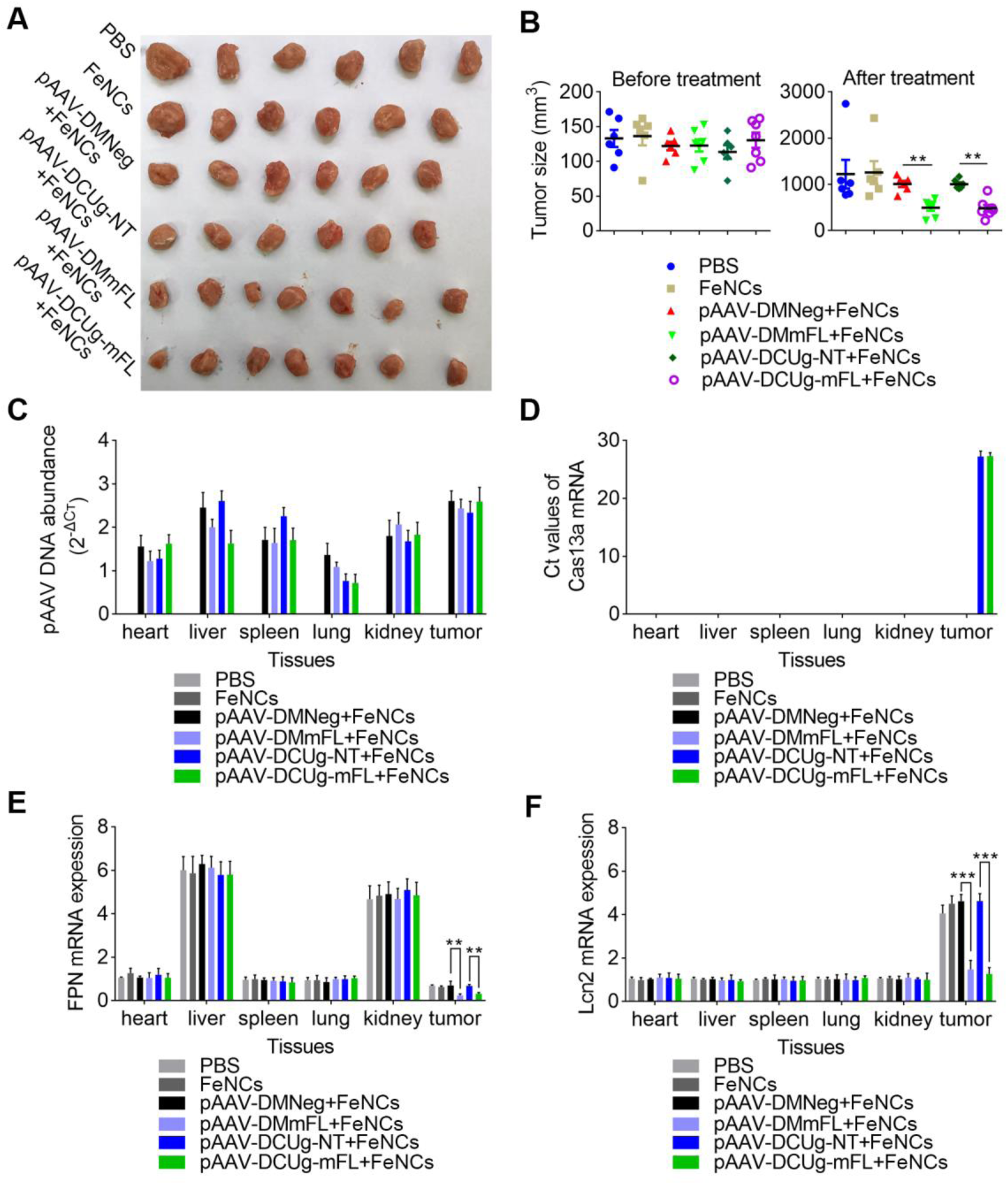
The in vivo antitumor effects of FeNCs-based GIFT. (A) Tumors of mice treated by FeNCs@DNA. (B) Tumor volumes before and after treatment. (C) Abundance of plasmid DNA in tissues. (D) Cas13a expression in tissues. (E) FPN expression in tissues. (F) Lcn2 expression in tissues. RQ, relative quantification. All values are mean± s.e.m. with n= 3. *, p < 0.05; **, p < 0.01; ***, p < 0.001.

To further confirm the molecular mechanism underlying GIFT, we detected the abundance of GIV DNA and the transcriptions of effector and target genes in various tissues. The results showed that the vector’s DNA distributed in all detected tissues, especially in tumor (Figures 6C and 7C). The transcription of effector gene Cas13a only expressed in tumor (Figures 6D and 7D), indicating the in vivo cancer-specificity of DMP-controlled effector gene expression. The detection of the expression of two target genes revealed that FNP was highly expressed in liver and kidney, while Lcn2 was highly expressed in tumor (Figures 6E and 6F; 7E and 7F). However, their expression was significantly knocked down only in tumor by the treatments of the combination of the GIVs (rAAV-DCUg-mFL/DMmFL and pAAV-DCUg-mFL/DMmFL) with IONPs (FeNPs and FeNCs) (Figures 6E and 6F; 7E and 7F). Their expression was not affected by any viruses and IONPs alone and the combination of the negative control GIVs (rAAV-DCUg-NT/DMNeg and pAAV-DCUg-NT/DMNeg) with IONPs (FeNPs and FeNCs) in all tissues (Figures 6E and 6F; 7E and 7F).

## 4. Discussion

Drug low response and resistance is a huge and wide obstacle to various cancer treatments. It is therefore urgent to find new treatment strategies for patients having no response or no longer benefiting from current therapies. Currently, IONPs have been successfully used as MRI contrast agent in cancer diagnosis and anemia treatment [38-40]. However, IONPs have not yet been used in the cancer treatment. Nevertheless, many studies have shown the intracellular degradation, release of iron ions, increase of intracellular ROS, and induction of apoptosis of IONPs. This process coincides with the mechanism of ferroptosis that has been intensively studied in recent years. Therefore, IONPs-induced ferroptosis provides a new hopeful cancer therapy. However, due to the importance of iron to cells, cells have evolved a set of mechanisms to maintain intracellular iron homeostasis. With these mechanisms, cells can effectively store and export the intracellular excess iron ions. Therefore, the IONPs-induced ferroptosis is too weak to treat cancer.

In this study, to validate GIFT, we selected FPN and Lcn2 as two target genes based on previous studies. In an our previous study, we found significant up-regulation of transcription of important genes responsible for exporting intracellular iron ion in cells treated by a DMSA-coated Fe_3_O_4_ nanoparticle, such as FPN and Lcn2 [29]. At present, ferroportin (FPN) is the only known cellular iron exporter and its loss- or gain-of-function disturbs iron export activity and induces iron metabolism disorders [41-43]. FPN has been found to be dysregulated in many cancers, such as breast tumor, prostate, ovarian, colorectal and multiple myeloma [44-49]. The enhanced expression of lipocalin-2 (Lcn2) is observed in many malignancies, which has reported as an alternative pathway for physiological iron delivery in recent years [50, 51]. Lcn2 is overexpressed in a variety of human cancers including breast, liver, and pancreatic cancer and facilitates tumorigenesis by promoting survival, growth, and metastasis [52]. The detection of Lcn2 expression in various mouse tissues in this study also revealed that the expression of Lcn2 is much higher in tumor than in other normal tissues (Figures 6F and 7F). A recent study revealed that a CRISPR/Cas9-mediated knockdown of Lcn2 significantly inhibits migration of human triple-negative breast cancer (TNBC) cells of the mesenchymal phenotype, weakens TNBC aggressiveness, and suppress in vivo TNBC tumor growth [53]. Therefore, we deduced that inhibition of the two genes should be fetal for the IONPs-treated cancer cells, because their inhibition would prevent cell from exporting IONPs-produced iron ions, which can thus enhance IONPs-induced ferroptosis.

At present, knocking down gene expression is easy by using several gene interfering tools such as siRNA and Cas13a. However, the key problem is how to selectively knock down target genes only in cancer cells. Otherwise, the IONPs-induced ferroptosis enhanced by gene interfering similarly damage normal cells. In recent years, we have developed and verified a new type of cancer cell-specific promoter that is consists of a NF-κB decoy and a minimal promoter (DMP) [34, 54]. Therefore, we used the promoter to control the expression of gene interfering vectors targeting FPN and Lcn2 in this study. The results demonstrated the promoter successfully control two gene interfering systems, Cas13a and miRNA [55, 56], to selectively function in cancer cells in vitro and in vivo.

In this study, we developed GIFT to cancers. Our study revealed GIFT is effective in all detected cancer cells representing various hematological and solid tumors, indicating GIFT is in potential a wide-spectrum cancer therapy. The mechanism underlying this new therapy is gene interfering-enhanced ferroptosis. This study demonstrated that GIFT only functioned with GIVs and IONPs together. Any GIVs and IONPs alone have no effects. It should be noted that GIFT is also very effective to cancer cells of clinical refractory cancers such as TNBC (MDA-MB-453) and pancreatic cancer (PANC-1). This is coincident with a recent new study that induced pancreatic tumor ferroptosis in mice by depleting cysteine via deletion of a system x_C_–subunit, *Slc7a11*, or administration of cyst(e)inase [57].

For the potential clinical application, this study tested three kinds of *in vivo* administration approaches of GIFT reagents. The first is two-time intravenous injection of rAAV and FeNPs. The second is one-time intravenous injection of rAAV and FeNPs. The third is one-time intravenous injection of FeNCs@DNA. The mice treatments revealed that three forms all functioned well, indicating both viral and non-viral vectors can be used by GIFT. AAV has already become a safe virus vector for the current human gene therapy [58]. However, it is still challenged by high production cost and sometimes immunological rejection. Therefore, the non-viral vectors such as nanomaterial have been rapidly developed [59], such as IONPs [60]. IONPs have already be used as clinical therapeutic and diagnostic reagents for their good biocompatibility [38-40]. This study tested a treatment of tumor-bearing mice with FeNCs@DNA. In this case, FeNCs take dual functions of IONPs to provide iron ions and non-viral vector to transfer GIVs.

In this study, we tested two gene interfering tools, CRISPR/Cas13a-gRNA and miRNA. In combination with DMP, they all functioned well both in cells and mice. Because AAV has a limited DNA packaging capacity (4Kb), several advantages of DMP and two gene interfering tools are helpful for packaging them into AAV. First, DMP is very short promoter (84 bp). Second, Cas13a can process its own gRNA precursors. Third, miRNA expression backbone is only 390 bp. These features is beneficial for packaging a GIV that co-expresses gRNAs or miRNAs targeting multiples genes in AAV. For example, this study constructed GIVs co-expressing gRNAs and miRNAs targeting both FPN and Lcn2 (pDCUg-hFL/pDCUg-mFL and pDMhFL/pDMmFL) and successfully packaged them into AAV. This is beneficial for taking advantage of synergistic or additive effect of different genes like FPN and Lcn2 used here.

## 5. Conclusion

We developed a new cancer therapy, gene-interfered ferroptosis therapy (GIFT), by combining cancer-specific gene interfering and IONPs-induced ferroptosis. In mechanism, it is in fact a gene interfering-enhanced ferroptosis. By targeting two genes (FPN and Lcn2) by a cancer cell-specific promoter (DMP) and two gene interfering tools (CRISPR/Cas13 and miRNA) and using two kinds of IONPs, we verified the in vitro anti-tumor effect of GIFT in a variety of cancer cells. We also verified the in vivo anti-tumor effect of GIFT by treating tumor-bearing mice with the AAV plus FeNPs and FeNCs@DNA. By performing well-designed experiments, we also validate the underlying mechanism, cancer specificity and wide spectrum of GIFT.

## Supplementary information

The supplementary information includes Figures S1–S27 (treatment of various cancer cells representing variant hematological and solid tumors with GIFT; flow cytometry assay of cell apoptosis), Tables S1–S3 (gRNA and microRNA target sequences and PCR primers), and vectors and their functional sequences (pDCUg, pDM, pDMhFL, and pDMmFL).

## Author Contributions

J.K.W. conceived the study and designed the experiments. J.L.G. performed main experiments. T.L. and N.L. prepared reagents and performed partial experiments. J.W. and J.L.G. wrote the manuscript with support from all authors.

## Supplementary information

**Fig.S1.**
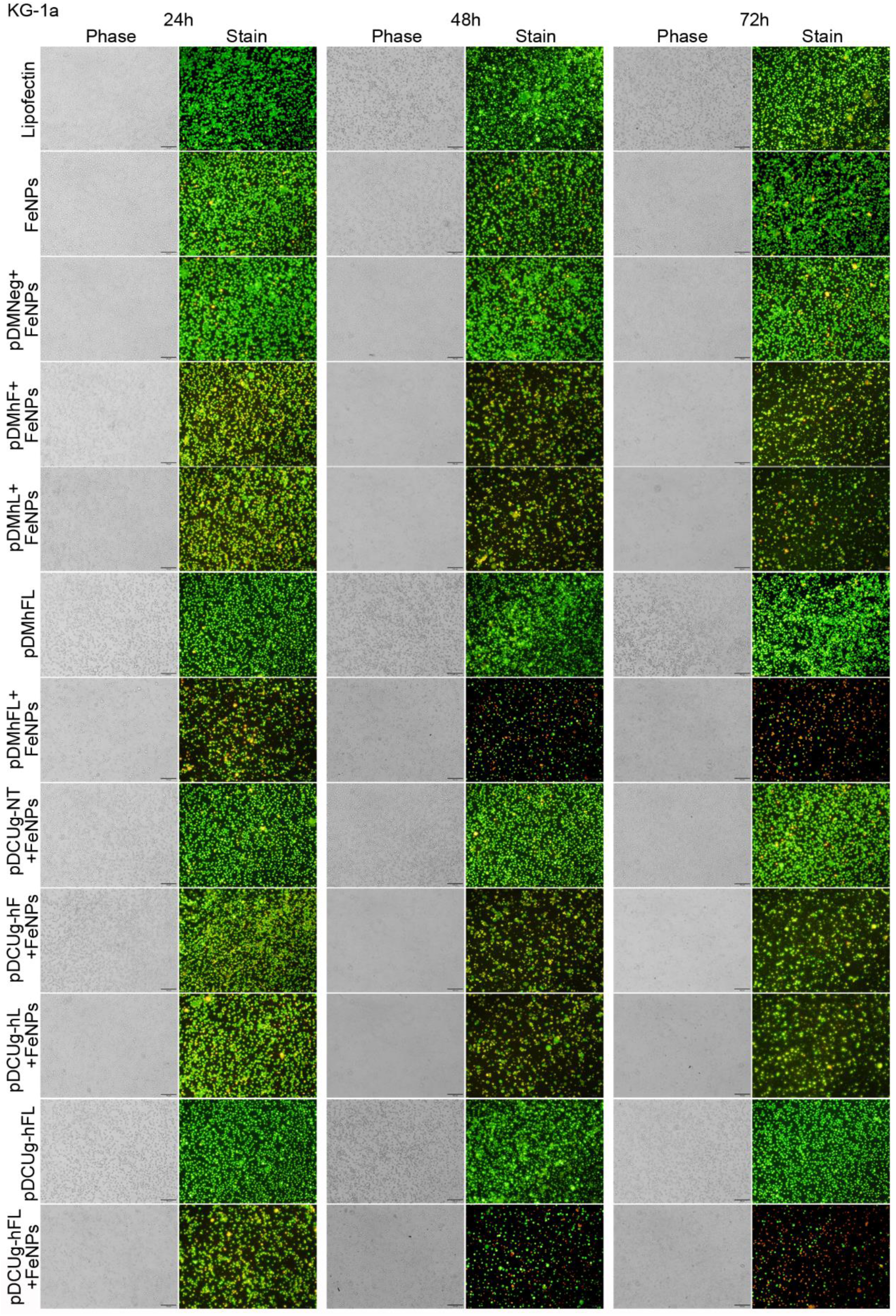
Treatment of KG-1a cells with gene interfering vectors and FeNPs.Cells were transfected by various plasmids, cultured for 24 h and then incubated with or without 50 μg/mL of FeNPs, and cells were cultured for another 72 h. At 24 h, 48 h and 72 h post FeNPs administration, cells were stained with acridine orange and ethidium bromide and imaged under a fluorescence microscope.pDMNeg/pDMhF/pDMhL/pDMhFL, plasmids expressing miRNAs under the control of DMP targeting no, human FPN,human Lcn2, or human FPN and Lcn2 transcripts;pDCUg-NT/pDCUg-hF/pDCUg-hL/pDCUg-hFL, plasmids expressing Cas13a under the control of DMP and gRNAs under the control of U6 pomoter targeting no, human FPN, human Lcn2, or human FPN and Lcn2 transcripts.

**Fig.S2.**
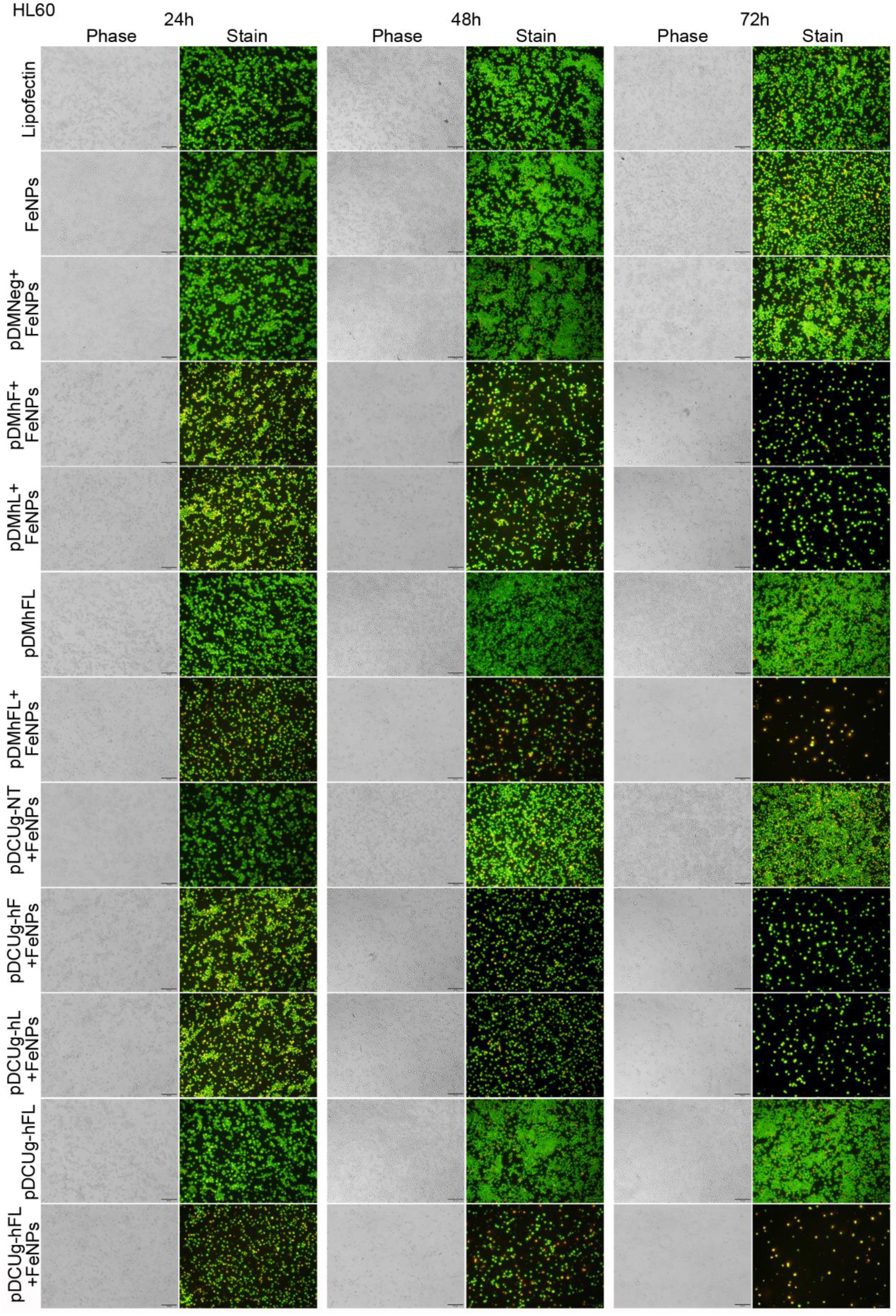
Treatment of HL60 cells with gene interfering vectors and FeNPs.Cells were transfected by various plasmids, cultured for 24 h and then incubated with or without 50 μg/mL of FeNPs, and cells were cultured for another 72 h. At 24 h, 48 h and 72 h post FeNPs administration, cells were stained with acridine orange and ethidium bromide and imaged under a fluorescence microscope. pDMNeg/pDMhF/pDMhL/pDMhFL, plasmids expressing miRNAs under the control of DMP targeting no, human FPN,human Lcn2, or human FPN and Lcn2 transcripts;pDCUg-NT/pDCUg-hF/pDCUg-hL/pDCUg-hFL, plasmids expressing Cas13a under the control of DMP and gRNAs under the control of U6 pomoter targeting no, human FPN, human Lcn2, or human FPN and Lcn2 transcripts.

**Fig.S3.**
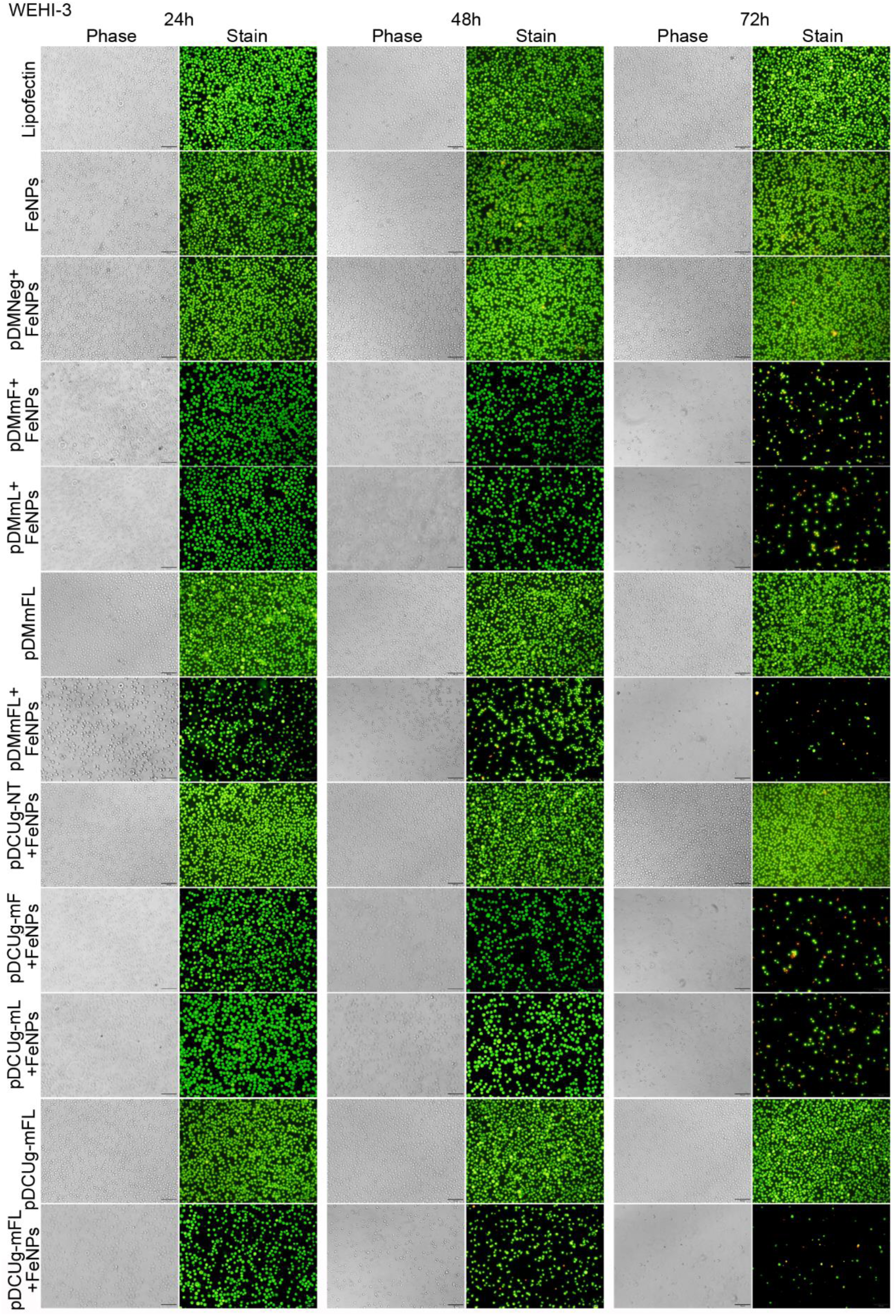
Treatment of WEHI-3 cells with gene interfering vectors and FeNPs.Cells were transfected by various plasmids, cultured for 24 h and then incubated with or without 50 μg/mL of FeNPs, and cells were cultured for another 72 h. At 24 h, 48 h and 72 h post FeNPs administration, cells were stained with acridine orange and ethidium bromide and imaged under a fluorescence microscope. pDMNeg/pDMmF/pDMmL/pDMmFL, plasmids expressing miRNAs under the control of DMP targeting no, mouse FPN, mouse Lcn2, or mouse FPN and Lcn2 transcripts;pDCUg-NT/pDCUg-mF/pDCUg-mL/pDCUg-mFL, plasmids expressing Cas13a under the control of DMP and gRNAs under the control of U6 pomoter targeting no, mouse FPN, mouse Lcn2, or mouse FPN and Lcn2 transcripts.

**Fig.S4.**
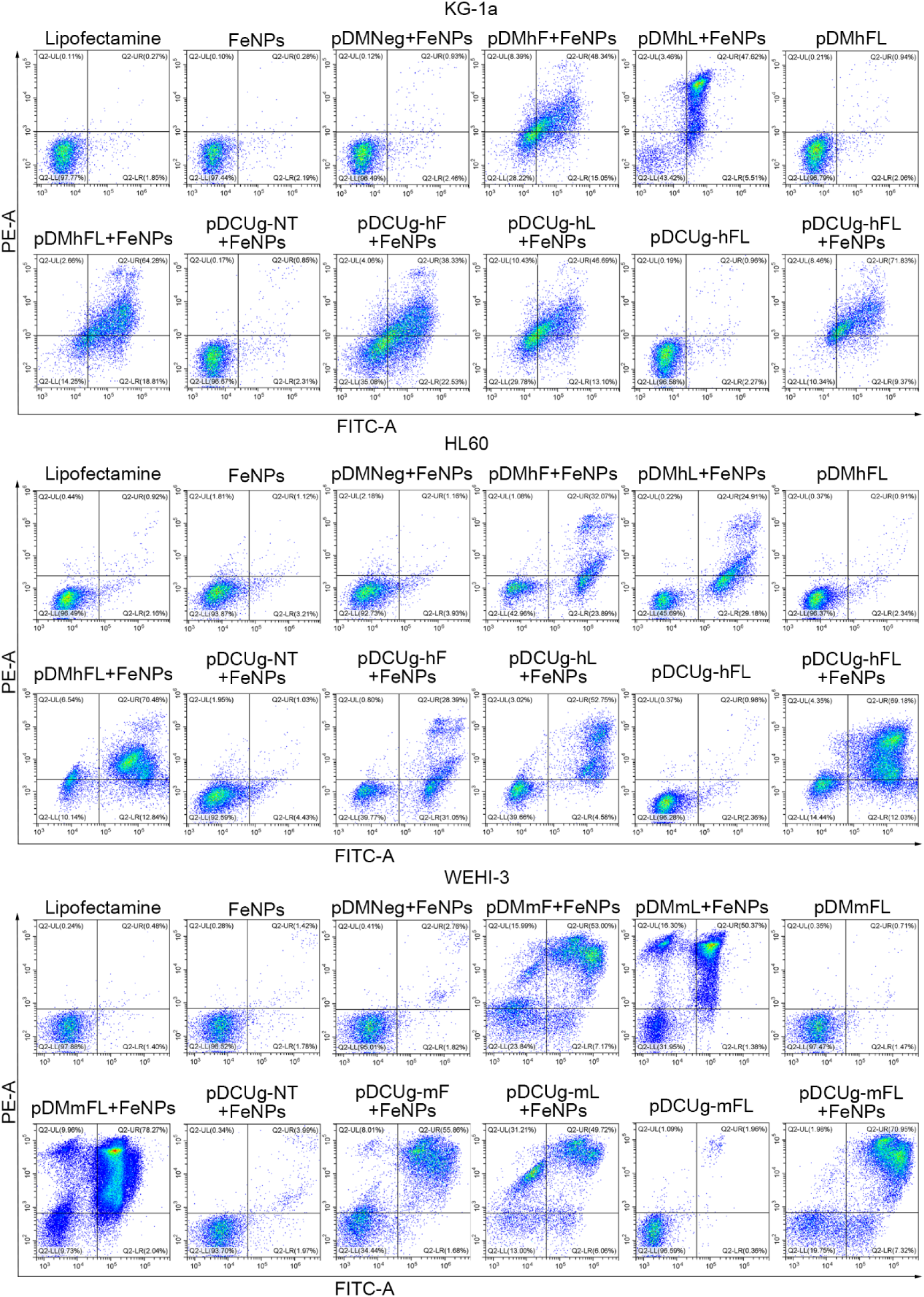
Flow cytometry analysis of apoptosis of cells treated with gene interfering vectors and FeNPs. KG-1a, HL60 and WEHI-3 cells were transfected by various plasmids, cultured for 24 h and then incubated with or without 50 μg/mL of FeNPs.Cells were cultured for another 72 h.Cells were collected at 72 h post FeNPs administration and detected with the Annexin V-FITC Apoptosis Detection Kit.The fluorescence intensity of cells was quantified by Flow Cytometer.pDMNeg/pDMhF/pDMhL/pDMhFL/pDMmF/pDMmL/pDMmFL, plasmids expressing miRNAs under the control of DMP targeting no, human FPN, human Lcn2, human FPN and Lcn2, mouse FPN, mouse Lcn2, or mouse FPN and Lcn2 transcripts; pDCUg-NT/pDCUg-hF/pDCUg-hL/pDCUg-hFL/pDCUg-mF/pDCUg-mL/pDCUg-mFL, plasmids expressing Cas13a under the control of DMP and gRNAs under the control of U6 promoter targeting no, human FPN, human Lcn2, human FPN and Lcn2, mouse FPN, mouse Lcn2, or mouse FPN and Lcn2 transcripts.

**Fig.S5.**
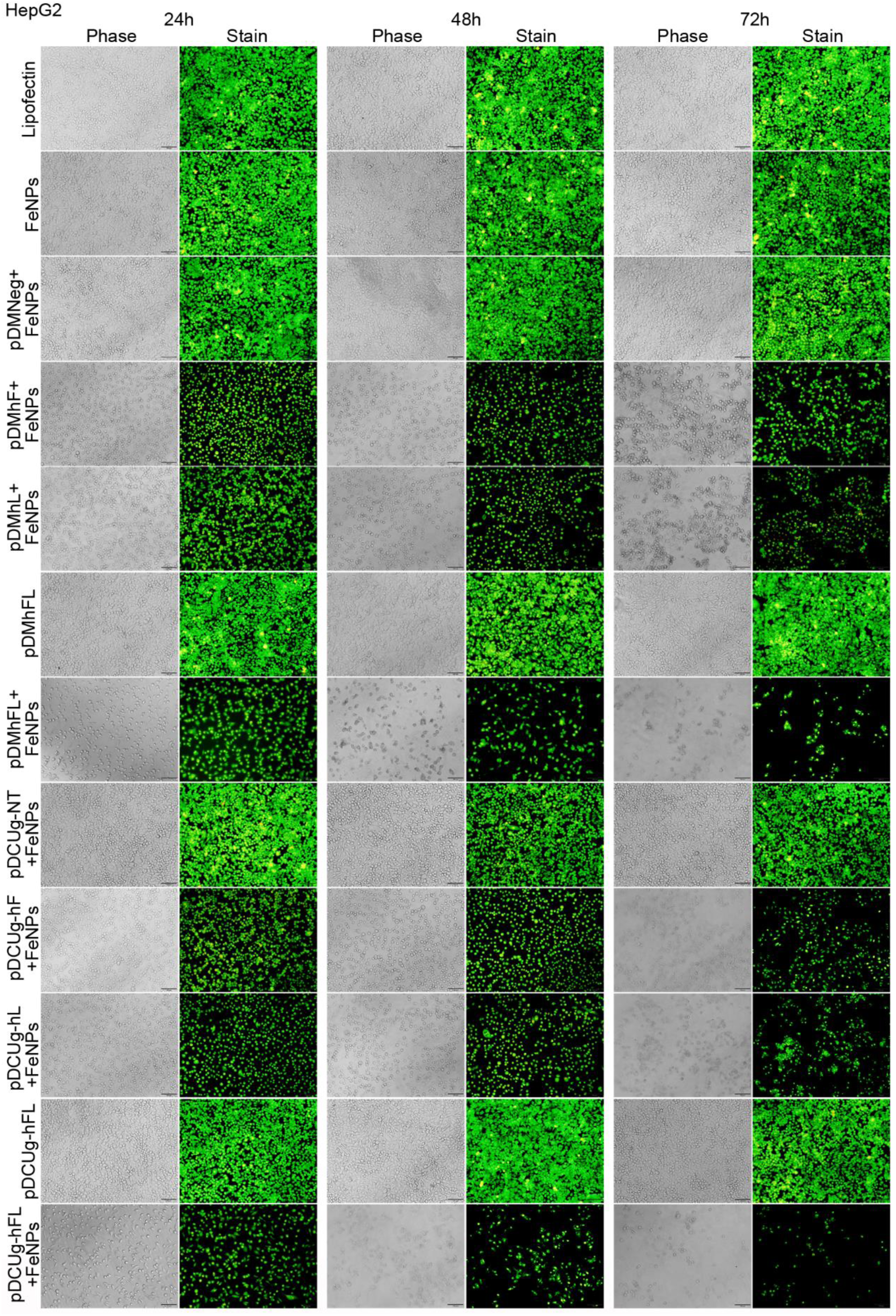
Treatment of HepG2 cells with gene interfering vectors and FeNPs.Cells were transfected by various plasmids, cultured for 24 h and then incubated with or without 50 μg/mL of FeNPs, and cells were cultured for another 72 h. At 24 h, 48 h and 72 h post FeNPs administration, cells were stained with acridine orange and ethidium bromide and imaged under a fluorescence microscope. pDMNeg/pDMhF/pDMhL/pDMhFL, plasmids expressing miRNAs under the control of DMP targeting no, human FPN, human Lcn2, or human FPN and Lcn2 transcripts; pDCUg-NT/pDCUg-hF/pDCUg-hL/pDCUg-hFL, plasmids expressing Cas13a under the control of DMP and gRNAs under the control of U6 pomoter targeting no, human FPN, human Lcn2, or human FPN and Lcn2 transcripts.

**Fig.S6.**
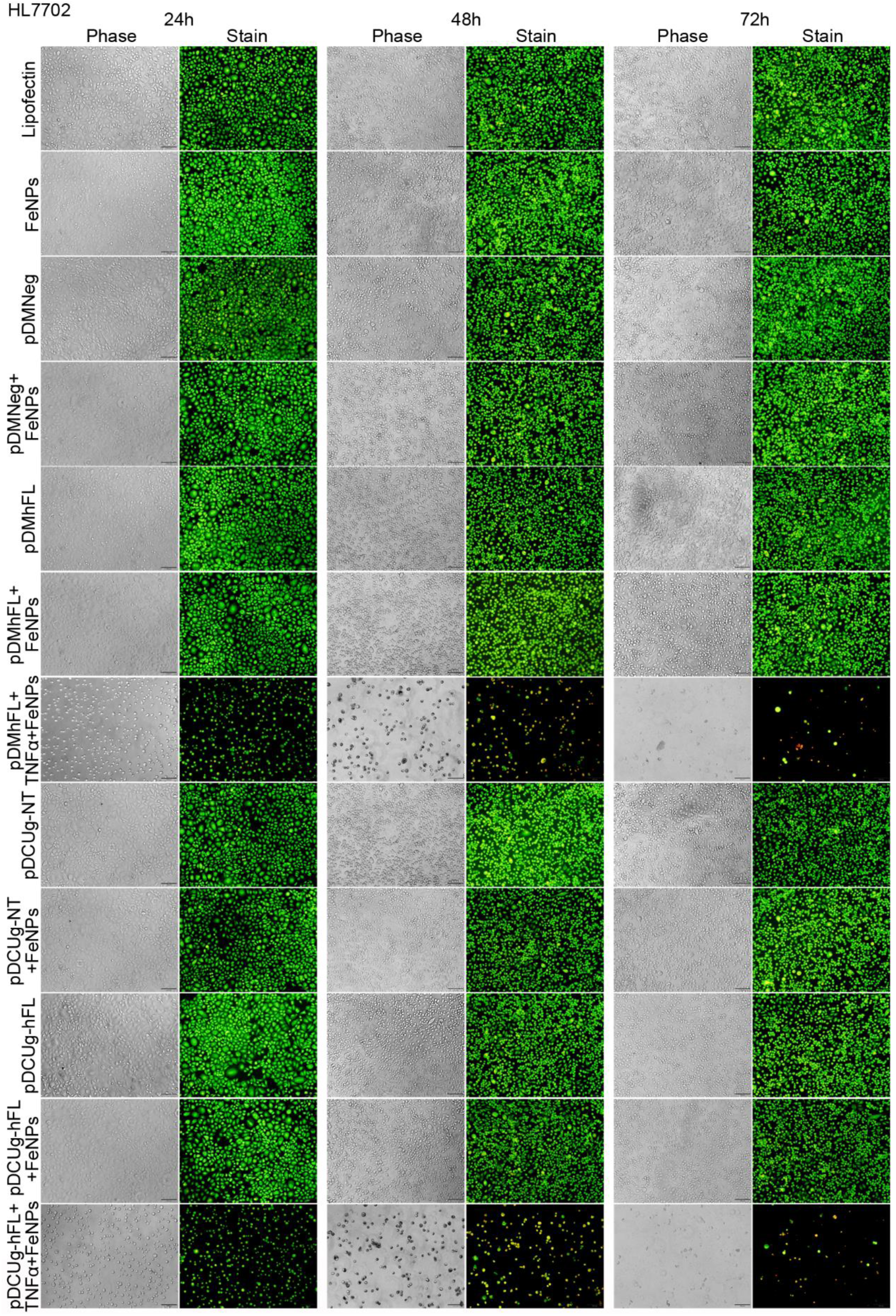
Treatment of HL7702 cells with gene interfering vectors and FeNPs.Cells were transfected by various plasmids, cultured for 24 h and then induced with or without TNF-α(10 ng/mL) for 1 h before incubating with or without 50 μg/mL of FeNPs, and cells were cultured for another 72 h. At 24 h, 48 h and 72 h post FeNPs administration, cells were stained with acridine orange and ethidium bromide and imaged under a fluorescence microscope. pDMNeg/pDMhFL, plasmids expressing miRNAs under the control of DMP targeting no, or human FPN and Lcn2 transcripts; pDCUg-NT/pDCUg-hFL, plasmids expressing Cas13a under the control of DMP and gRNAs under the control of U6 promoter targeting no, or human FPN and Lcn2 transcripts.

**Fig.S7.**
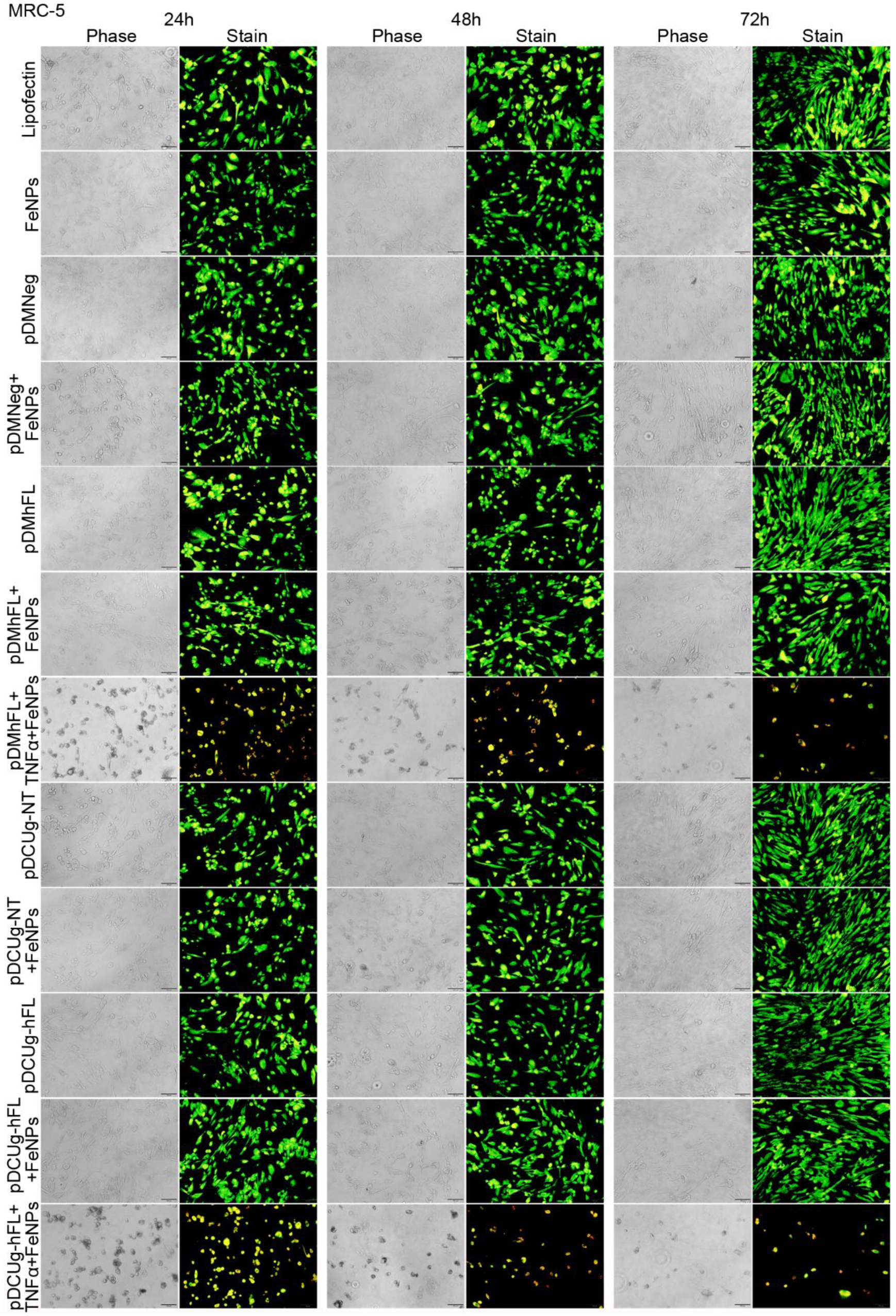
Treatment of MRC-5 cells with gene interfering vectors and FeNPs.Cells were transfected by various plasmids, cultured for 24 h and then induced with or without TNF-α (10 ng/mL) for 1h before incubating with or without 50 μg/mL of FeNPs, and cells were cultured for another 72 h. At 24 h, 48 h and 72 h post FeNPs administration, cells were stained with acridine orange and ethidium bromide and imaged under a fluorescence microscope. pDMNeg/pDMhFL, plasmids expressing miRNAs under the control of DMP targeting no, or human FPN and Lcn2 transcripts; pDCUg-NT/pDCUg-hFL, plasmids expressing Cas13a under the control of DMP and gRNAs under the control of U6 promoter targeting no, or human FPN and Lcn2 transcripts.

**Fig.S8.**
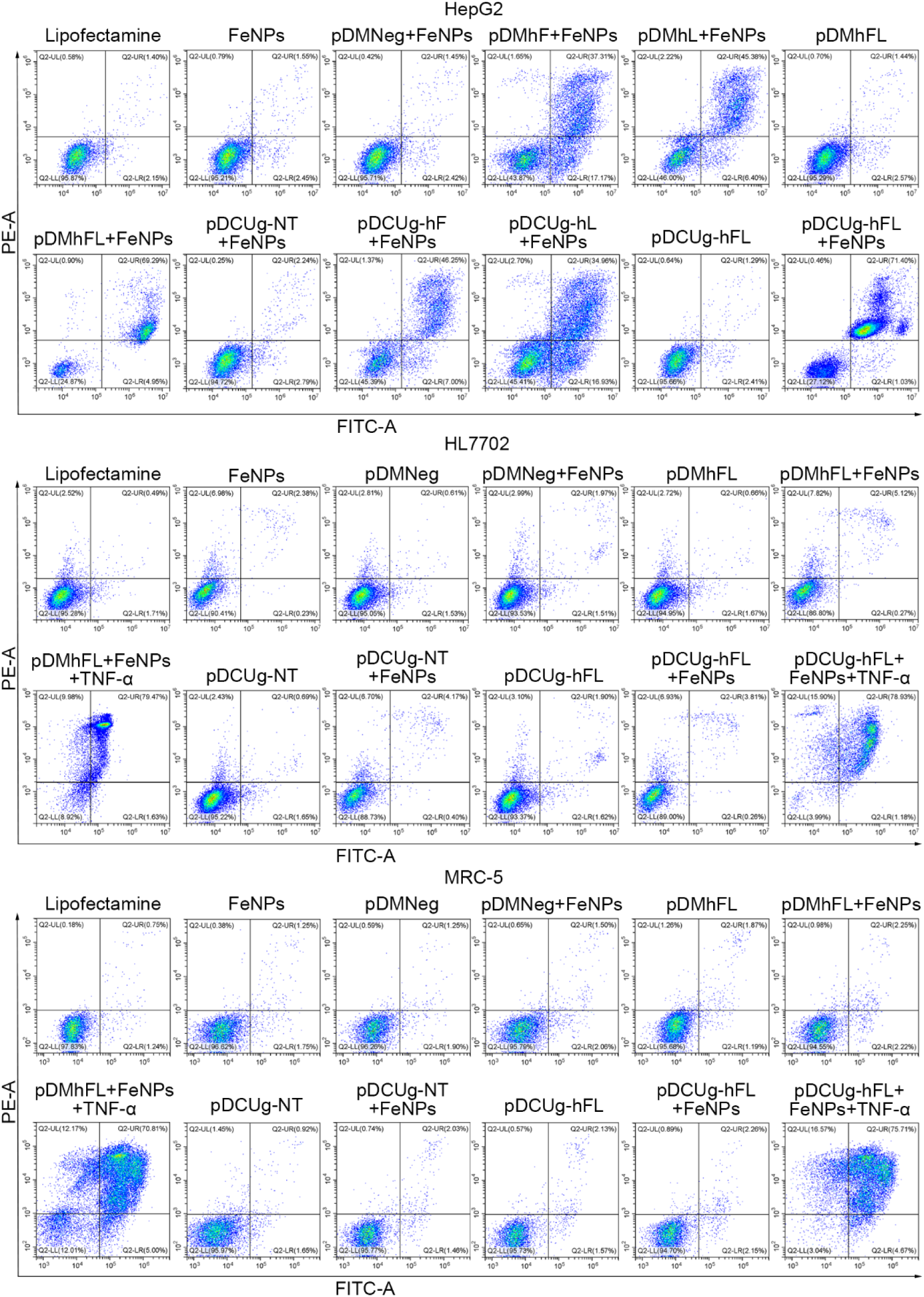
Flow cytometry analysis of cell apoptosis of cells treated with gene interfering vectors and FeNPs. HepG2, HL7702 and MRC-5 cells were transfected by various plasmids, cultured for 24 h and then incubated with or without 50 μg/mL of FeNPs, and cells were cultured for another 72 h. For HL7702 and MRC-5,cells were first induced with or without TNF-α (10 ng/mL) for 1h before FeNPs treatment. Cells were collected at 72 h post FeNPs administration and detected with the Annexin V-FITC Apoptosis Detection Kit. The fluorescence intensity of cells was quantified by Flow Cytometer. pDMNeg/pDMhF/pDMhL/pDMhFL, plasmids expressing miRNAs under the control of DMP targeting no, human FPN, human Lcn2, or human FPN and Lcn2 transcripts; pDCUg-NT/pDCUg-hF/pDCUg-hL/pDCUg-hFL, plasmids expressing Cas13a under the control of DMP and gRNAs under the control of U6 promoter targeting no, human FPN, human Lcn2, or human FPN and Lcn2 transcripts.

**Fig.S9.**
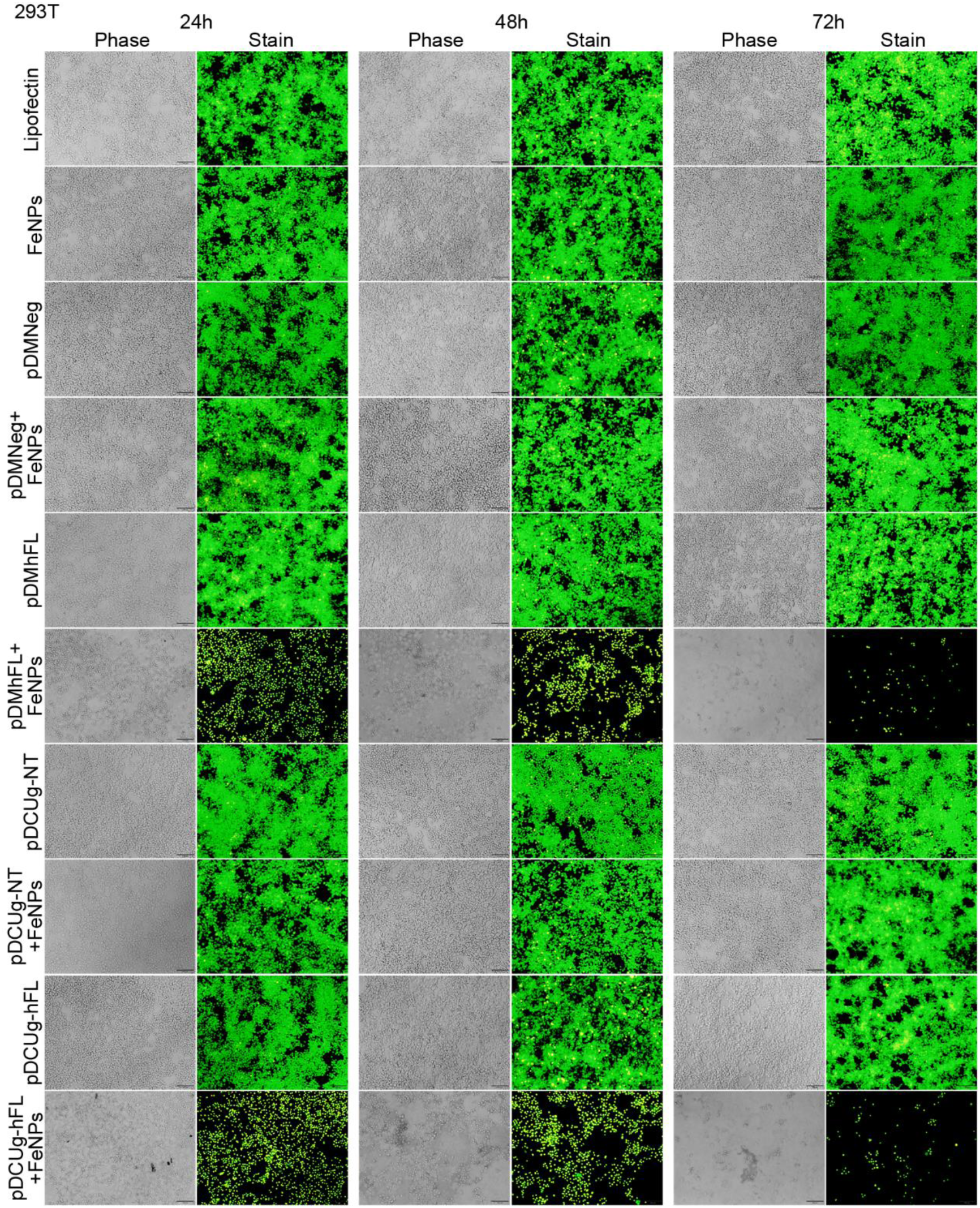
Treatment of HEK-293T cells with gene interfering vectors and FeNPs. Cells were transfected by various plasmids, cultured for 24 h and then incubated with or without 50 μg/mL of FeNPs, and cells were cultured for another 72 h. At 24 h, 48 h and 72 h post FeNPs administration, cells were stained with acridine orange and ethidium bromide and imaged under a fluorescence microscope. pDMNeg/pDMhFL, plasmids expressing miRNAs under the control of DMP targeting no or human FPN and Lcn2 transcripts; pDCUg-NT/pDCUg-hFL, plasmids expressing Cas13a under the control of DMP and gRNAs under the control of U6 promoter targeting no, or human FPN and Lcn2 transcripts.

**Fig.S10.**
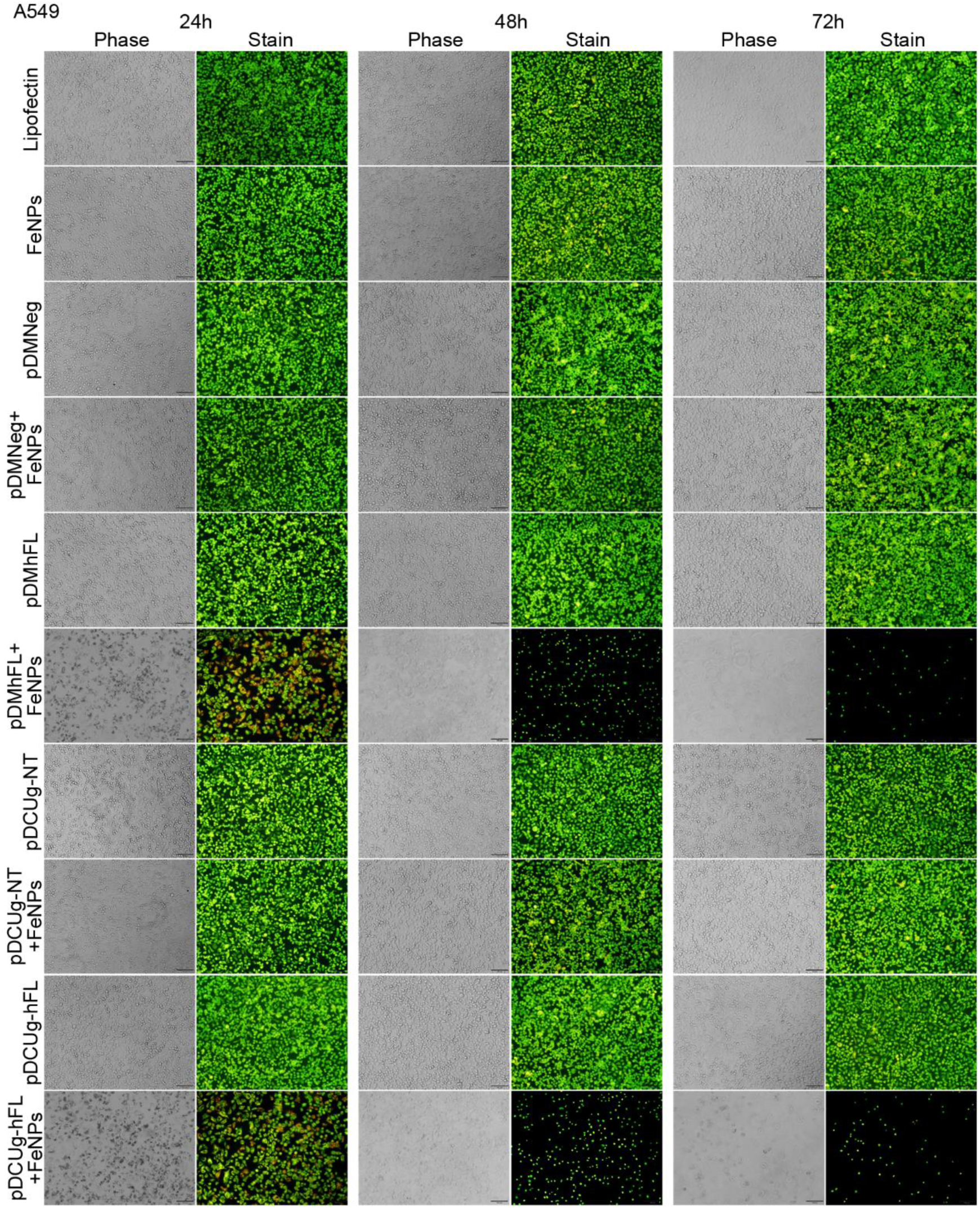
Treatment of A549 cells with gene interfering vectors and FeNPs. Cells were transfected by various plasmids, cultured for 24 h and then incubated with or without 50 μg/mL of FeNPs, and cells were cultured for another 72 h. At 24 h, 48 h and 72 h post FeNPs administration, cells were stained with acridine orange and ethidium bromide and imaged under a fluorescence microscope. pDMNeg/pDMhFL, plasmids expressing miRNAs under the control of DMP targeting no or human FPN and Lcn2 transcripts; pDCUg-NT/pDCUg-hFL, plasmids expressing Cas13a under the control of DMP and gRNAs under the control of U6 promoter targeting no or human FPN and Lcn2 transcripts.

**Fig.S11.**
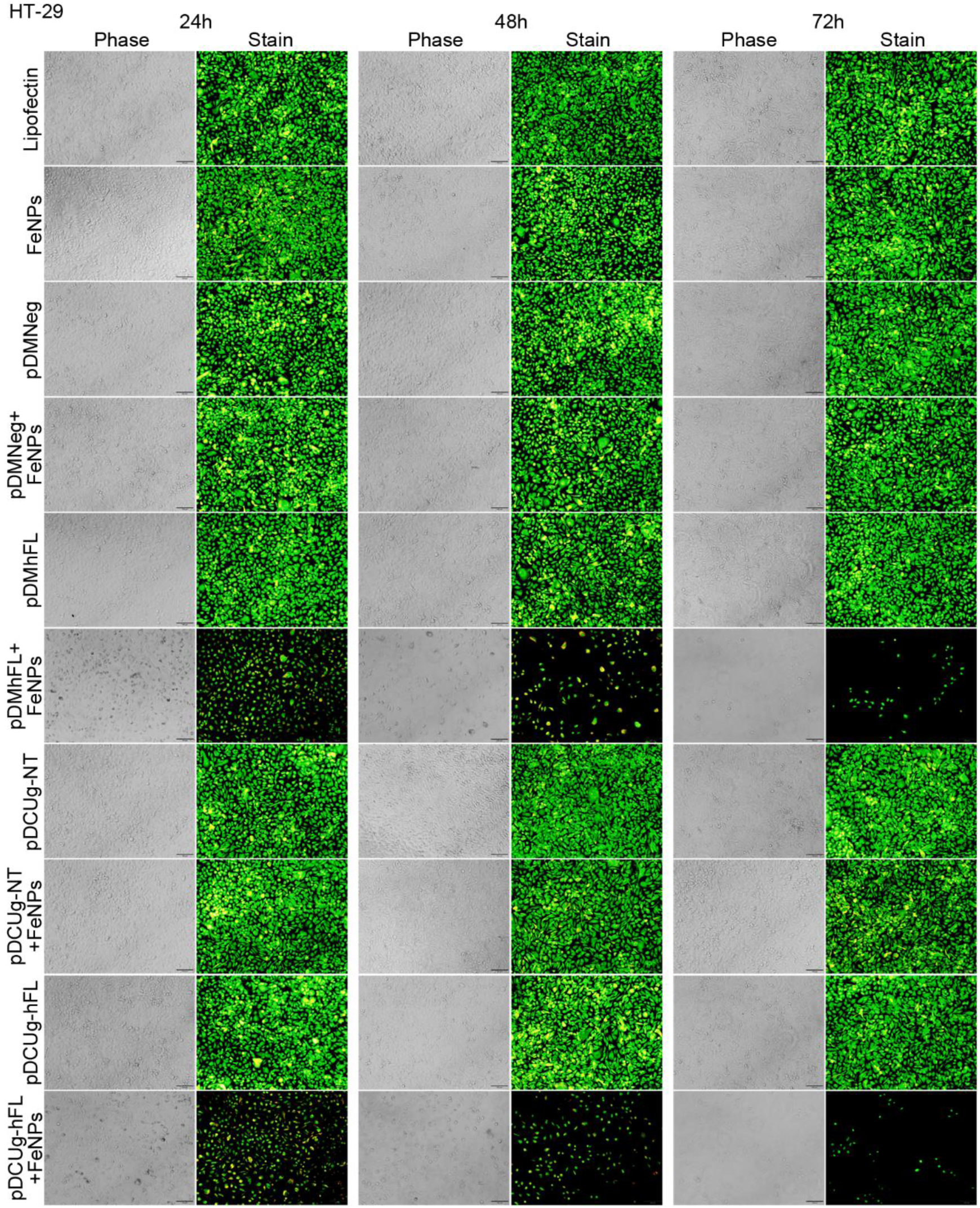
Treatment of HT-29 cells with gene interfering vectors and FeNPs. Cells were transfected by various plasmids, cultured for 24 h and then incubated with or without 50 μg/mL of FeNPs, and cells were cultured for another 72 h. At 24 h, 48 h and 72 h post FeNPs administration, cells were stained with acridine orange and ethidium bromide and imaged under a fluorescence microscope. pDMNeg/pDMhFL, plasmids expressing miRNAs under the control of DMP targeting no or human FPN and Lcn2 transcripts; pDCUg-NT/pDCUg-hFL, plasmids expressing Cas13a under the control of DMP and gRNAs under the control of U6 promoter targeting no or human FPN and Lcn2 transcripts.

**Fig.S12.**
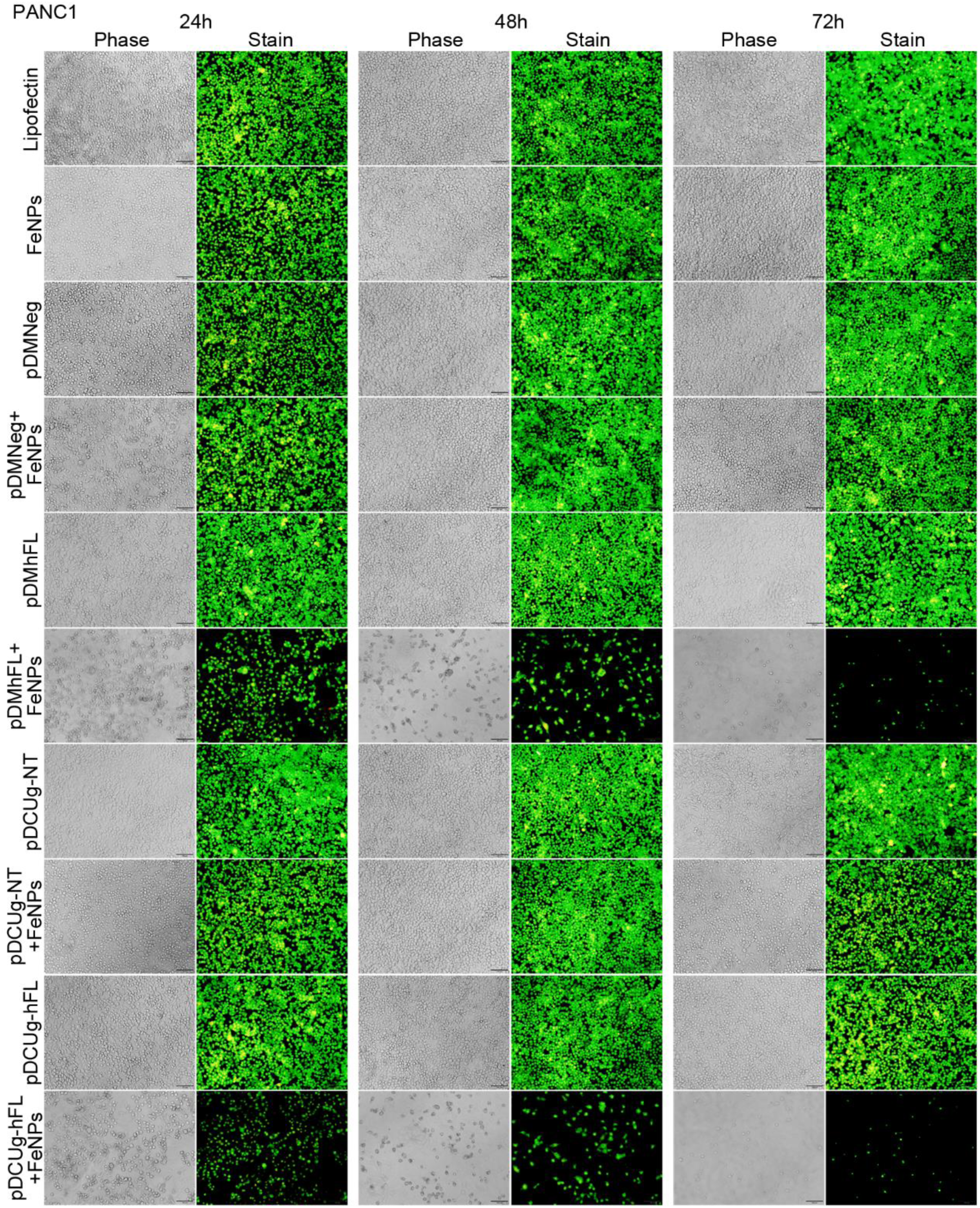
Treatment of PANC1 cells with gene interfering vectors and FeNPs. Cells were transfected by various plasmids, cultured for 24 h and then incubated with or without 50 μg/mL of FeNPs, and cells were cultured for another 72 h. At 24 h, 48 h and 72 h post FeNPs administration, cells were stained with acridine orange and ethidium bromide and imaged under a fluorescence microscope. pDMNeg/pDMhFL, plasmids expressing miRNAs under the control of DMP targeting no or human FPN and Lcn2 transcripts; pDCUg-NT/pDCUg-hFL, plasmids expressing Cas13a under the control of DMP and gRNAs under the control of U6 promoter targeting no or human FPN and Lcn2 transcripts.

**Fig.S13.**
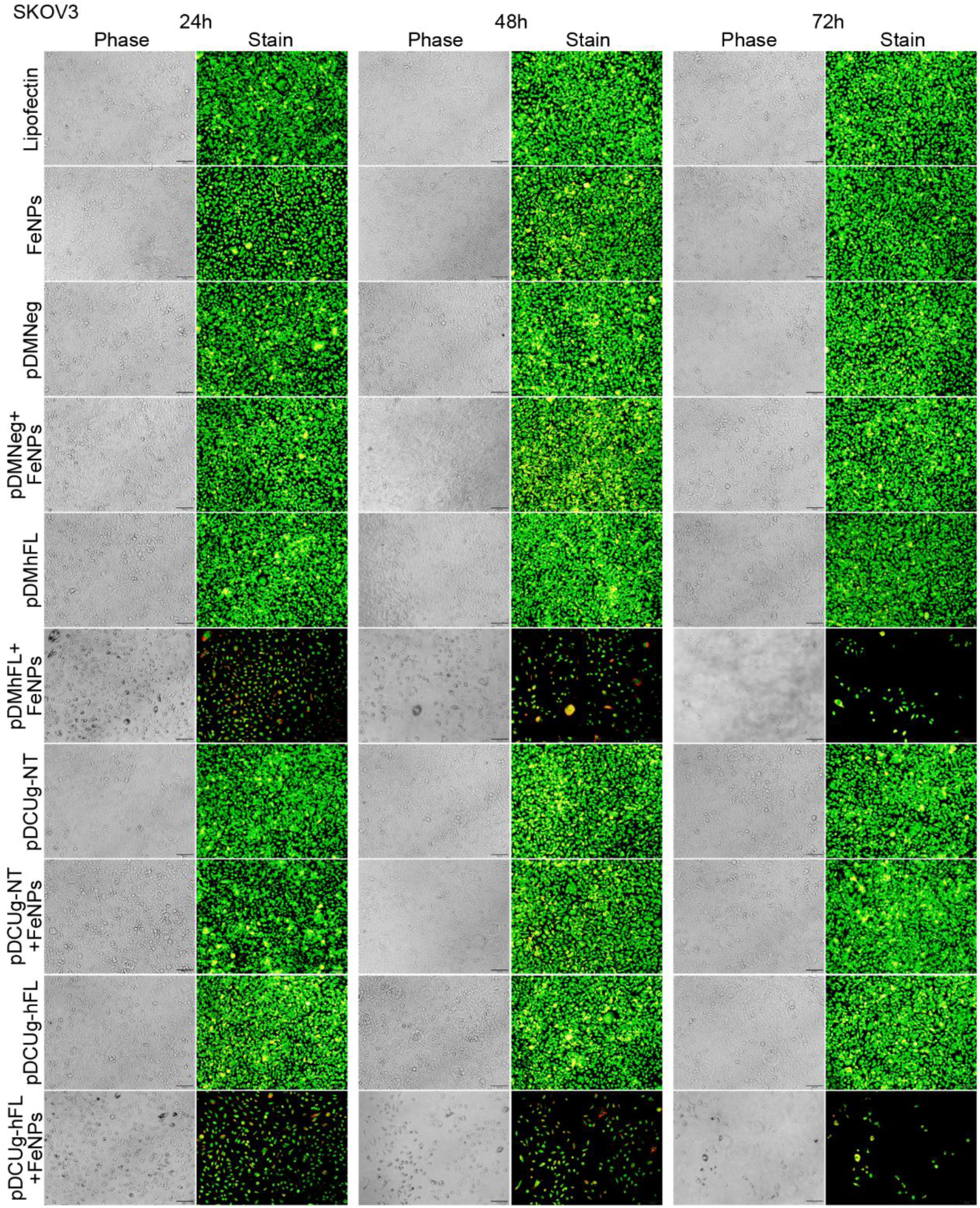
Treatment of SKOV3 cells with gene interfering vectors and FeNPs. Cells were transfected by various plasmids, cultured for 24 h and then incubated with or without 50 μg/mL of FeNPs, and cells were cultured for another 72 h. At 24 h, 48 h and 72 h post FeNPs administration, cells were stained with acridine orange and ethidium bromide and imaged under a fluorescence microscope. pDMNeg/pDMhFL, plasmids expressing miRNAs under the control of DMP targeting no or human FPN and Lcn2 transcripts; pDCUg-NT/pDCUg-hFL, plasmids expressing Cas13a under the control of DMP and gRNAs under the control of U6 promoter targeting no or human FPN and Lcn2 transcripts.

**Fig.S14.**
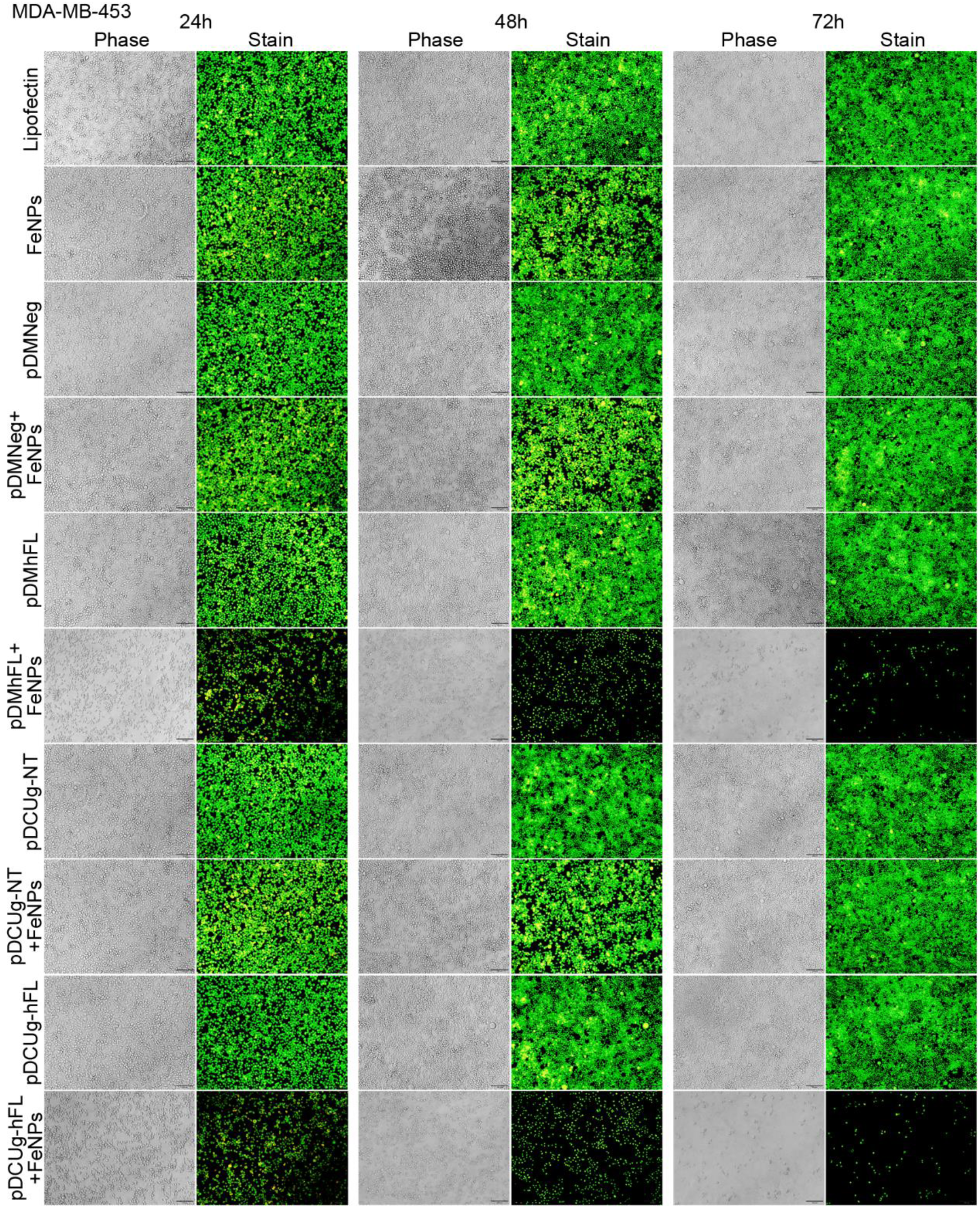
Treatment of MDA-MB-453 cells with gene interfering vectors and FeNPs. Cells were transfected by various plasmids, cultured for 24 h and then incubated with or without 50 μg/mL of FeNPs, and cells were cultured for another 72 h. At 24 h, 48 h and 72 h post FeNPs administration, cells were stained with acridine orange and ethidium bromide and imaged under a fluorescence microscope. pDMNeg/pDMhFL, plasmids expressing miRNAs under the control of DMP targeting no or human FPN and Lcn2 transcripts; pDCUg-NT/pDCUg-hFL, plasmids expressing Cas13a under the control of DMP and gRNAs under the control of U6 promoter targeting no or human FPN and Lcn2 transcripts.

**Fig.S15.**
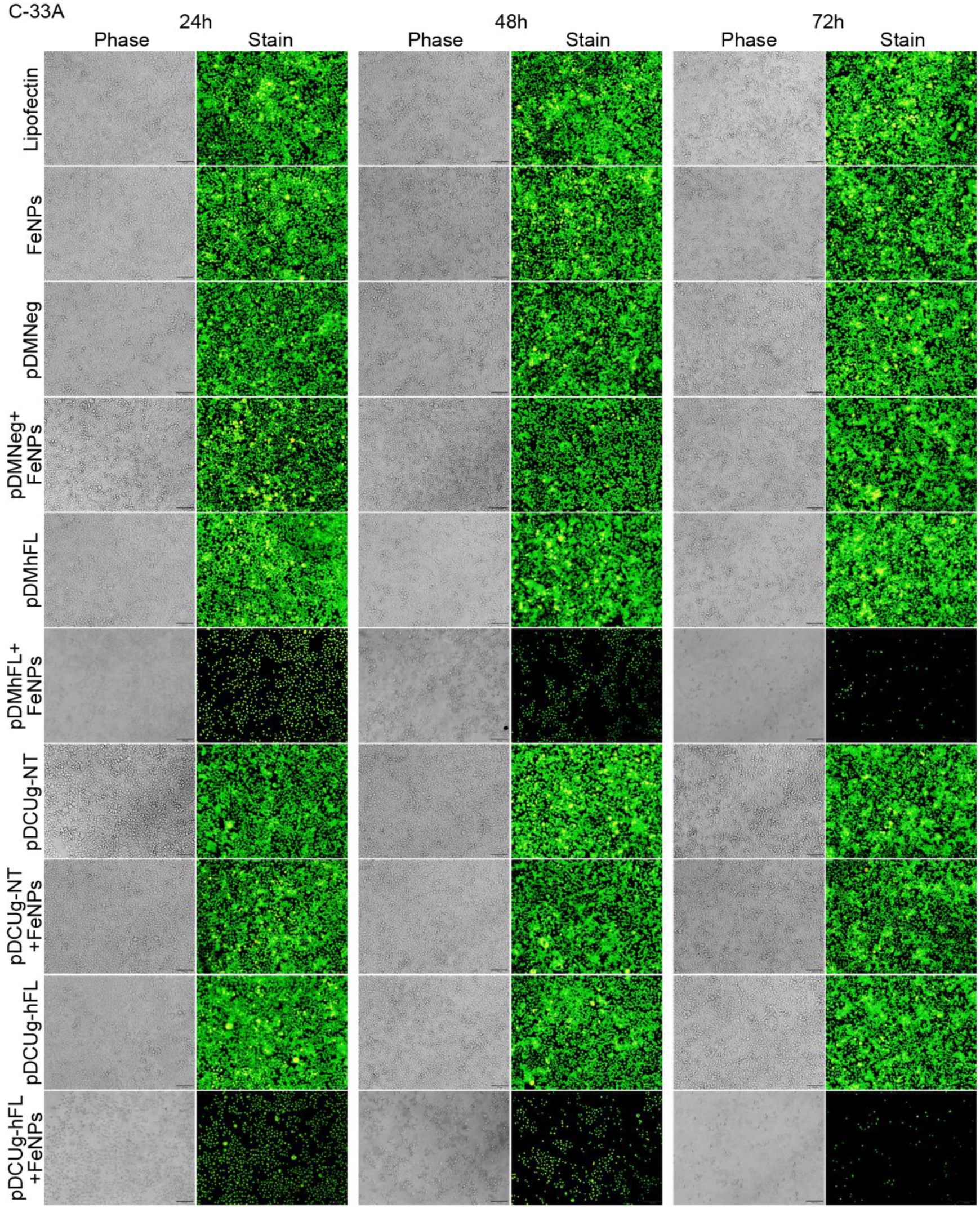
Treatment of C-33A cells with gene interfering vectors and FeNPs. Cells were transfected by various plasmids, cultured for 24 h and then incubated with or without 50 μg/mL of FeNPs, and cells were cultured for another 72 h. At 24 h, 48 h and 72 h post FeNPs administration, cells were stained with acridine orange and ethidium bromide and imaged under a fluorescence microscope. pDMNeg/pDMhFL, plasmids expressing miRNAs under the control of DMP targeting no or human FPN and Lcn2 transcripts; pDCUg-NT/pDCUg-hFL, plasmids expressing Cas13a under the control of DMP and gRNAs under the control of U6 promoter targeting no or human FPN and Lcn2 transcripts.

**Fig.S16.**
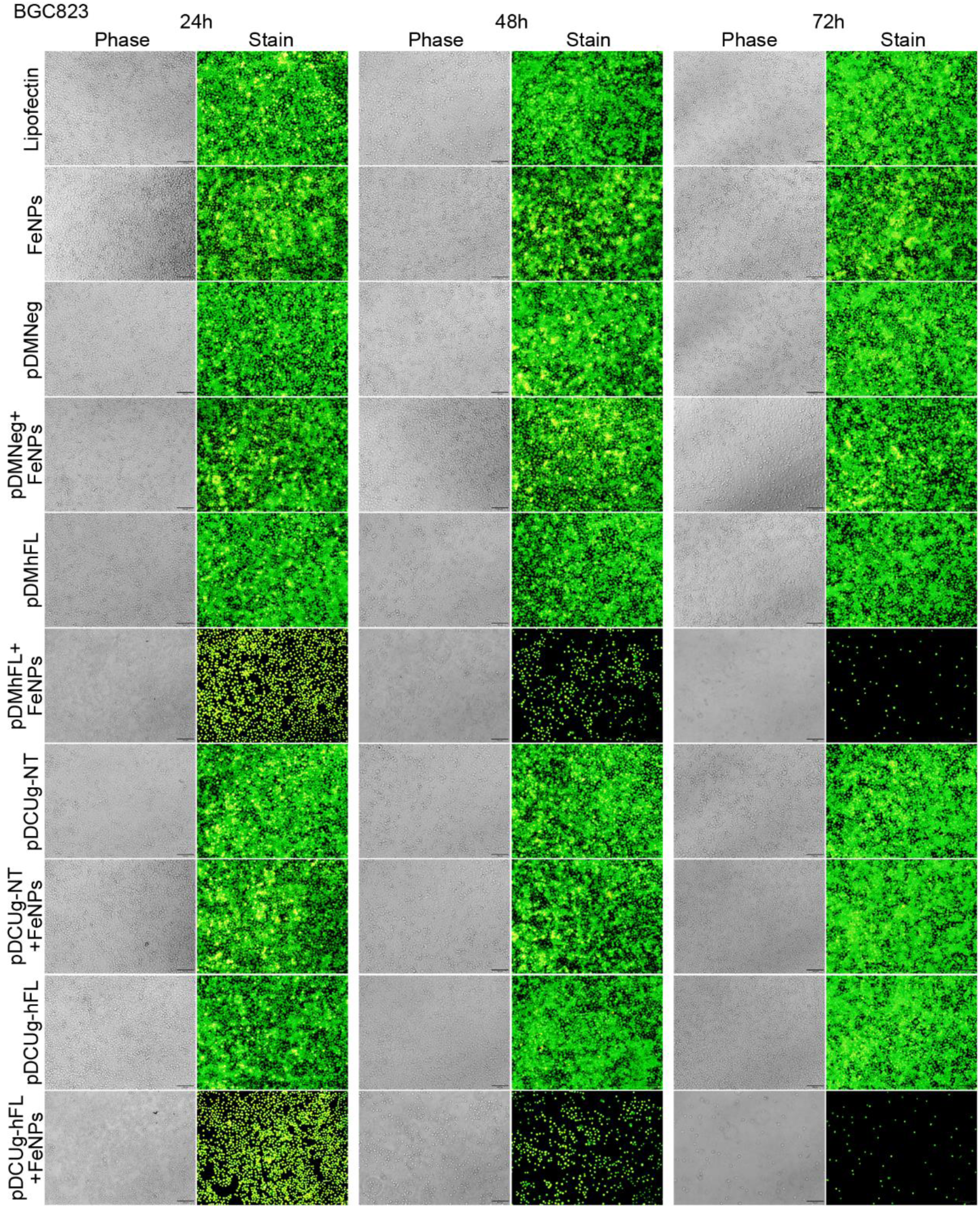
Treatment of BGC823 cells with gene interfering vectors and FeNPs. Cells were transfected by various plasmids, cultured for 24 h and then incubated with or without 50 μg/mL of FeNPs, and cells were cultured for another 72 h. At 24 h, 48 h and 72 h post FeNPs administration, cells were stained with acridine orange and ethidium bromide and imaged under a fluorescence microscope. pDMNeg/pDMhFL, plasmids expressing miRNAs under the control of DMP targeting no or human FPN and Lcn2 transcripts; pDCUg-NT/pDCUg-hFL, plasmids expressing Cas13a under the control of DMP and gRNAs under the control of U6 promoter targeting no or human FPN and Lcn2 transcripts.

**Fig.S17.**
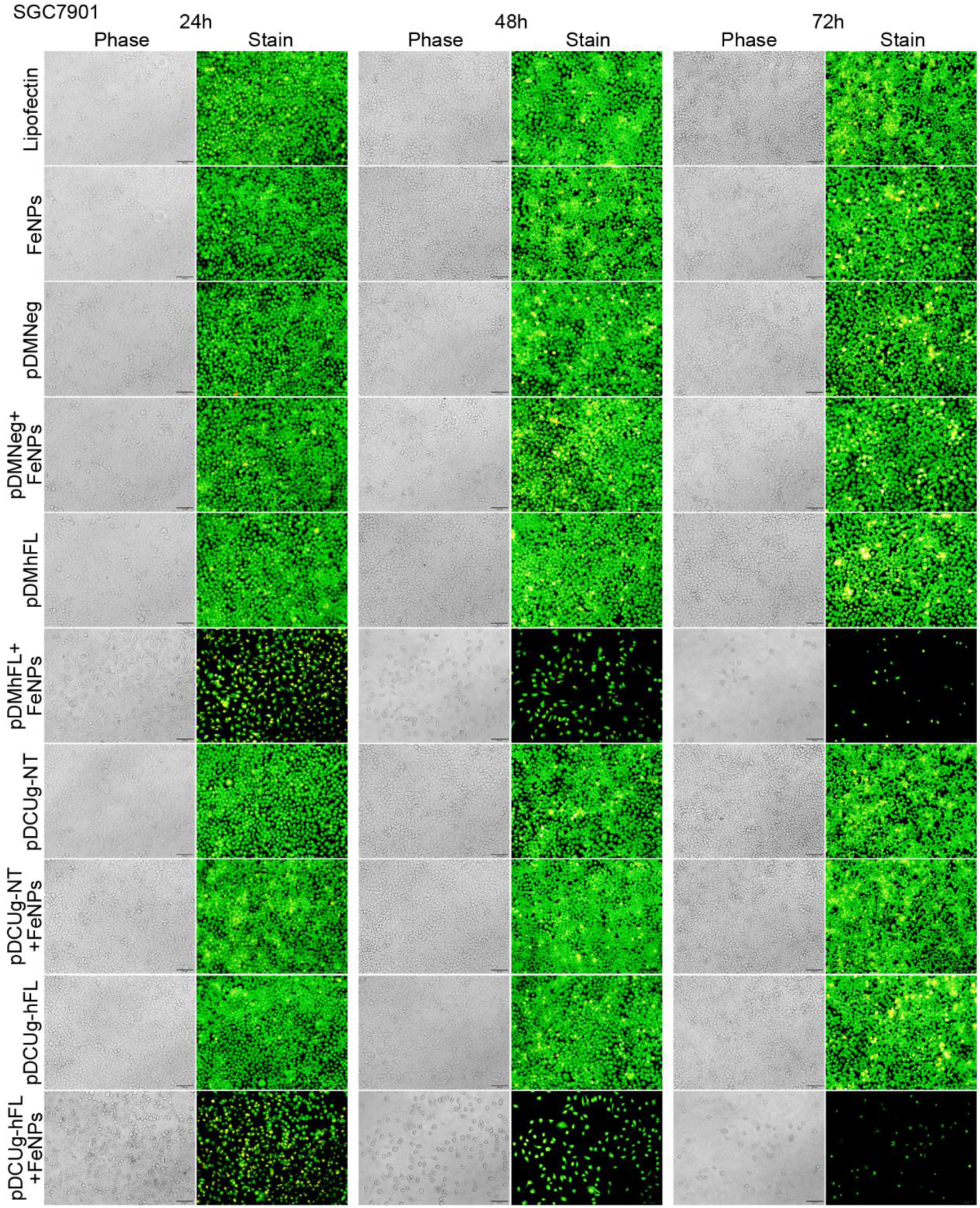
Treatment of SGC7901 cells with gene interfering vectors and FeNPs. Cells were transfected by various plasmids, cultured for 24 h and then incubated with or without 50 μg/mL of FeNPs, and cells were cultured for another 72 h. At 24 h, 48 h and 72 h post FeNPs administration, cells were stained with acridine orange and ethidium bromide and imaged under a fluorescence microscope. pDMNeg/pDMhFL, plasmids expressing miRNAs under the control of DMP targeting no or human FPN and Lcn2 transcripts; pDCUg-NT/pDCUg-hFL, plasmids expressing Cas13a under the control of DMP and gRNAs under the control of U6 promoter targeting no or human FPN and Lcn2 transcripts.

**Fig.S18.**
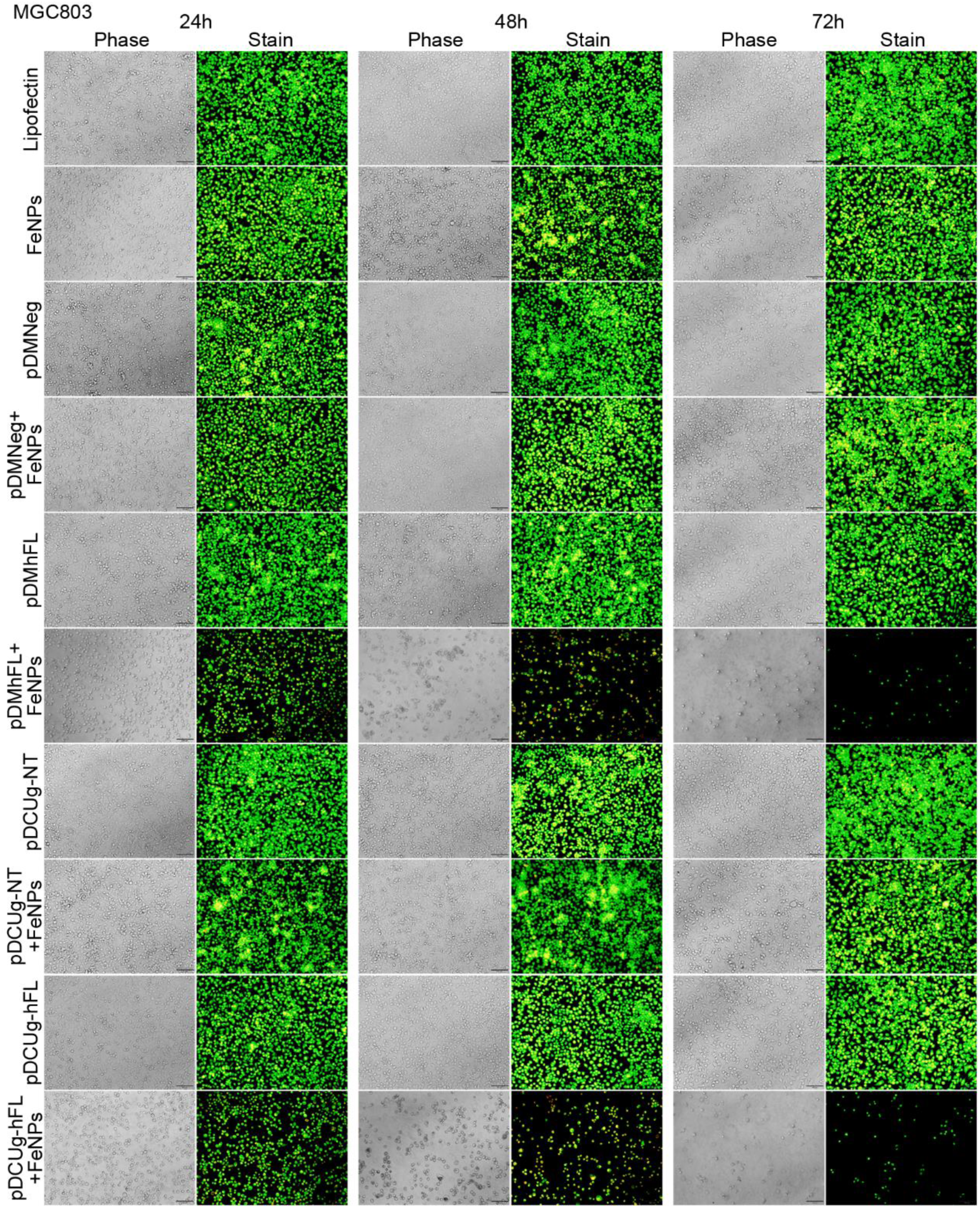
Treatment of MGC-803 cells with gene interfering vectors and FeNPs. Cells were transfected by various plasmids, cultured for 24 h and then incubated with or without 50 μg/mL of FeNPs, and cells were cultured for another 72 h. At 24 h, 48 h and 72 h post FeNPs administration, cells were stained with acridine orange and ethidium bromide and imaged under a fluorescence microscope. pDMNeg/pDMhFL, plasmids expressing miRNAs under the control of DMP targeting no or human FPN and Lcn2 transcripts; pDCUg-NT/pDCUg-hFL, plasmids expressing Cas13a under the control of DMP and gRNAs under the control of U6 promoter targeting no or human FPN and Lcn2 transcripts.

**Fig.S19.**
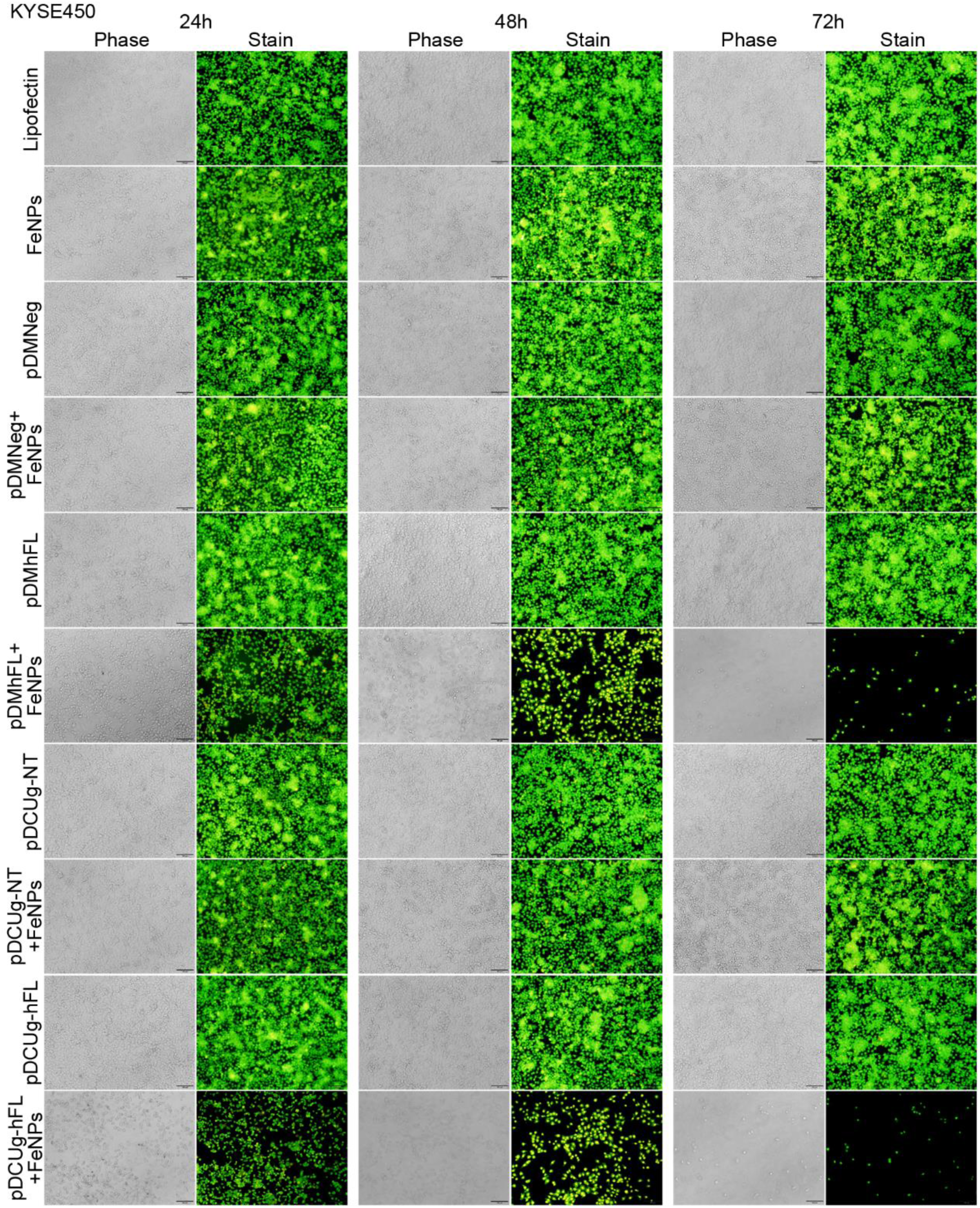
Treatment of KYSE450 cells with gene interfering vectors and FeNPs. Cells were transfected by various plasmids, cultured for 24 h and then incubated with or without 50 μg/mL of FeNPs, and cells were cultured for another 72 h. At 24 h, 48 h and 72 h post FeNPs administration, cells were stained with acridine orange and ethidium bromide and imaged under a fluorescence microscope. pDMNeg/pDMhFL, plasmids expressing miRNAs under the control of DMP targeting no or human FPN and Lcn2 transcripts; pDCUg-NT/pDCUg-hFL, plasmids expressing Cas13a under the control of DMP and gRNAs under the control of U6 promoter targeting no or human FPN and Lcn2 transcripts.

**Fig.S20.**
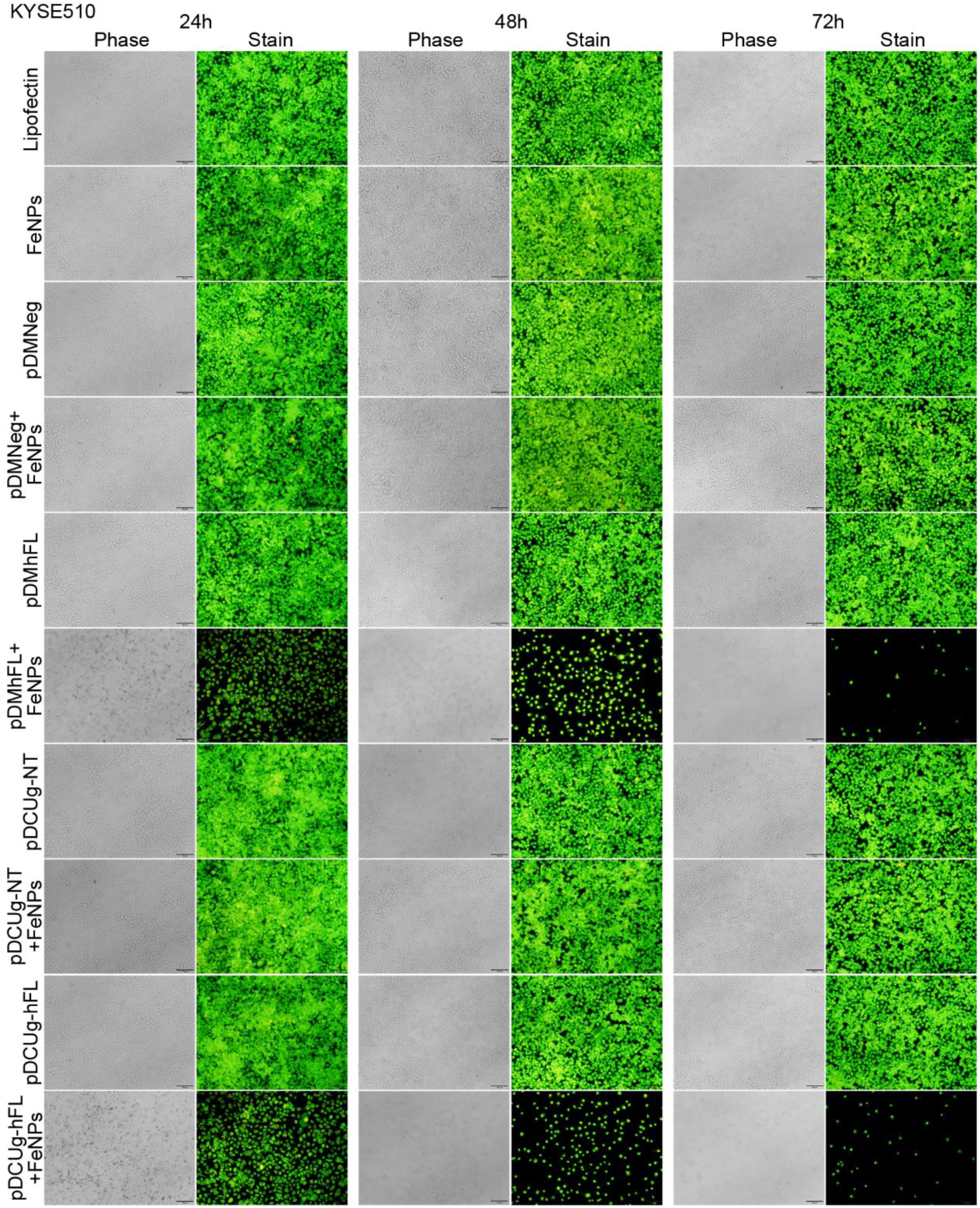
Treatment of KYSE510 cells with gene interfering vectors and FeNPs. Cells were transfected by various plasmids, cultured for 24 h and then incubated with or without 50 μg/mL of FeNPs, and cells were cultured for another 72 h. At 24 h, 48 h and 72 h post FeNPs administration, cells were stained with acridine orange and ethidium bromide and imaged under a fluorescence microscope. pDMNeg/pDMhFL, plasmids expressing miRNAs under the control of DMP targeting no or human FPN and Lcn2 transcripts; pDCUg-NT/pDCUg-hFL, plasmids expressing Cas13a under the control of DMP and gRNAs under the control of U6 promoter targeting no or human FPN and Lcn2 transcripts.

**Fig.S21.**
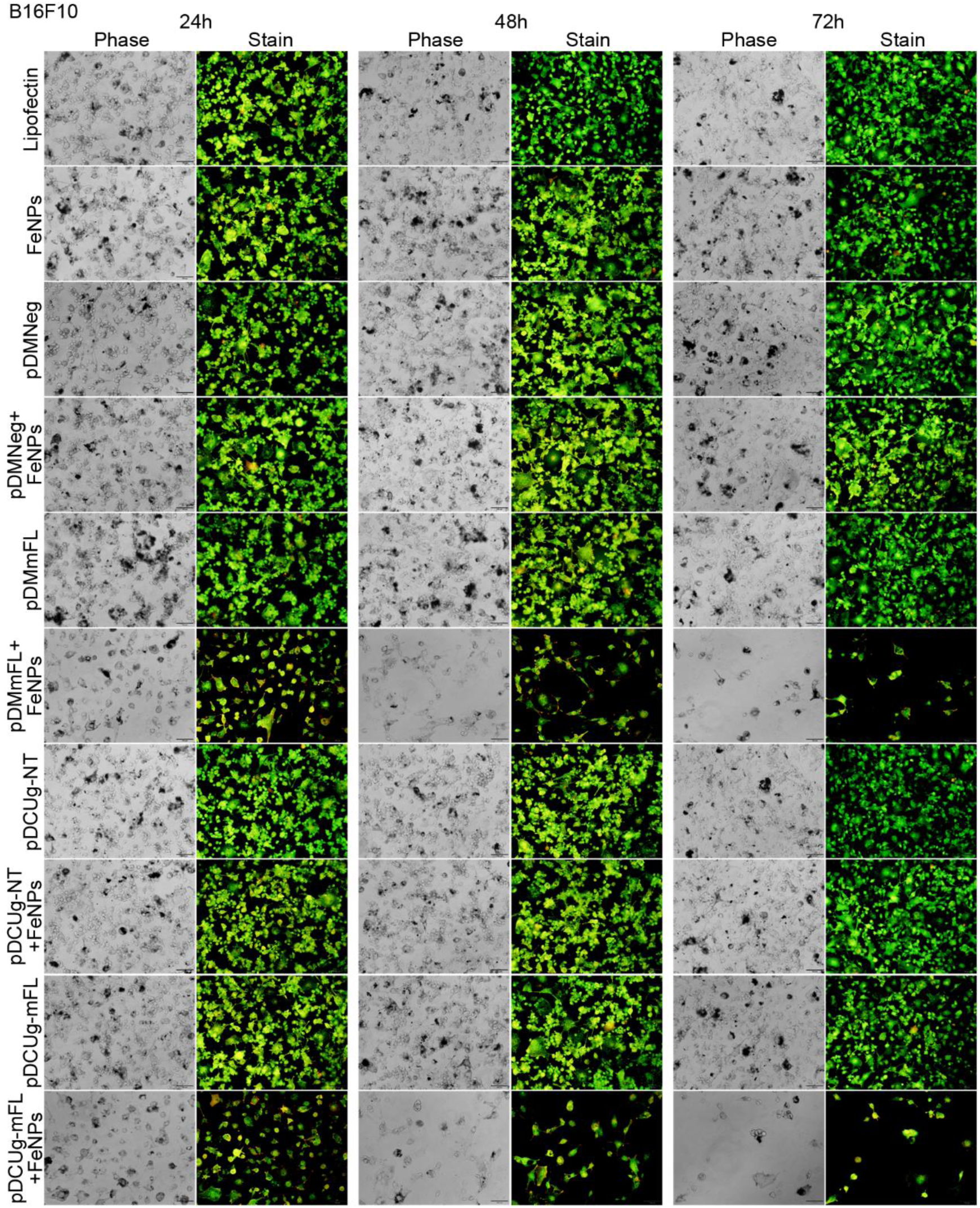
Treatment of B16F10 cells with gene interfering vectors and FeNPs. Cells were transfected by various plasmids, cultured for 24 h and then incubated with or without 50 μg/mL of FeNPs, and cells were cultured for another 72 h. At 24 h, 48 h and 72 h post FeNPs administration, cells were stained with acridine orange and ethidium bromide and imaged under a fluorescence microscope. pDMNeg/pDMmFL, plasmids expressing miRNAs under the control of DMP targeting no or mouse FPN and Lcn2 transcripts; pDCUg-NT/pDCUg-mFL, plasmids expressing Cas13a under the control of DMP and gRNAs under the control of U6 promoter targeting no or mouse FPN and Lcn2 transcripts.

**Fig.S22.**
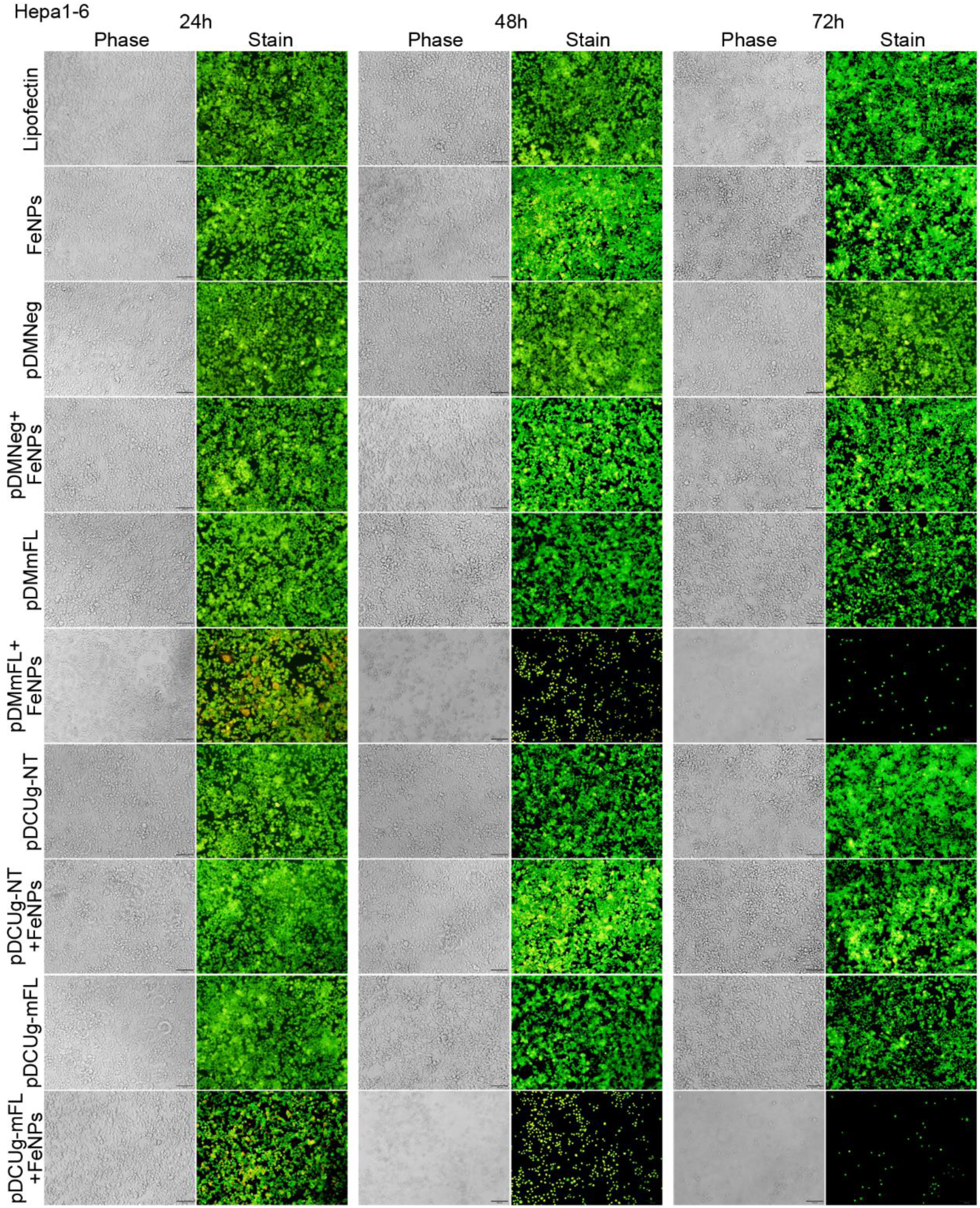
Treatment of Hepa1-6 cells with gene interfering vectors and FeNPs. Cells were transfected by various plasmids, cultured for 24 h and then incubated with or without 50 μg/mL of FeNPs, and cells were cultured for another 72 h. At 24 h, 48 h and 72 h post FeNPs administration, cells were stained with acridine orange and ethidium bromide and imaged under a fluorescence microscope. pDMNeg/pDMmFL, plasmids expressing miRNAs under the control of DMP targeting no or mouse FPN and Lcn2 transcripts; pDCUg-NT/pDCUg-mFL, plasmids expressing Cas13a under the control of DMP and gRNAs under the control of U6 promoter targeting no or mouse FPN and Lcn2 transcripts.

**Fig.S23.**
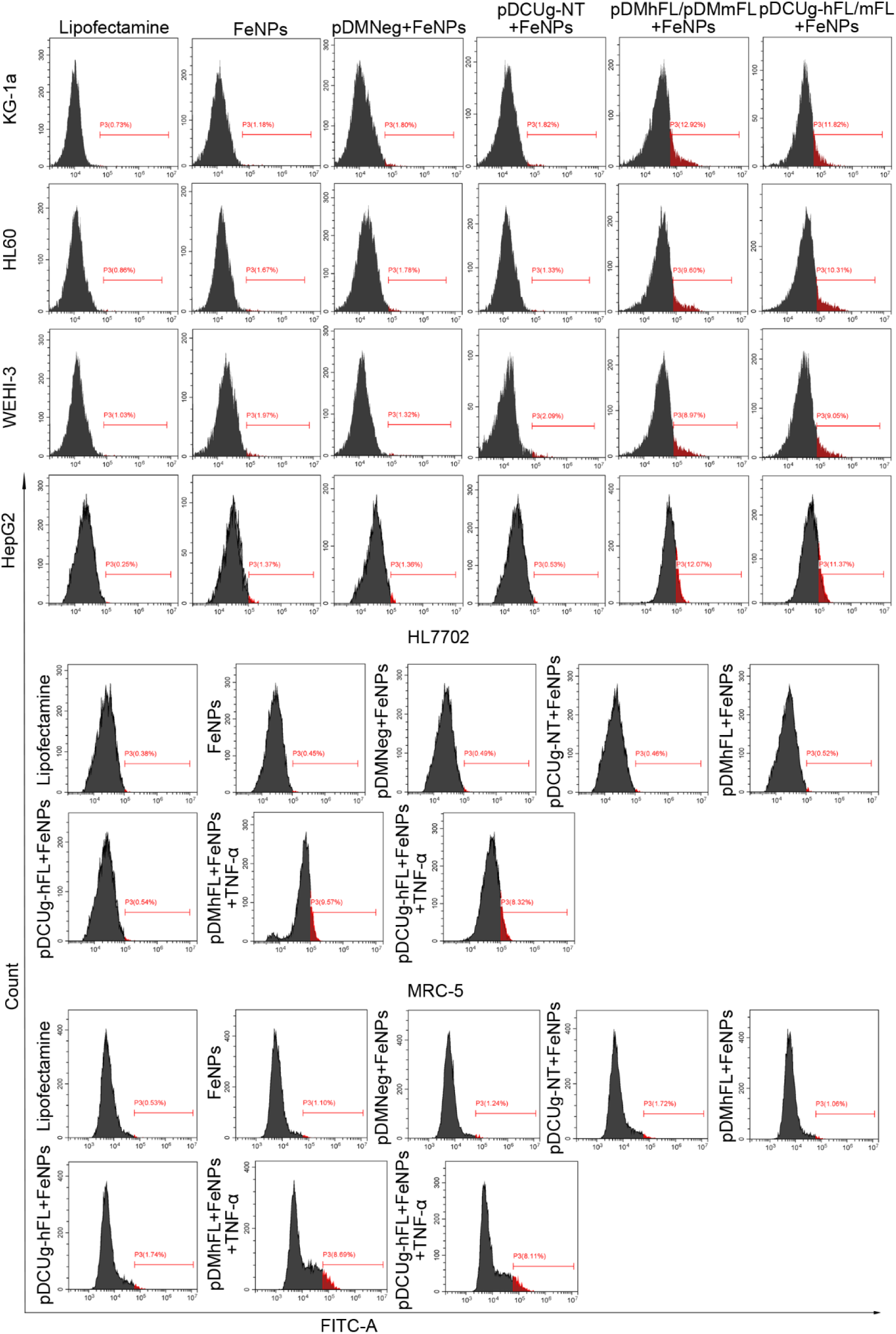
Flow cytometry analysis of ROS levels of cells treated with gene interfering vectors and FeNPs. Cells were transfected by various plasmids, cultured for 24 h and then incubated with or without 50 μg/mL of FeNPs, and cells were cultured for another 48h. For HL7702 and MRC-5, cells were first induced with or without TNF-α (10 ng/mL) for 1h before FeNPs treatment. Cells were collected at 48h post FeNPs administration and stained with DCFH-DA using Reactive Oxygen Species Assay Kit. ROS changes indicated by fluorescence shift were analyzed by Flow Cytometer. pDMNeg/pDMhFL/pDMmFL, plasmids expressing miRNAs under the control of DMP targeting no or human FPN and Lcn2, or mouse FPN and Lcn2 transcripts; pDCUg-NT/pDCUg-hFL/pDCUg-mFL, plasmids expressing Cas13a under the control of DMP and gRNAs under the control of U6 promoter targeting no or human FPN and Lcn2, or mouse FPN and Lcn2 transcripts.

**Fig.S24.**
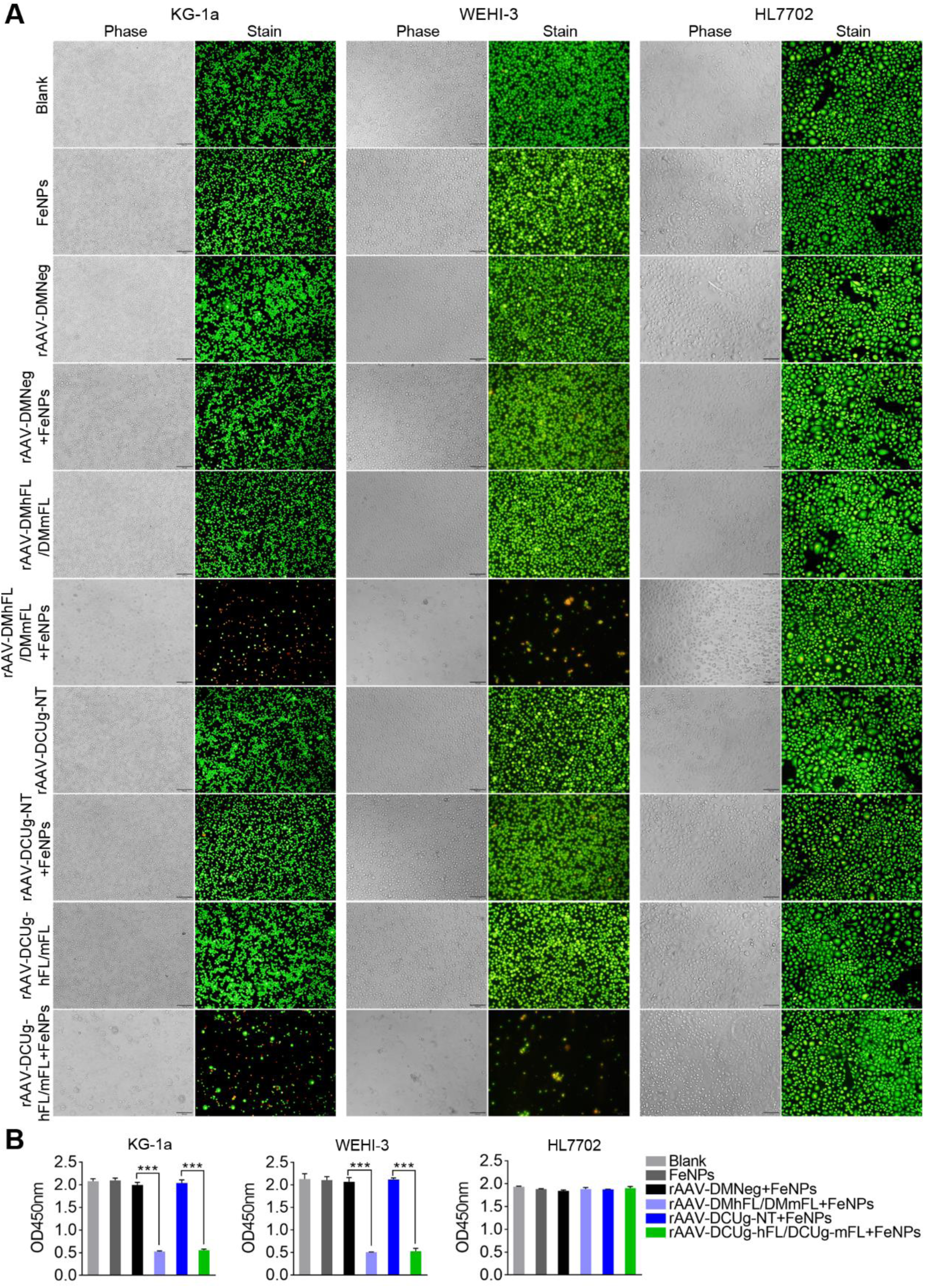
Evaluation of rAAVs in vitro. KG-1a, WEHI-3 and HL7702 cells were seeded into 24-well plates (1×10^5^ cells/well) and cultivated for 12 h. Cells were then transfected with the various viruses at the dose of 1×10^5^ vg per cell. The transfected cells were cultured for 24 h and then incubated with or without 50 μg/mL FeNPs. Cells were cultured for another 72 h. All cells were stained with acridine orange and ethidium bromide and imaged by optical microscope, and cell viability was evaluated using a CCK-8 assay. (A) Representative images of cells at 72 h post FeNPs administration. (B) The cells viability at 72 h post transfection. All values are mean± s.e.m. with n= 3. *, p < 0.05; **, p < 0.01; ***, p < 0.001. rAAV-DMNeg/rAAV-DMhFL/rAAV-DMmFL, viruses expressing miRNAs under the control of DMP targeting no or human FPN and Lcn2, or mouse FPN and Lcn2 transcripts; rAAV-DCUg-NT/rAAV-DCUg-hFL/rAAV-DCUg-mFL, viruses expressing Cas13a under the control of DMP and gRNAs under the control of U6 promoter targeting no or human FPN and Lcn2, or mouse FPN and Lcn2 transcripts.

**Fig.S25.**
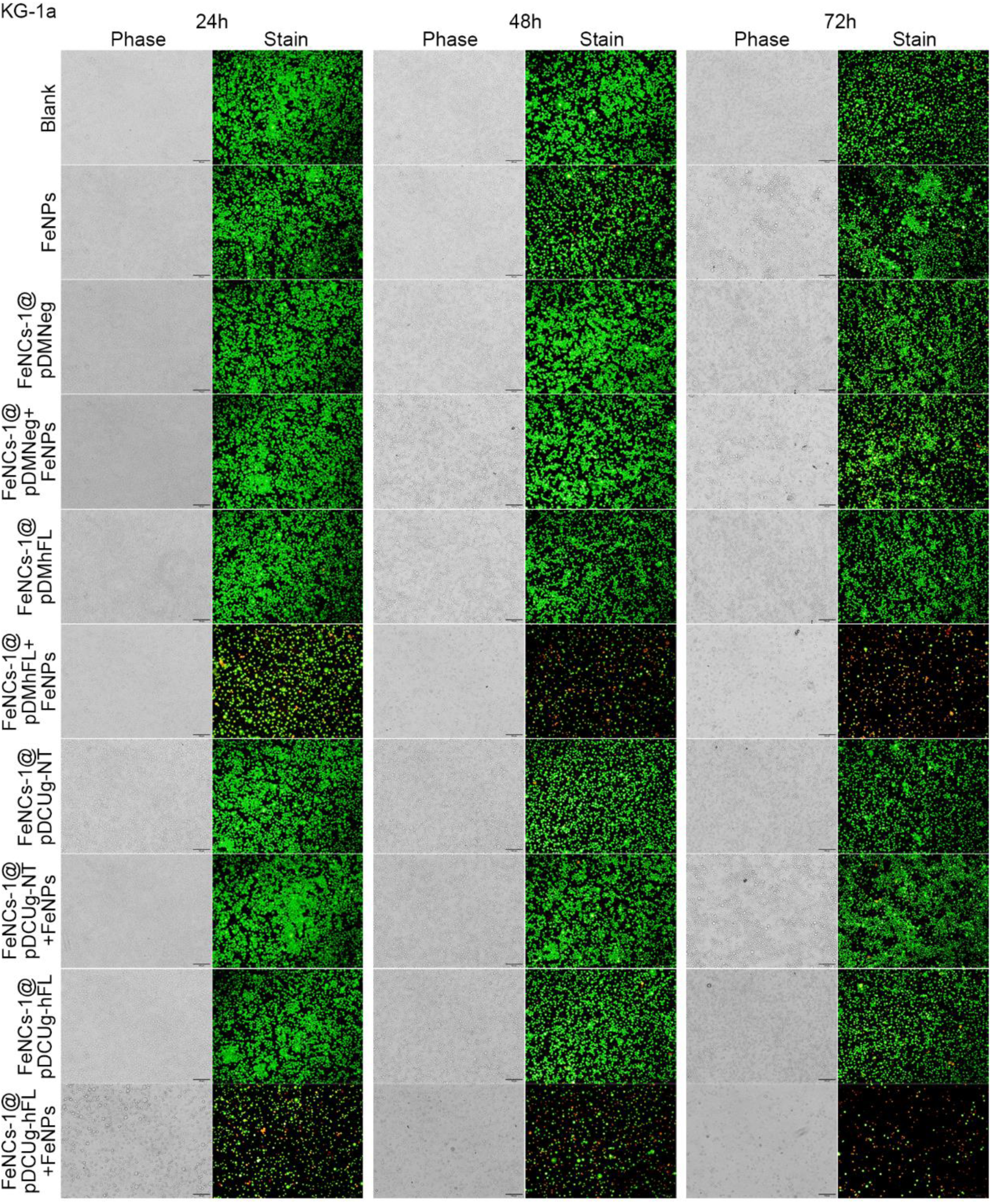
Transfection of KG-1a cells with various plasmids by Fe nanocarriers (FeNCs). Cells (1×10^5^) were seeded into 24-well plates overnight before transfection. Cells were treated with Fe nanocarrier (0.5 μg) and 500 ng of various plasmids according to the manufacturer’s instructions. The transfected cells were cultured for 24 h and then incubated with or without 50 μg/mL of FeNPs, and cells were cultured for another 72 h. At 24 h, 48 h and 72 h post FeNPs administration, cells were stained with acridine orange and ethidium bromide and imaged under a fluorescence microscope.

**Fig.S26.**
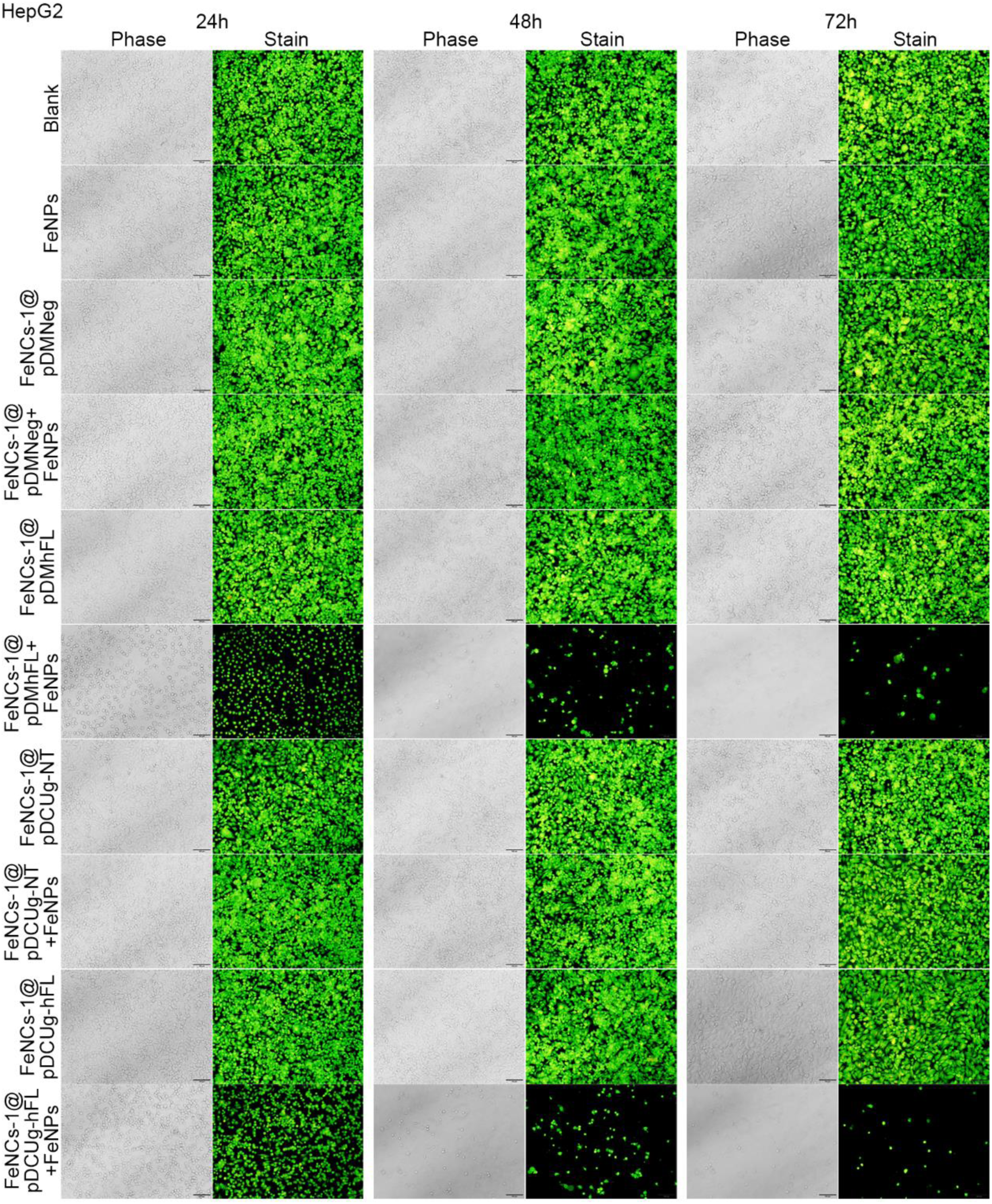
Transfection of HepG2 cells with various plasmids by Fe nanocarriers (FeNCs). Cells (1×10^5^) were seeded into 24-well plates overnight before transfection. Cells were treated with FeNCs (0.5 μg) and 500 ng of various plasmids according to the manufacturer’s instructions. The transfected cells were cultured for 24 h and then incubated with or without 50 μg/mL of FeNPs, and cells were cultured for another 72 h. At 24 h, 48 h and 72 h post FeNPs administration, cells were stained with acridine orange and ethidium bromide and imaged under a fluorescence microscope.

**Fig.S27.**
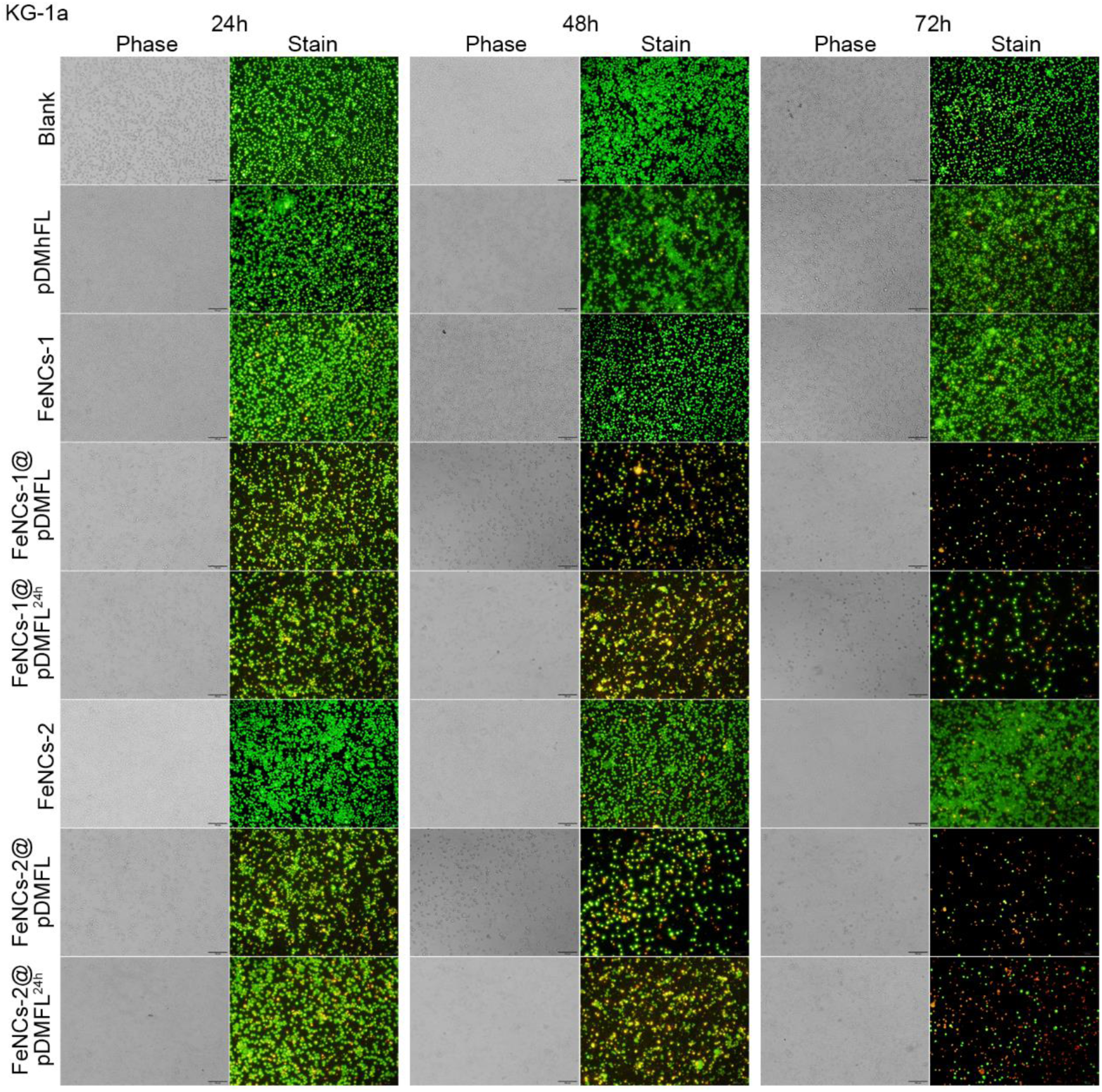
Transfection of KG-1a cells with various plasmids by two Fe nanocarriers (FeNCs). Cells (1×10^5^) were seeded into 24-well plates overnight before transfection. Cells were treated with various plasmids and 50 μg/mL of FeNCs (FeNCs-1 and FeNCs-2), respectively. All cells were cultured for another 72 h. At 24 h, 48 h and 72 h post FeNCs administration, cells were stained with acridine orange and ethidium bromide and imaged under a fluorescence microscope. pDMFL+ FeNCs-1/FeNCs-2, plasmids and FeNCs were mixed and added directly to the cells; pDMFL+ FeNCs-1^24h^/FeNCs-2^24h^, plasmids and FeNCs were mixed and added to the cells after 24 h.

## Supplementary Data

**Table S1.**
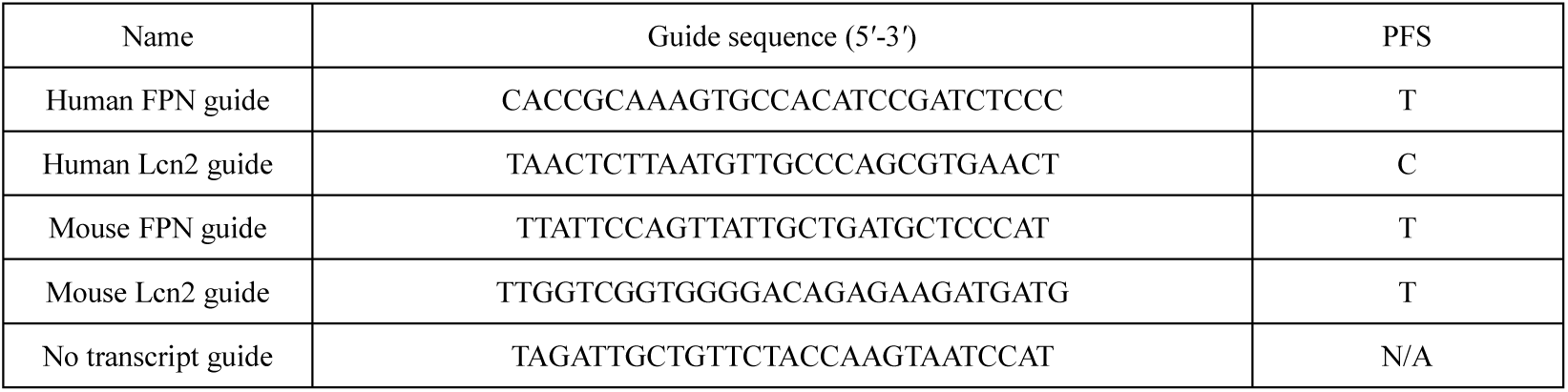
Guide sequences of Cas13a for interference experiments.

**Table S2.**
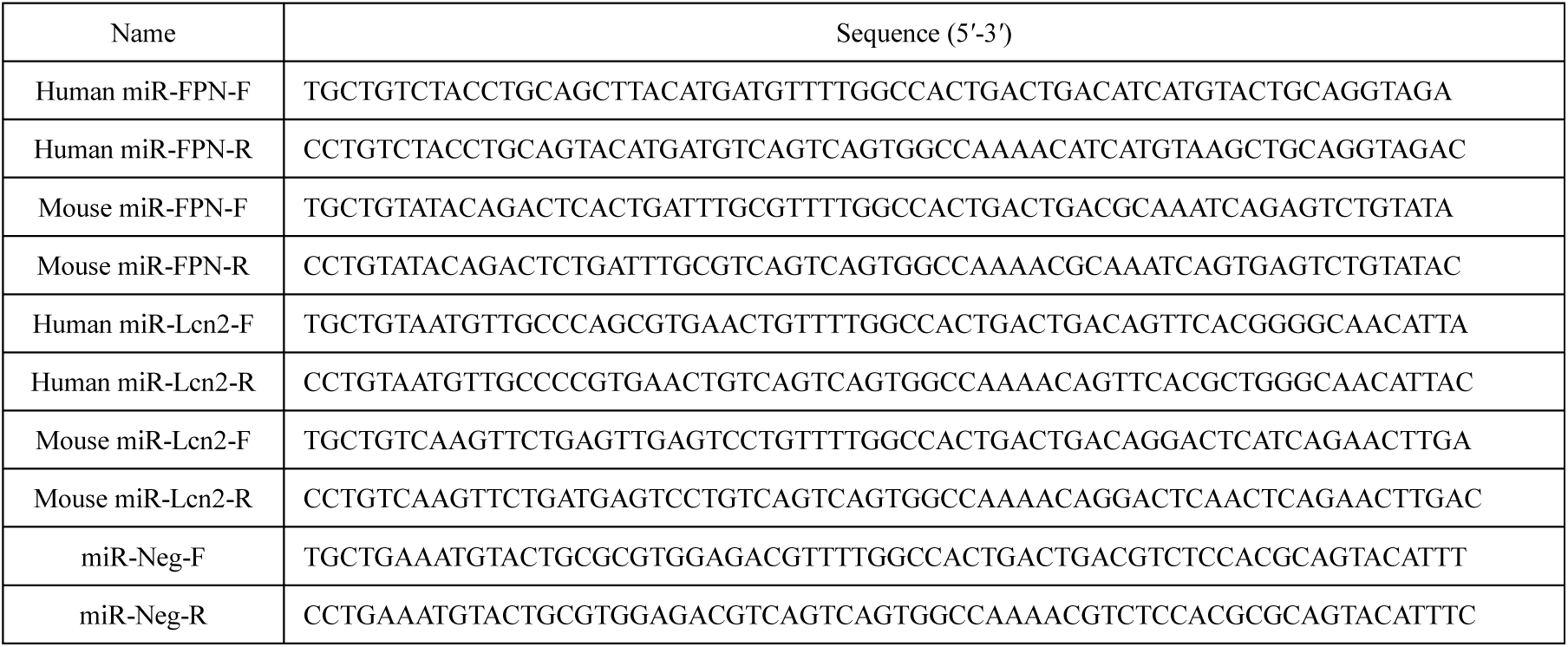
Oligonucleotides sequences used in miRNA interference experiments.

**Table S3.**
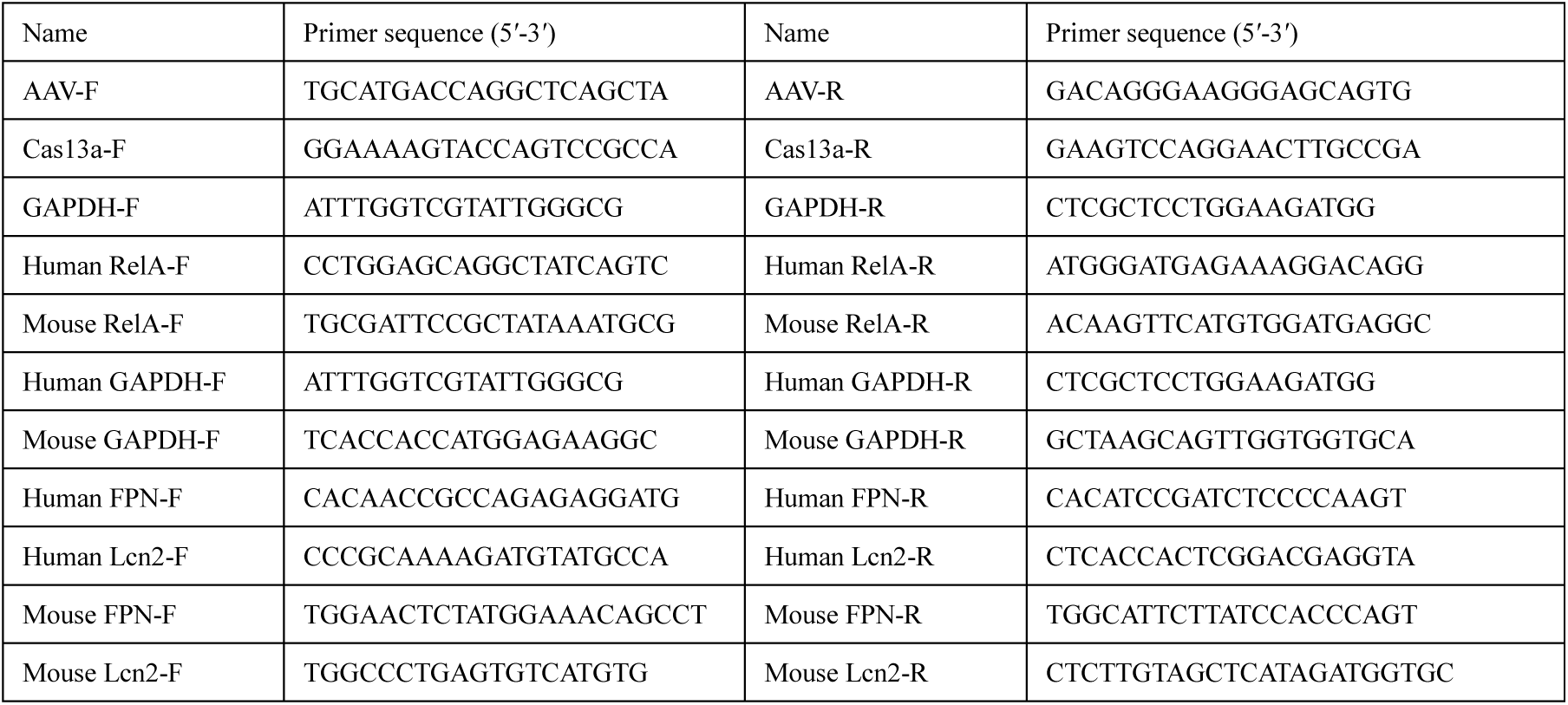
Primer sequences used for qPCR.

## Plasmids and the functional sequences

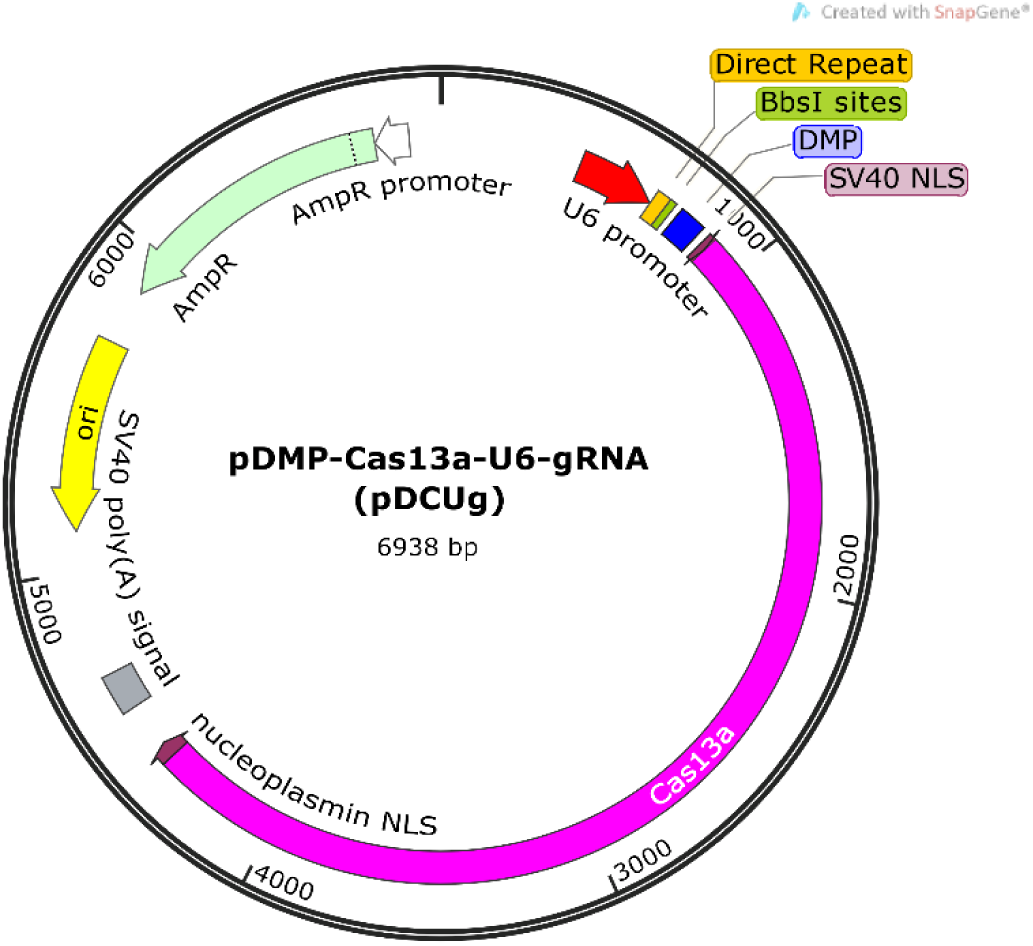

### pDMP-Cas13a-U6-gRNA

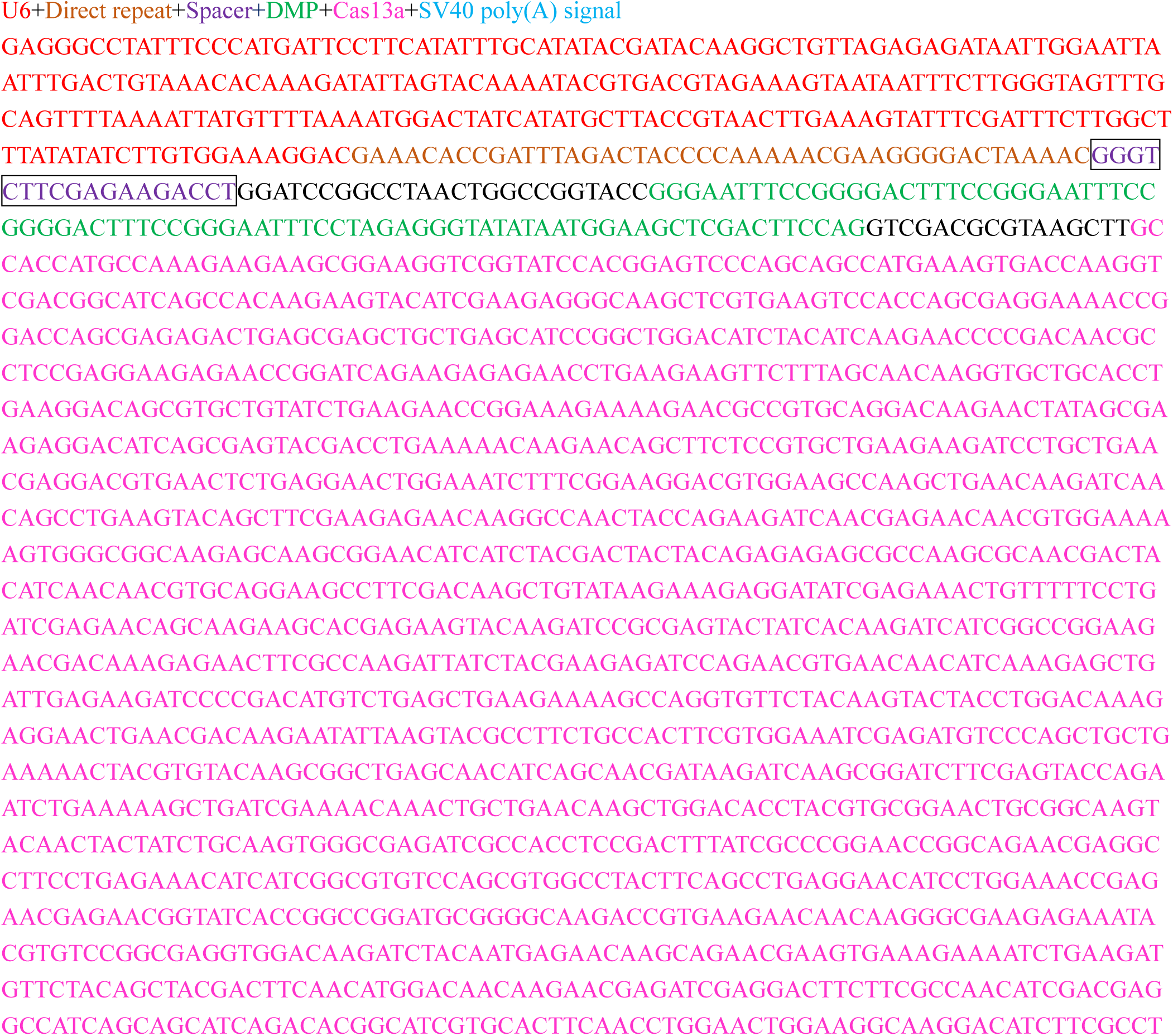

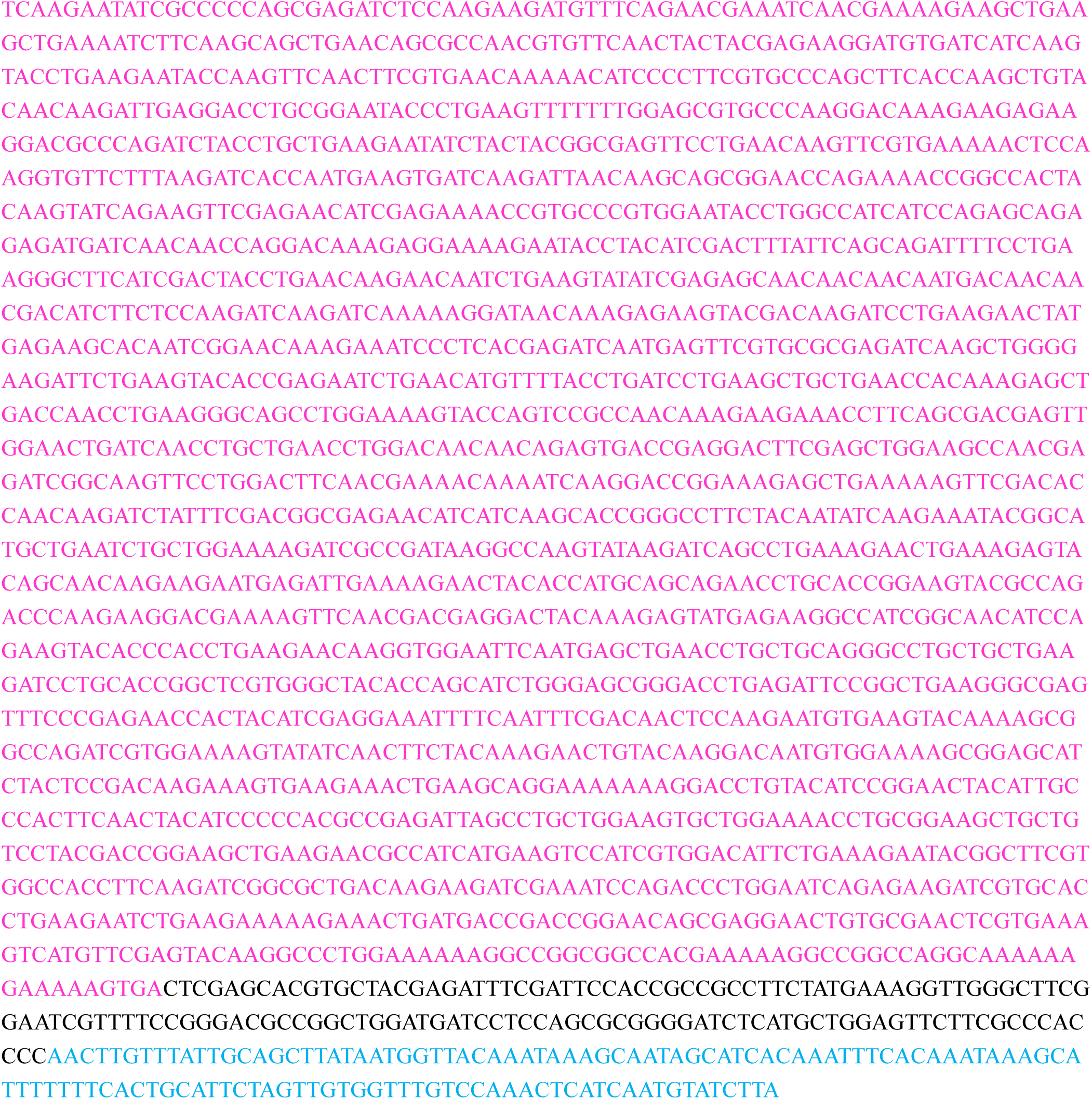

The spacer sequence in above skeleton vector can be replaced by the following sequences for constructing pDMP-Cas13a-U6-gRNAs targeting particular genes:

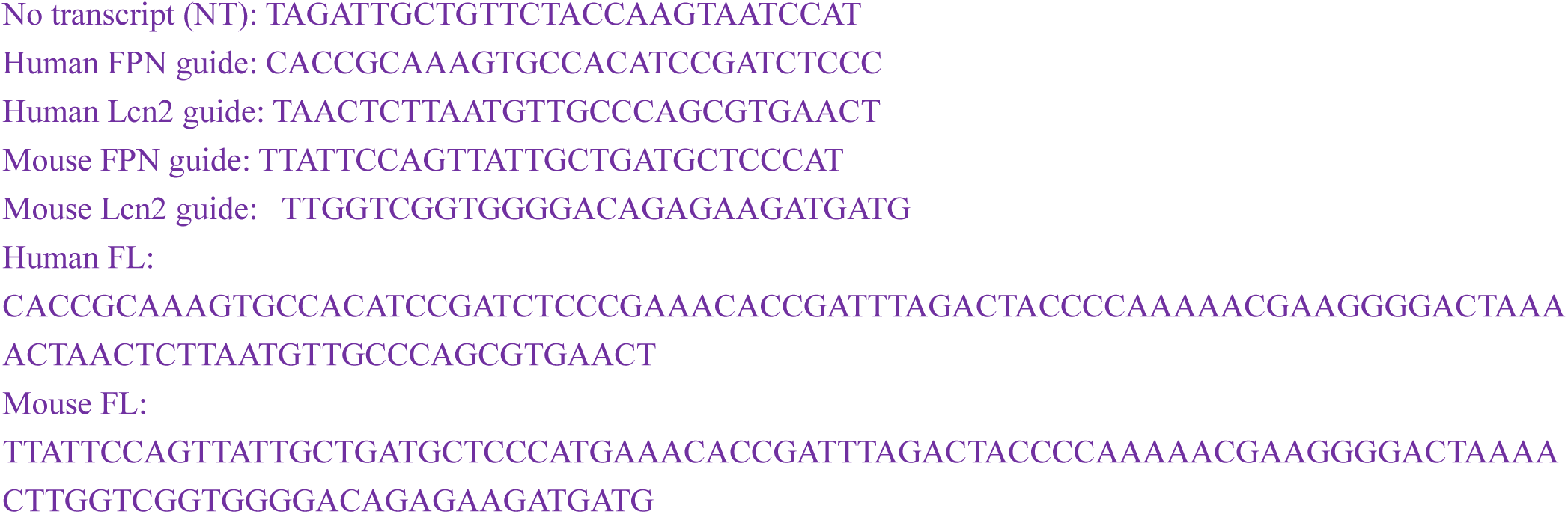

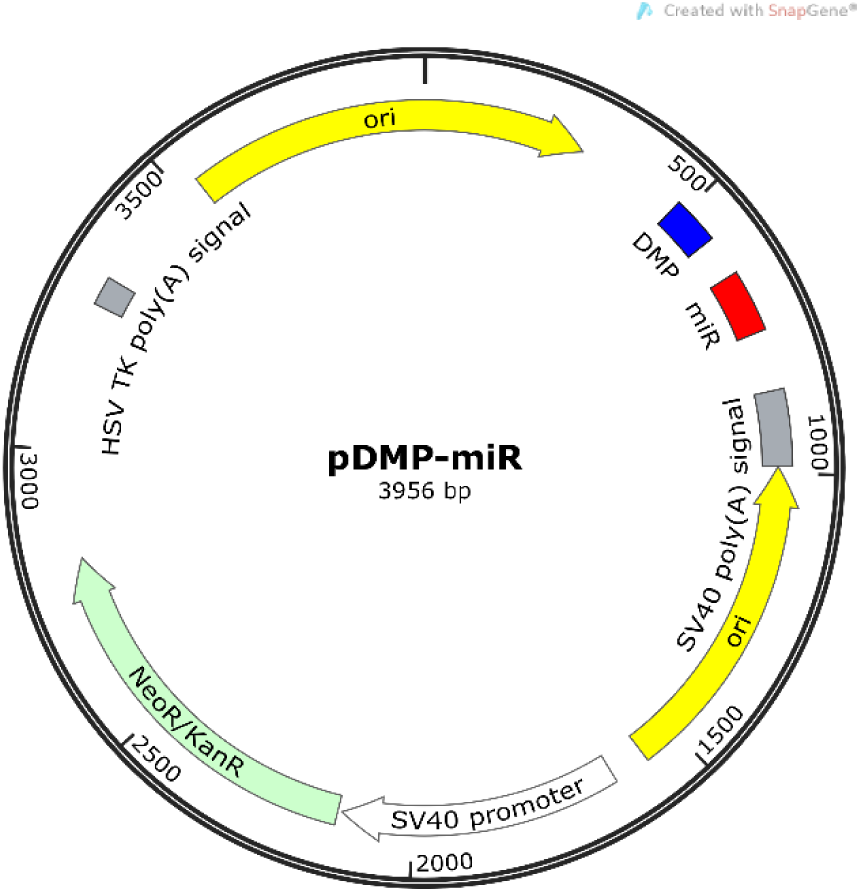

### pDMP-miR

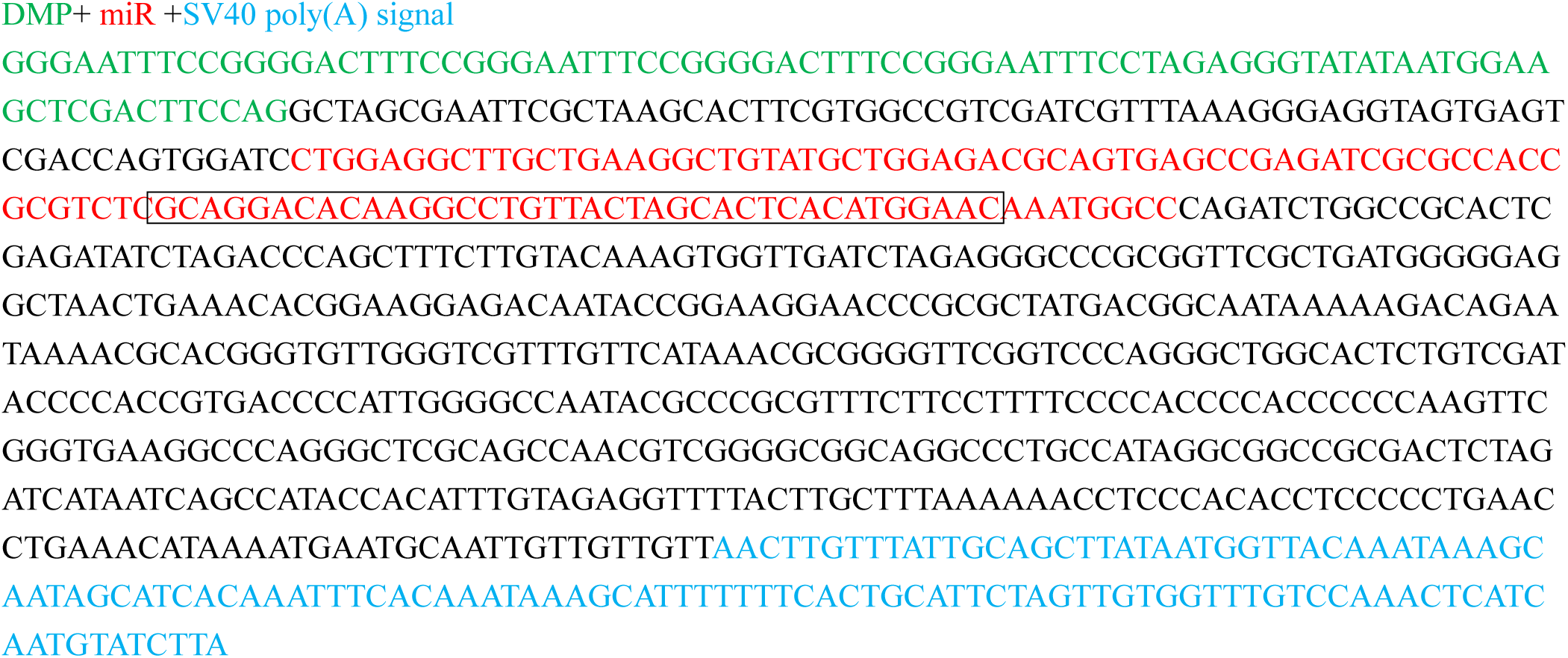

The sequence in the box in above skeleton vector can be replaced by the following sequences for constructing pDMP-miRNAs targeting particular genes:

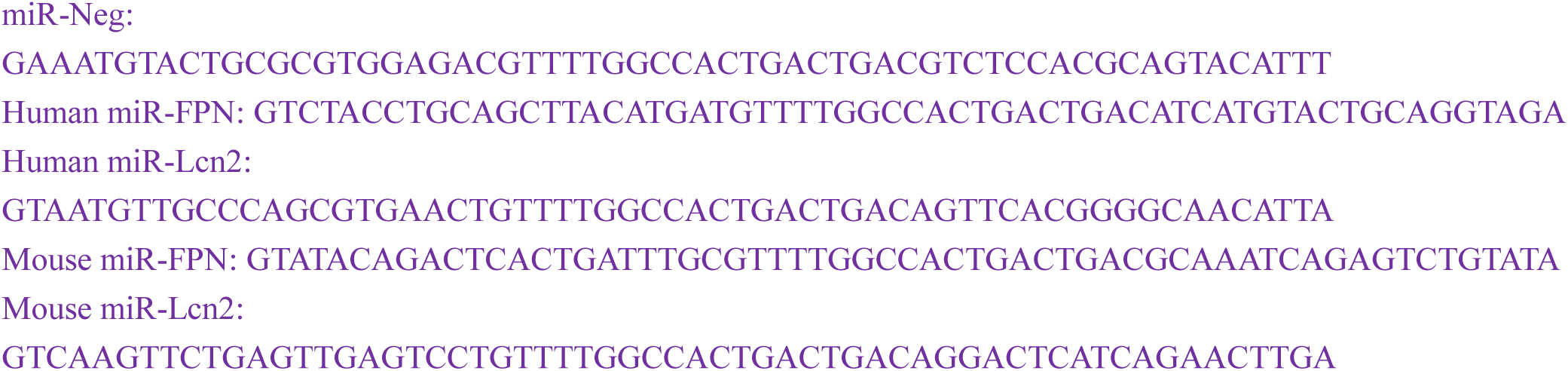

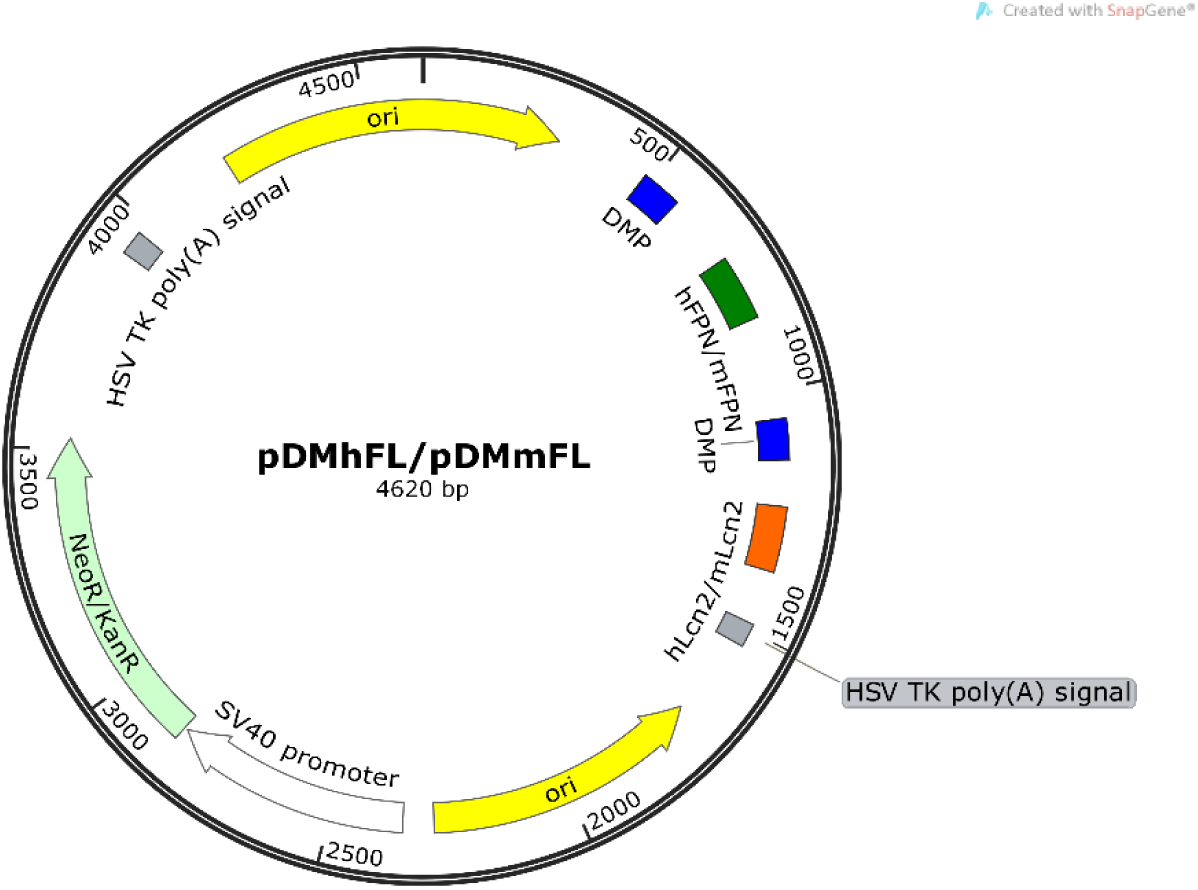

### pDMP-miR-FPN-DMP-Lcn2 (pDMhFL)

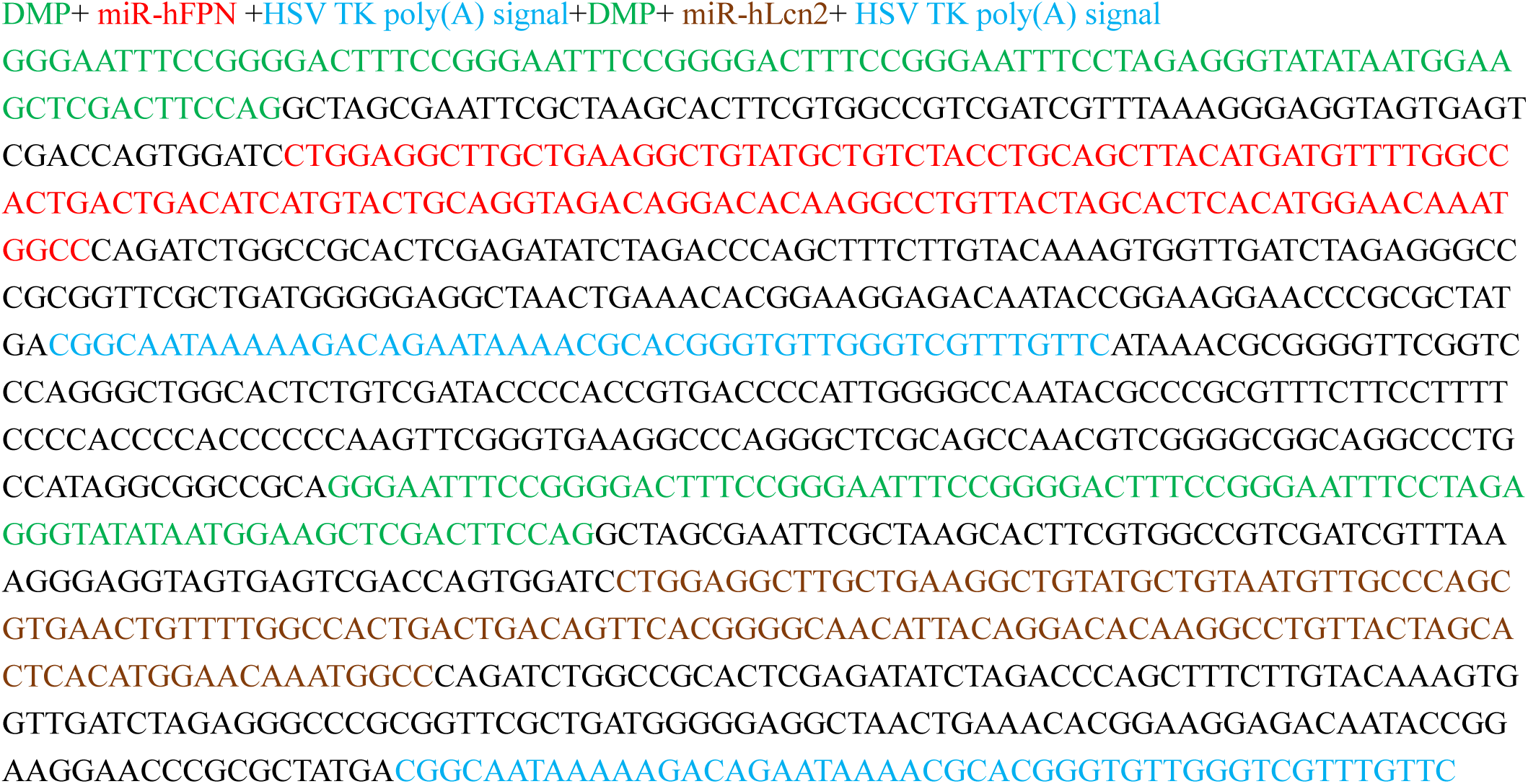

### pDMP-miR-FPN-DMP-Lcn2 (pDMmFL)

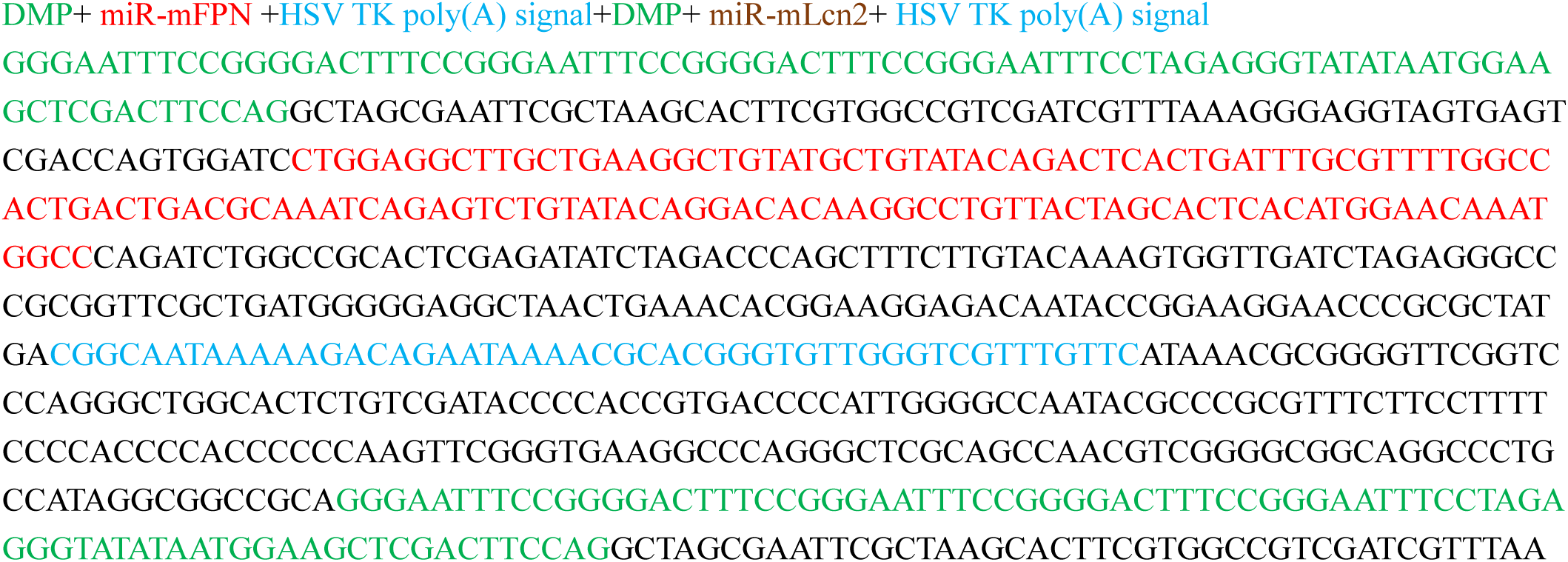

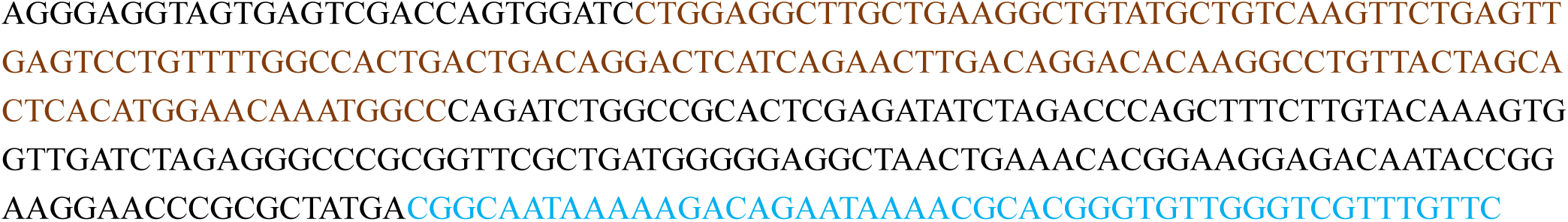

